# MS-DIAL 4: accelerating lipidomics using an MS/MS, CCS, and retention time atlas

**DOI:** 10.1101/2020.02.11.944900

**Authors:** Hiroshi Tsugawa, Kazutaka Ikeda, Mikiko Takahashi, Aya Satoh, Yoshifumi Mori, Haruki Uchino, Nobuyuki Okahashi, Yutaka Yamada, Ipputa Tada, Paolo Bonini, Yasuhiro Higashi, Yozo Okazaki, Zhiwei Zhou, Zheng-Jiang Zhu, Jeremy Koelmel, Tomas Cajka, Oliver Fiehn, Kazuki Saito, Masanori Arita, Makoto Arita

**Affiliations:** RIKEN Center for Integrative Medical Sciences, Japan; RIKEN Center for Sustainable Resource Science, Japan; Clinical Omics Unit, Kazusa DNA Research Institute, Japan; Bruker, Japan; Graduate School of Pharmaceutical Sciences, Keio University, Japan; Graduate School of Information Science and Technology, Osaka University, Japan; Department of Genetics, The Graduate University for Advanced Studies, SOKENDAI, Japan; NGAlab, Spain; Graduate School of Bioresources, Mie University, Japan; Interdisciplinary Research Center on Biology and Chemistry, Chinese Academy of Sciences, China; University of Chinese Academy of Sciences, China; Department of Pathology, University of Florida, USA; Department of Environmental Health Sciences, Yale School of Public Health, USA; Institute of Physiology of the Czech Academy of Sciences, Czech Republic; West Coast Metabolomics Center, UC Davis, USA; Graduate School of Pharmaceutical Sciences, Chiba University, Japan; National Institute of Genetics, Japan; Graduate School of Medical Life Science, Yokohama City University, Japan

## Abstract

We formulated mass spectral fragmentations of lipids across 117 lipid subclasses and included ion mobility tandem mass spectrometry (MS/MS) to provide a comprehensive lipidome atlas with retention time, collision cross section, and MS/MS information. The all-in-one solution from import of raw MS data to export of a common output format (mztab-M) was packaged in MS-DIAL 4 (http://prime.psc.riken.jp/) providing an enhanced standardized untargeted lipidomics procedure following lipidomics standards initiative (LSI) semi-quantitative definitions and shorthand notation system of lipid structures with a 1–2% estimated false discovery rate, which will contribute to harmonizing lipidomics data across laboratories to accelerate lipids research.

---

Lipids play three important roles in biology: as cellular membranes, for energy storage and use, and as signaling molecules. Biological functions are largely driven by lipid structural diversity (>10,000)^1^. MS-based untargeted lipidomics exploits highly similar fragmentation patterns within each lipid subclass species, allowing for more comprehensive annotation compared to hydrophilic metabolites, natural products, or exposome studies^2^. Despite various lipid identification software tools^3^, numerous signals with statistical significance remain unidentified in lipidomic studies. Therefore, improved and systematic lipid annotation and quantification is needed that utilizes retention time (RT), collision cross section (CCS), *m/z*, isotopic ions, adduct information, and MS/MS fragmentations. Such standardization of lipidomics results increases confidence in lipid annotations, quantifications and lipid nomenclature. Moreover, an automated data processing workflow enables the scientific community to retrieve history logs and to accelerate transparent data science in lipidomics analyses^5^.

Here, we developed a novel untargeted lipidomics platform, packaged in MS-DIAL 4 (http://prime.psc.riken.jp/) and compliant with LSI, by processing 1,056 lipidomics samples across 81 studies acquired on 10 independent liquid chromatography (LC)-MS platforms. Samples analyzed included human plasma, 19 mouse tissues, 4 mammalian cultured cell types, 9 algal species, and 3 plant species to increase coverage of lipid subclasses unique to various organisms (Fig. 1**, Supplementary Table 1, and Supplementary Note 1**). Data mining was conducted iteratively. First, lipids were quantified according to LSI guidelines (level 2 and 3 in https://lipidomics-standards-initiative.org/) using the MS-DIAL “bootstrap” version^6^ (see **Supplementary Methods**). Next, unknown MS/MS spectra were elucidated by analyzing authentic standards, mining literature, or predicting the putative structure from fragment ion evidence. Upon formulating mass fragmentation for the representative lipid structure in both ESI(+)-MS/MS and ESI(−)-MS/MS spectra, we expanded the scheme to various acyl chain varieties, referencing the heuristic MS/MS spectra in MSP format to filter noisy spectra via a classical spectral similarity calculation^7^. An example of this process is shown for *N*-acyl glycylserine, which is unique to MS-DIAL libraries (Fig. 1). After confirming the scalability of lipid subclass-associated characteristic product ions and neutral losses across various acyl chain species, a decision tree algorithm yielded an appropriate lipid structure annotation based on the MS/MS spectrum^8^ (see additional details in **Supplementary Methods** and **Supplementary Fig. 1**). Finally, MS/MS spectral libraries and decision trees for 177 ionized forms of 117 lipid subclasses were integrated into MS-DIAL 4. The classifications followed the LipidMAPS^9^ definition and the structures are represented by a shorthand notation system^10^ (**Supplementary Figs. 2–7, Supplementary Table 2, and Supplementary Note 2**): the specifications for the characters virgule “/”, underline “_”, semicolon “;”, rings/double bond equivalents, and atom strings such as “O-”, “N-”, and “P-” are fully detailed in **Supplementary Note 2**. Notably, these lipids were characterized in biological samples and formulated based on experimental data rather than *in silico*, with the coverage outperforming that of existing lipidomics software programs (**Table 1**, **Supplementary Table 3, and Supplementary Methods**): MS-DIAL 4 extended the number of lipid subclasses in the database to yield 3- and 1.5-fold coverage compared to that in the previous versions of MS-DIAL and other software programs, respectively. Moreover, MS-DIAL 4 access to decision tree annotation lacking in the prior versions provides appropriate structure representation of 117 lipid subclasses through fragment evidence for species-, molecular species-, and *sn*-position level annotations to unequivocally translate lipidomics data into biology for advancing biomarker and drug development and clinical application. Overall, we profiled 8,051 unique lipids from 117 lipid subclasses, with 6,570 characterized at the molecular species level including confirmed acyl chain-specific fragments (**Supplementary Table 4**). All results including MS-DIAL source codes, mass spectral libraries, and semi-quantitative values defined as LSI level 2 or 3 are managed in our RIKEN PRIMe website (http://prime.psc.riken.jp/) (**Supplementary Data 1**), and all MS raw data is available at the DropMet section via the indices DM0022, DM0030, and DM0031.

**Fig. 1.**
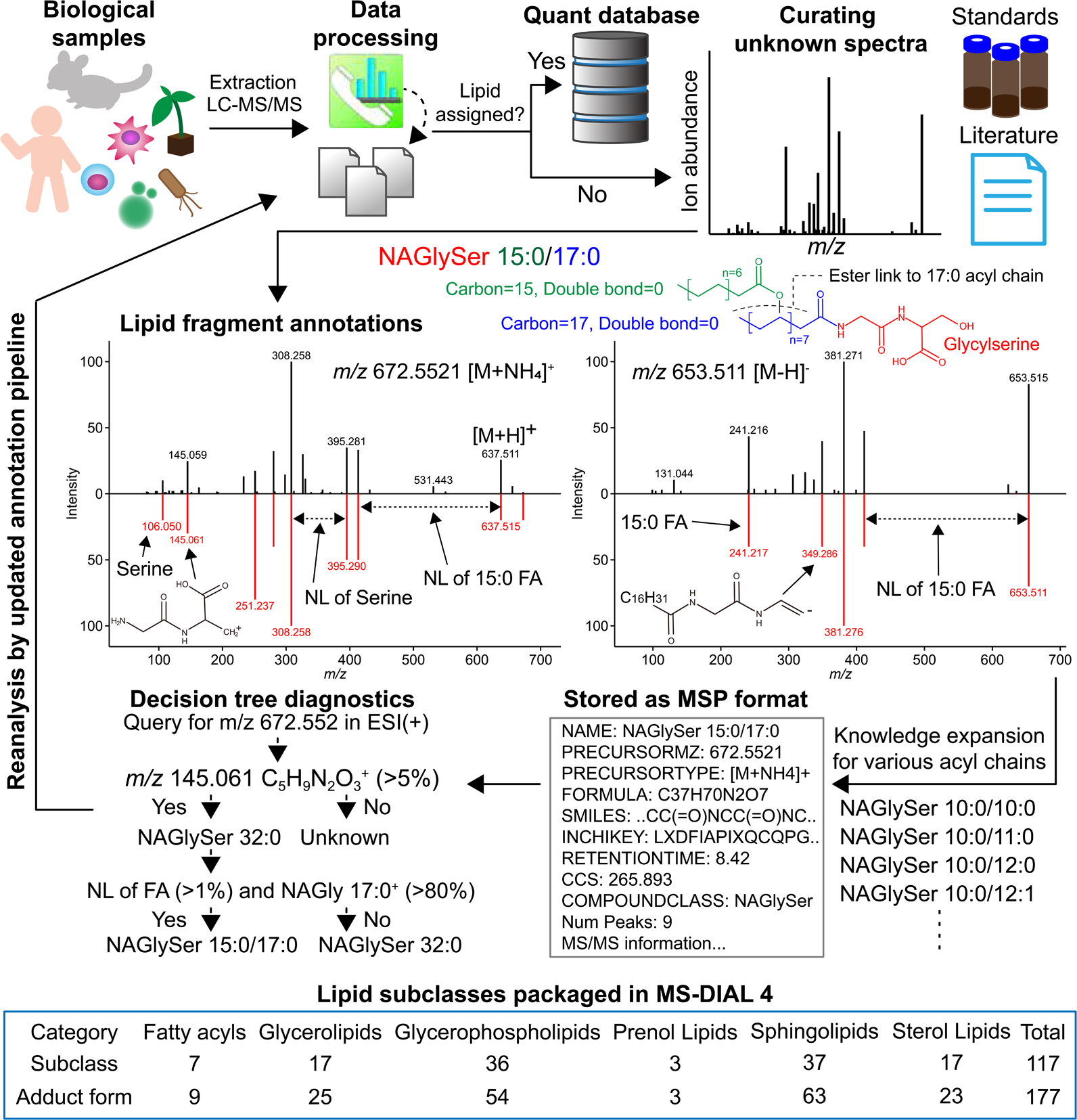
Overview of MS-DIAL 4 development. Data for 1,056 datasets acquired using liquid chromatography-tandem mass spectrometry (LC-MS/MS) and LC-ion mobility (IM)-MS/MS for human, mouse, algae, and plant biological samples were processed to determine lipid subclass fragmentation patterns. The characterized lipids were stored in our lipidomics quant database; unknowns were elucidated after excluding potential different ionized forms of known metabolites, e.g. in source fragments and different adduct types. Following the elucidation of the potential structure by curating internal standard data, literature, and lipid-specific fragment ions, subclass specific mass fragmentation rules were formulated for both positive- and negative ion MS/MS spectra, and heuristic MSP spectral libraries were generated for various acyl chains by propagating the fragmentation knowledge. The decision tree algorithm to evaluate the existence of lipid subclass-associated fragment ions and acyl chains was coded to represent the lipid structure at the species (NAGlySer 32:0) or molecular species level (NAGlySer 15:0/17:0). Iteratively, the same data was reprocessed by the updated lipidomics workflow. Finally, the platform for characterizing 177 ionized forms of 117 lipid subclasses and handling any type of LC-(IM)-MS/MS data was released as the MS-DIAL 4 software.

**Fig. 2.**
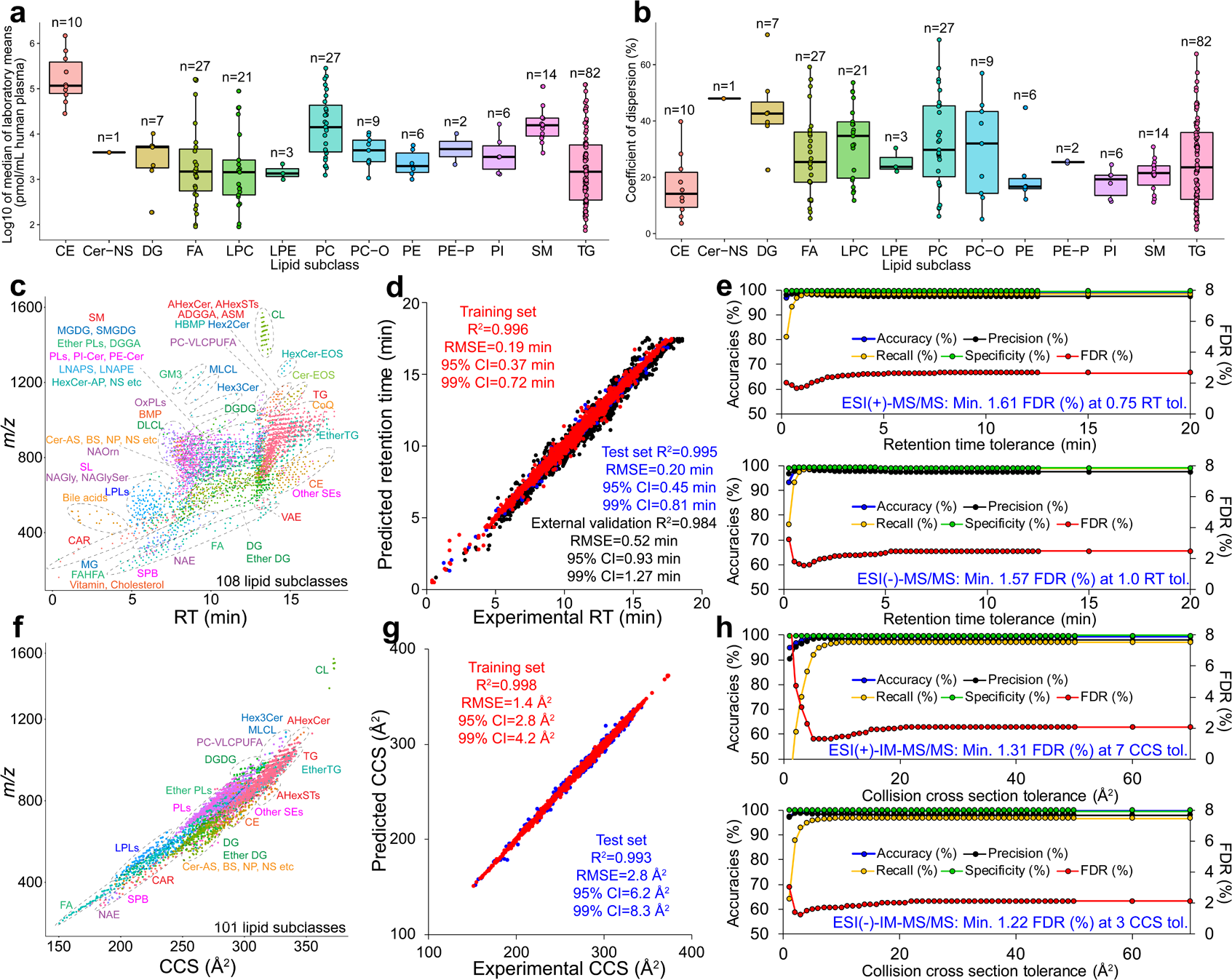
Evaluations for quantification, annotation, and scalability in MS-DIAL 4. **a**, The concentrations of 215 lipid molecules quantified by the internal standards having the same polar head moiety were separately described in each lipid subclass (CE, cholesteryl ester; Cer-NS, ceramide containing sphingosine and normal fatty acid; DG, diacylglycerol; FA, free fatty acid; LPC, lysophosphatidylcholine; LPE, lysophosphatidylethanolamine; PC, phosphatidylcholine; PC-O, alkylacyl PC; PE, phosphatidylethanolamine; PE-P, plasmenyl PE; PI, phosphatidylinositol; SM, sphingomyelin; TG, triacylglycerol). Lipid concentrations in human plasma were obtained across five or more platforms. The concentration was represented by the log 10 transformed value of the median value of platform means. **b**, The coefficient of dispersion values were calculated to evaluate the lipid concentration consensus among platforms. **c**, Retention time (RT) and *m/z* values of 4,303 lipids of 108 lipid subclasses resolved by the molecular species level, including positive- and negative ion mode results. **d**, Experimental- and predicted RT values by our machine learning method for training-, test-, and external validation sets. R-square, root mean squared error (RMSE), 95% confidence interval (95% CI), and 99% CI are shown. **e**, Using the set of true positives and true negatives for liquid chromatography-tandem mass spectrometry (LC-MS/MS) lipidomics data, the accuracy, precision, recall, specificity, and false discovery rate (FDR) of our annotation workflow were evaluated with various RT tolerance settings for positive (top)- and negative (bottom) ion modes. **f**, Collision cross section (CCS) and *m/z* values of 3,570 molecules of 101 lipid subclasses resolved by molecular species level, including the positive- and negative ion mode results. **g**, Experimental- and predicted CCS values by our machine learning method for training and test sets. R-square, RMSE, 95% CI, and 99% CI are shown. **h**, Using the set of true positives and true negatives for LC-IM-MS/MS lipidomics data, the accuracies and FDR as in (**e**) were evaluated for various CCS tolerances with 1.5 min RT tolerance.

**Table 1.** Comparison of MS-DIAL 4 with other lipidomics software tools.

| Software name | Peak detection | Peak alignment | Ion mobility data support | MS/MS similarity calculation | Decision tree annotation* | Supported lipid subclasses** |
| --- | --- | --- | --- | --- | --- | --- |
| MS-DIAL 4 | Yes | Yes | Yes | Yes | Yes | 143 (117) |
| MS-DIAL 3 | Yes | Yes | No | Yes | No | 52 (52) |
| Lipidex | No | No | No | Yes | No | 62 (39) |
| LipidMatch-2.0.2 | No | No | No | No | Yes | 90 (58) |
| LIQUID_v7.4.6988 | Yes | No | No | Yes | No | 96 (31) |
| LipidHunter2 | Yes | No | No | Yes | No | 14 (14) |
| Greazy | No | No | No | Yes | No | 35 (23) |
| MZmine2 2.51 | Yes | Yes | No | Yes | No | 31 (15) |

The evaluated tool name, version, and the function/utility existence for peak detection/picking, peak alignment/data integration, and tandem mass spectrometry (MS/MS) annotation method are listed, along with the count of annotatable lipid subclasses by the default library.
\*Provides appropriate structure representation of lipids by fragment evidence for species level, molecular species level, *sn*-position level, and full structure level annotations. \*\*Count of annotatable lipid subclasses (count of lipid subclasses characterized in this study). For other tools, the annotatable counts were determined by checking supported lipid subclasses in the respective databases.

MS-DIAL 4 was validated using three LC-MS study subsets (Fig. 2). First, we processed NIST human plasma (SRM 1950) lipidomics data acquired on eight independent platforms with different extraction methods, internal standards, and instruments including machines of Waters, SCIEX, Thermo Fisher, Bruker, and Agilent Technologies. We annotated 961 unique lipids in total at the molecular species level from the data of eight platforms, with the maximum, minimum, and average of annotated lipid counts among the platforms being 495, 118, and 318, respectively. This result indicated that lipid extraction procedures and MS machines substantively influence the annotations even with an identical data processing platform. Consensus values were estimated by the median of platform means and the sample coefficient of dispersion (COD) values as previously proposed for interlaboratory comparison^11^, considering 215 lipids annotated by at least five platforms that were quantified according to LSI level 2 definition. Similar to previously reported amounts^12^, human plasma lipid concentration ranged from 100 nM to 1 mM (Fig. 2a); the semi-quantitative values were also validated using LipidQC^13^ software (**Supplementary Table 5**). Among the plasma lipid species, 81% exhibited an estimated COD <40%, considered acceptable for quality control activities (Fig. 2b). Averaged CODs for LipidMAPS categories were 27% (FAs; *n* = 27), 27% (GLs; *n* = 89), 30% (GPs; *n* = 74), 23% (SPs; *n* = 15), and 17% (STs; *n* = 10), indicating that MS-DIAL 4 automated quantification provides validated cross-laboratory lipidomics data under reverse-phase LC-MS conditions.

Second, processing 850 files of 61 studies acquired in the same LC conditions resolved 4,303 lipids of 108 subclasses at the molecular species level (Fig. 2c and **Supplementary Table 6**). After tuning the RT prediction model providing <1.0 min error for 95% confidence interval (CI) for an external validation set (Fig. 2d), we estimated the false discovery rate (FDR) using the validation sets containing true positives of 12,798 and 10,600 and true negatives of 64,121 and 30,290 for positive and negative mode data, respectively (**Supplementary Methods**, Fig. 2e and **Supplementary Data 2**). Critically, true negatives denote uncharacterized MS/MS spectra in MS-DIAL 4, whereas true positives were created by MS specialists curating all annotatable MS/MS spectra at the molecular species level in 850 analysis files. The evaluation set excluded lipid subclasses requiring distinct internal standard RTs such as free fatty acid (FA), bile acids, and *N*-acyl ethanolamines. The minimum estimated FDR values were 1.61 and 1.57% in positive and negative sets at 0.75- and 1.00-min RT tolerances, respectively, maintaining an FDR <5% even without RT information. Our annotation was further evaluated using parallel accumulation-serial fragmentation (PASEF) data available at Bruker timsTOF Pro machine, providing four lipid physicochemical properties, i.e. RT, *m/z*, CCS, and MS/MS via LC-ion mobility data-dependent-MS/MS acquisition (LC-IM-DDA) (Fig. 2f**, Supplementary Figs. 8 and 9**)^14^. Notably, MS-DIAL 4 is the first vendor-free metabolomics and lipidomics software able to handle Bruker IM-DDA and Agilent- and Waters IM data-independent acquisition (DIA) with the support of IM chromatogram deconvolution. As IM-MS facilitates metabolite characterizations with the CCS information that constitutes an orthogonal physicochemical property from RT and *m/z* independent from mass or lipophilicity, the IM data support provides a more confident data processing platform for autonomous metabolomics and lipidomics. Using PASEF-based lipidomics data, we established a comprehensive CCS lipid library using a machine learning technique with 95% CI = 6.2 Å^2^ (Fig. 2g and **Supplementary Table 7**). We estimated FDR using the sets of true positives of 2,598 and 1,670 and true negatives of 30,131 and 20,737 in positive and negative ion mode data, respectively (Fig. 2h), yielding estimated FDRs of 2.82 and 2.22% at 5 and 2 Å^2^ tolerances, respectively, albeit 1.31 and 1.22% when including predicted RT information (**Supplementary Fig. 10**). The ultimate atlas of RT, CCS, and MS/MS for 600,737 lipid structures is packaged in MS-DIAL 4. Particularly, the CCS values were predicted using a training set of 3,570 lipid ions from 101 subclasses, outperforming previous CCS prediction studies for lipids^15^. MS-DIAL 4 now provides the most comprehensive resource containing the mass fragmentations for lipid subclasses and the first all-in-one solution for ion mobility centric metabolomics and lipidomics research.

Finally, our platform was showcased by illuminating common and specific mammalian tissue lipids to examine actual molecular structural diversity, evaluating 112 general and six special lipid subclasses containing very-long-chain polyunsaturated FAs^16^ (VLC-PUFAs) (**Supplementary Fig. 11**). Using hierarchical clustering analysis of species counts per lipid subclass, mouse feces, cultured cell, and mouse brain tissue lipidomes clearly segregated from those of other tissues, whereas human/mouse plasma, small/large intestine, muscle tissues (skeletal muscle and heart), and metabolic- and immune organs were clustered. In addition to the common lipid subclasses that were detected in most tissues and cells and also characterized in other software programs, our platform also revealed tissue-specific lipids. For example, numerous lipid subclasses were found to be unique to fecal matter, reflecting microbiome-specific lipids including e.g. *N*-acyl amides of FA ester of hydroxy FA (FAHFA) that interact with G-protein-coupled receptors^17^. Moreover, we annotated ether phosphatidylglycerol (ether PG), digalatosyl diacylglycerol (DGDG), its alkylacyl type, sulfonolipid (SL), phosphoinositol-ceramide (PI-Cer), acylsterol glycosides (AHexST), and diacylglyceryl glucuronide containing an additional *O*-acyl chain on the uronosyl moiety (ADGGA) in mouse feces, of unknown biological importance. Mouse brain tissue contained a hexosyl ceramide (AHexCer), containing an *O*-acyl hexosyl moiety, whose structure was putatively proposed by indirect evidence of fragment ion and RT behavior compared to the acyl chain positional isomer (HexCer-EOS) with the additional acyl chain in the omega-*O*-acyl backbone (**Supplementary Fig. 12**). Skin and kidney samples contained various types of ceramides including the epidermal acyl ceramide (Cer-EOS, HexCer-EOS), and abundant di- and trihexosyl ceramides (Hex2Cer and Hex3Cer), respectively. Acylated sphingomyelin incorporating an *O*-acyl chain in the hydroxy moiety of the long-chain base (ASM) was detected in the kidney tissue and 3T3-L1 cells upon adipocyte-like differentiation. The testis and brain, in part, contained seminolipid (ether SMGDG), and the eye and testis, and adrenal gland tissues exhibited phosphatidylcholine (PC), Cer-NS, SM, cholesteryl ester (CE), DG, and triacylglycerol (TG) containing VLC-PUFAs of 30:5, 30:6, 32:5, 32:6, 34:5, and 34:6 *O*-acyl or *N*-acyl chains. Thus, lipid structures may be specialized by the polar head moiety as well as the acyl chain quality, as previously proposed in mammalian ceramides^18^. Overall, our lipidomics platform encompassed 112 lipid subclasses in mammalian tissues and cells whereas five subclasses such as diacylglyceryl-3-*O*-carboxyhydroxymethylcholine (DGCC) and diacylglyceryl trimethylhomoserine (DGTS) were only annotated in our algae data. However, such algal lipids are potentially also characterizable in human biospecimens such as periodontal plaque and fecal matter^19^, enhancing our understanding of human disease-associated lipid metabolism. Our lipidomics coverage thus outperforms existing reports and encompasses mammalian lipid diversity, highlighting the obvious advantage of the untargeted lipidomics approach with our computational MS platform.

Our lipidomics platform facilitated the illumination of lipid diversity with a small false positive rate (<2% FDR) and will assist discovering new physiological roles and mechanisms of phenotype modulation, i.e., the quality of lipids in health and disease (“LipoQuality”)^20^; semi-quantitative values of lipids obtained in this study can also be investigated in the responsible database (http://lipidbank.jp/). Although several lipid subclasses including AHexCer, ADGGA, and ASM are characterized only through indirect MS data-based evidence (**Supplementary Fig. 12**), the information will facilitate annotation of unknown lipids whose structures will ultimately be confirmed by their authentic standards. Moreover, our study revealing structures of previously unknown lipids leads to next steps identifying lipid metabolisms by integrating other omics data with the characterization of lipid localizations through imaging MS, with computational MS for such an integrative analysis and databasing expected to be addressed in the future. Overall, MS-DIAL 4 enhances the standardized lipidomics procedure from import of raw LC-MS/MS and LC-IM-MS/MS data to mztab-M format export following LSI classification, nomenclature, and semi-quantitative definitions, thereby contributing to harmonizing lipidomics data across laboratories to accelerate lipids research.

## Author contributions

H.T., K.I., Mas.A. and Mak.A. designed the research. H.T. developed all computational programs, and M.T. and I.T. contributed to source coding for the lipid characterizations and IM-DIA-MS deconvolution, respectively. H.T. and M.T. mainly curated lipid mass spectra. H.T., K.I., Y.M., H.U., and Y.H. analyzed biological samples, and J.K., T.C., O.F., and K.S. provided LC-MS/MS data of NIST human plasma. H.T. and A.S. performed software comparisons, and H.T., A.S., P.B., Z.Z., and Z.-J.Z performed RT and CCS predictions for metabolites. H.T., M.T., H.U., N.O., Y.O., and J.K. performed the lipid MS/MS library creations, and Y.Y. created the lipidomics database. H.T. wrote the manuscript. H.T., K.I., Mas.A., and Mak.A. thoroughly discussed this project, and all authors helped to improve the manuscript.

## Acknowledgements

This work was mainly supported by the JSPS Grant-in-Aid for Scientific Research on Innovative Areas “LipoQuality” (15H05897, 15H05898 for Mak.A.). We thank Steve Madden and John Fjeldsted (Agilent Technologies) for sharing the MIDAC SDK and Sven Brehmer (Bruker) for sharing the tims SDK, and Waters developmental team for sharing the SDK for travelling wave ion mobility data. We thank Xiangdong Li for his help in determining library inconsistencies and errors in conjunction with his development of Lipid Annotator. We also thank Koji Takano (RIKEN) for LC-MS analysis, Nils Hoffmann and Steffen Neumann (Leibniz Institute) for the discussion of mztab-M export, and Gerhard Liebisch (University Hospital Regensburt) for the discussion of lipid nomenclature. This work was partially supported by AMED-LEAP under grant number JP18gm0010003 (for Mak.A.), RIKEN Pioneering Project “Glyco-lipidologue Initiative”, JSPS KAKENHI (18H02432, 18K19155 for H.T.), JST National Bioscience Database Center (NBDC for Mas.A.), the Czech Science Foundation (20-21114S for T.C.) and the Czech Ministry of Education, Youth and Sports (LTAUSA19124 for T.C.).

## Data and Source code availability statement

All data resources, program packages, and source codes are freely available at http://prime.psc.riken.jp/.

## Additional information

**Supplementary information** is available for this paper as Supplementary Figures 1–12, Supplementary Tables 1–7, Supplementary Methods containing Supplementary Notes 1–2, and Supplementary Data 1–2.

---

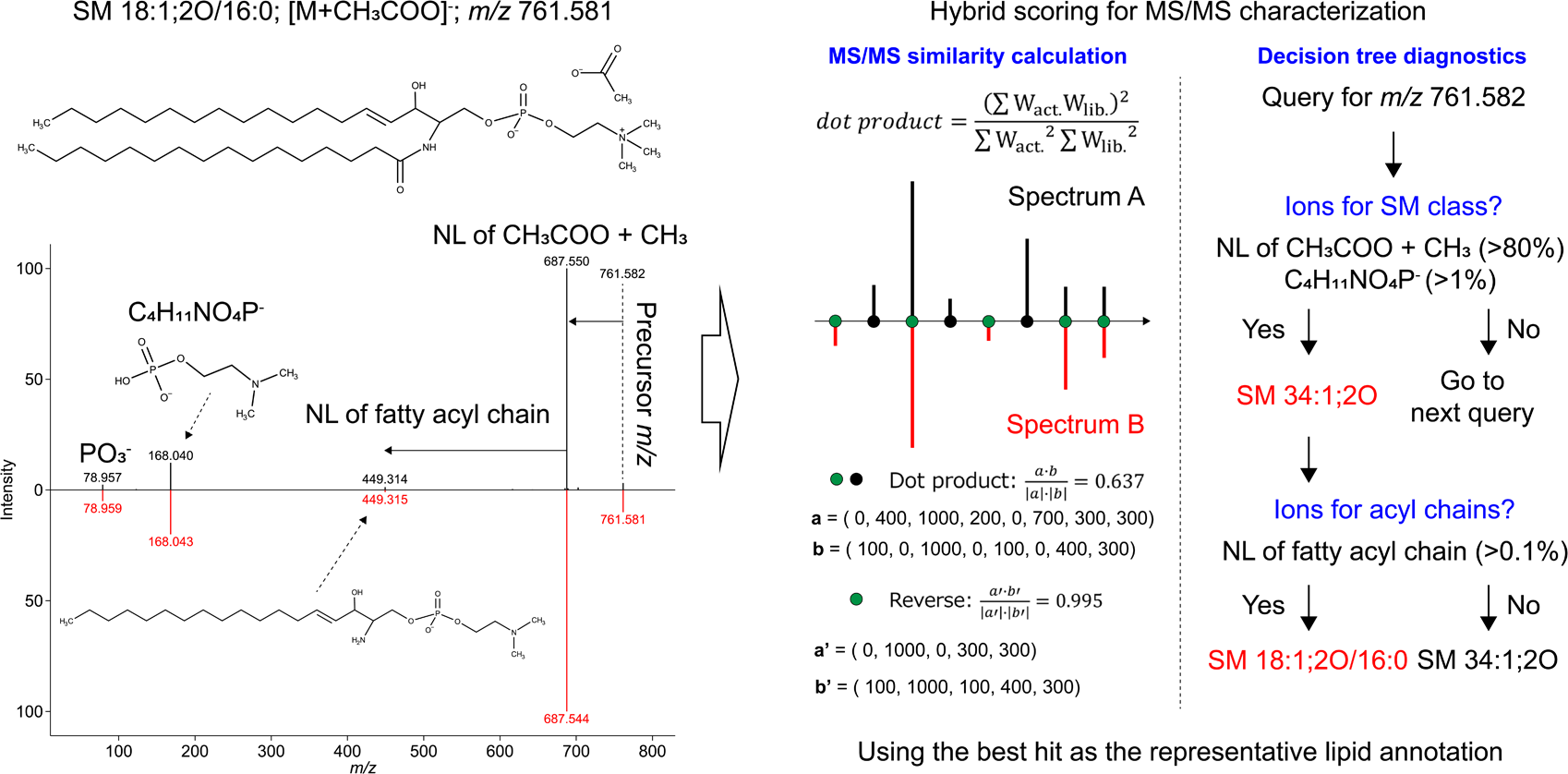

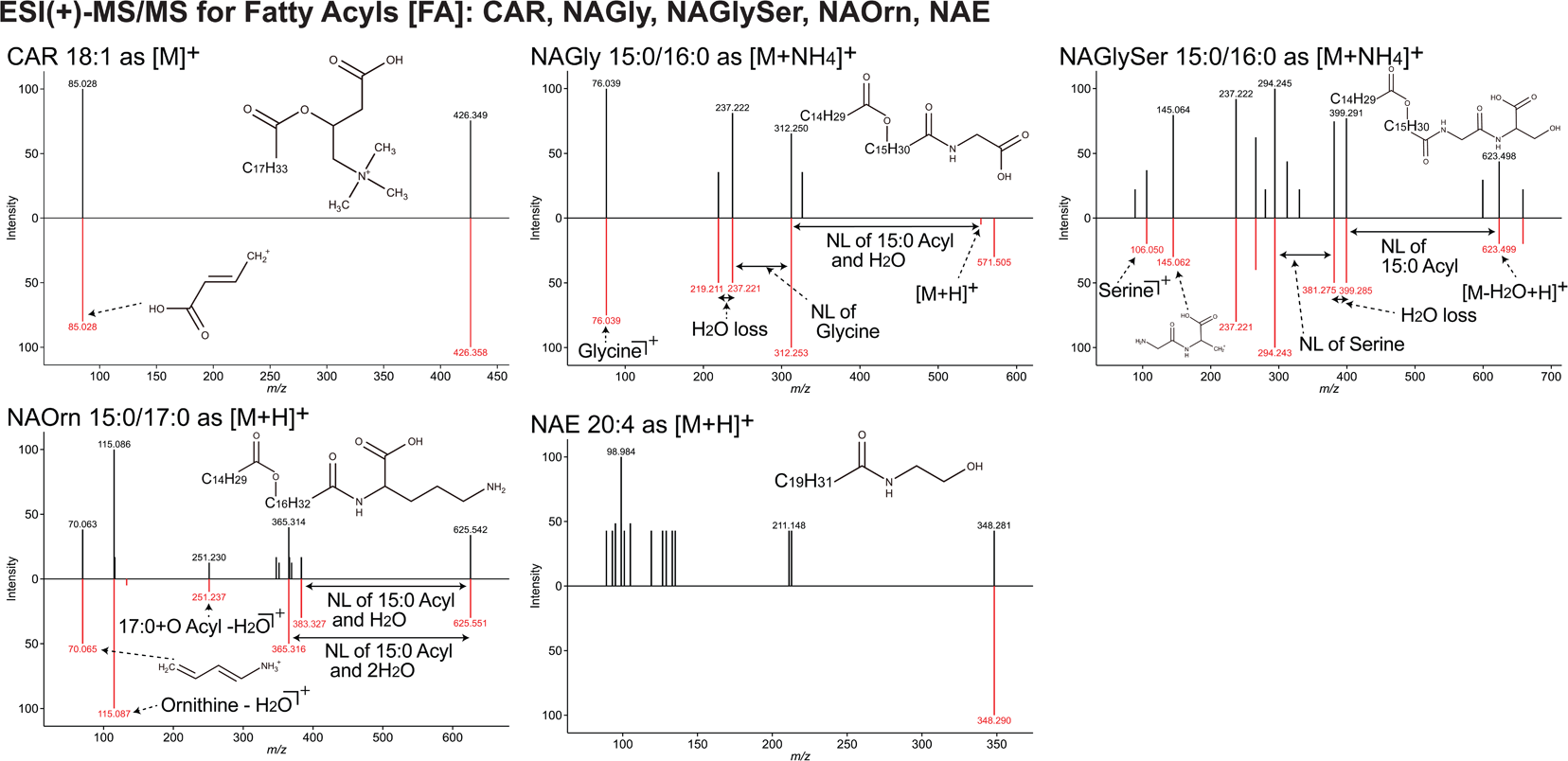

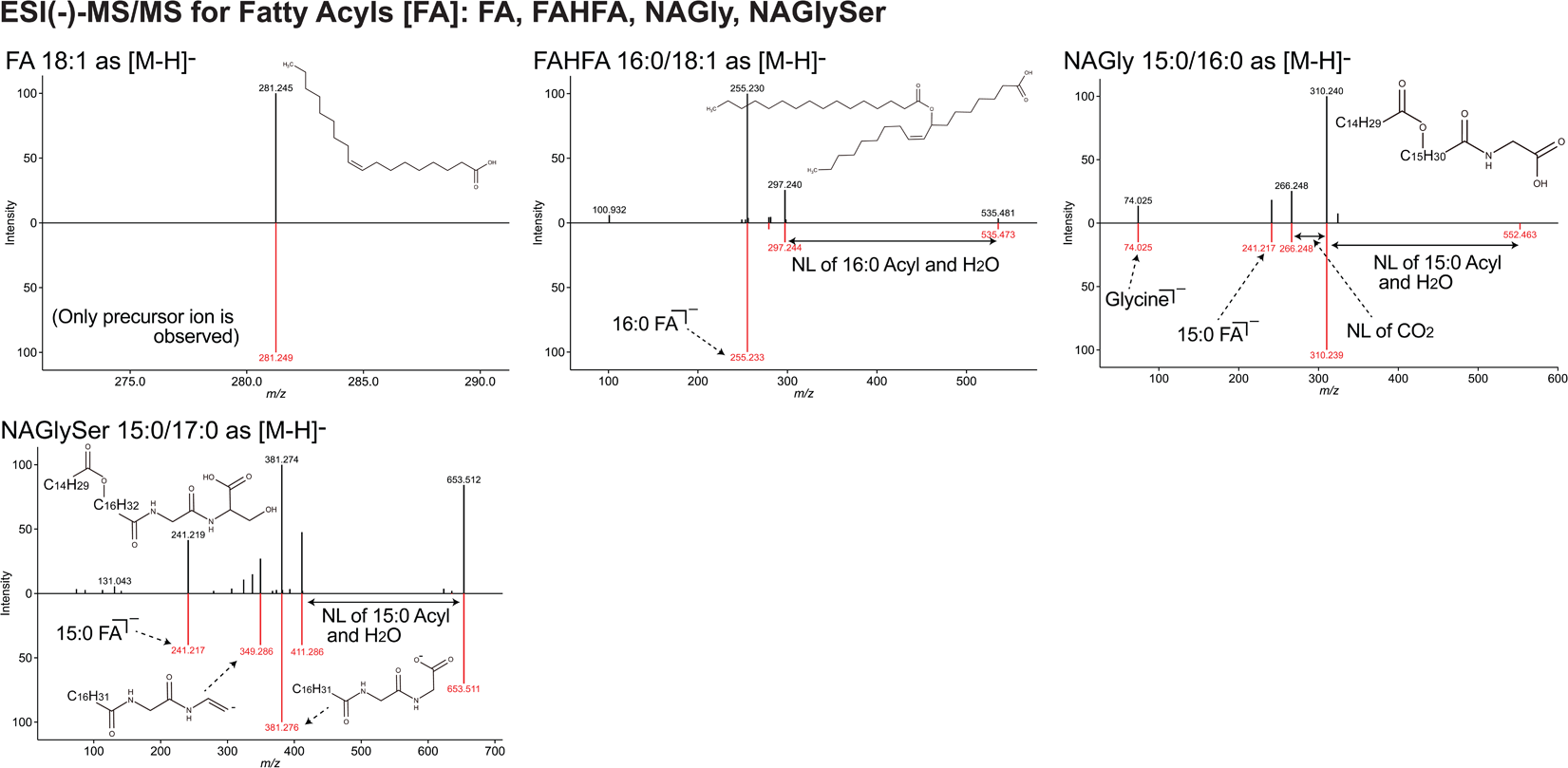

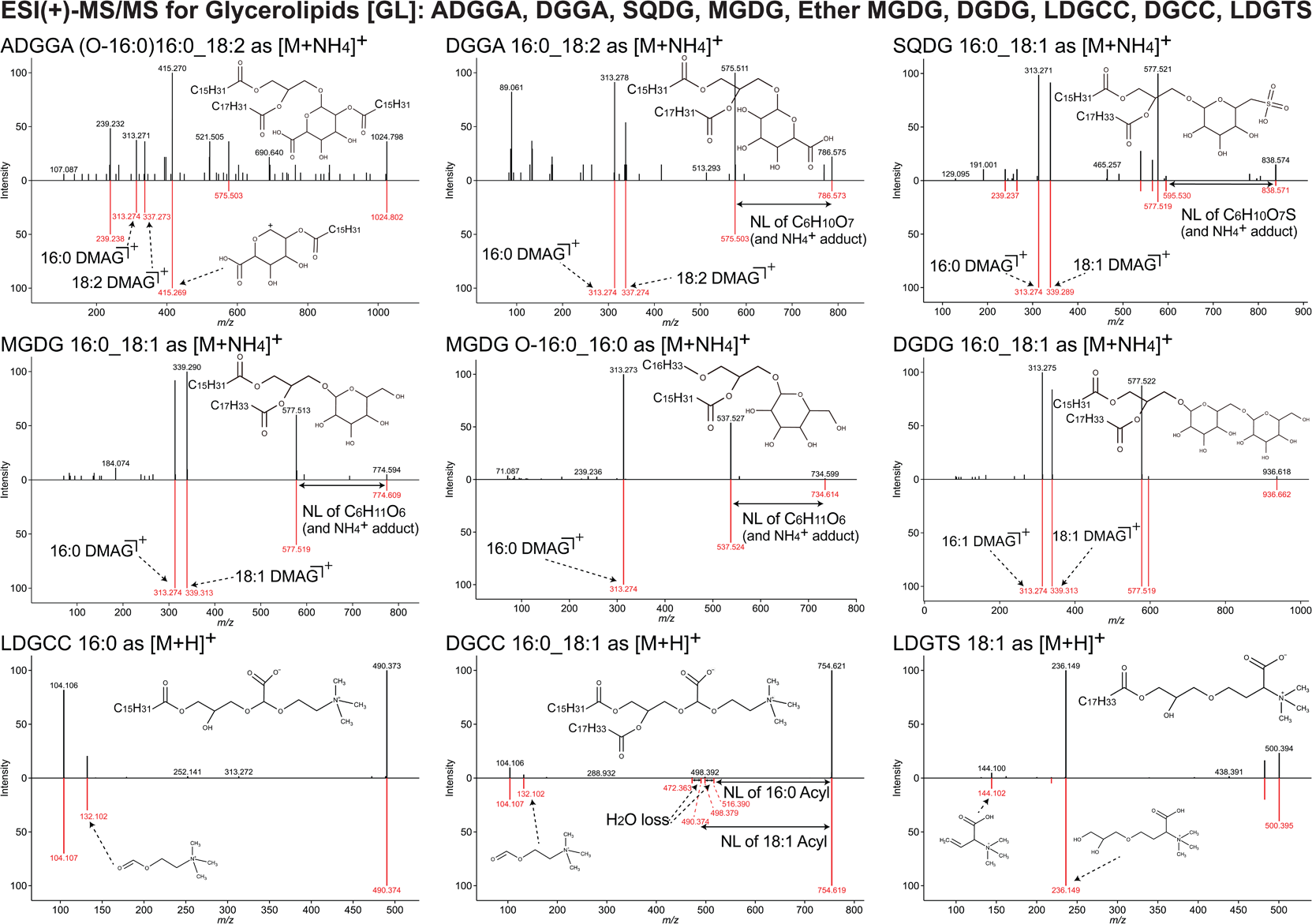

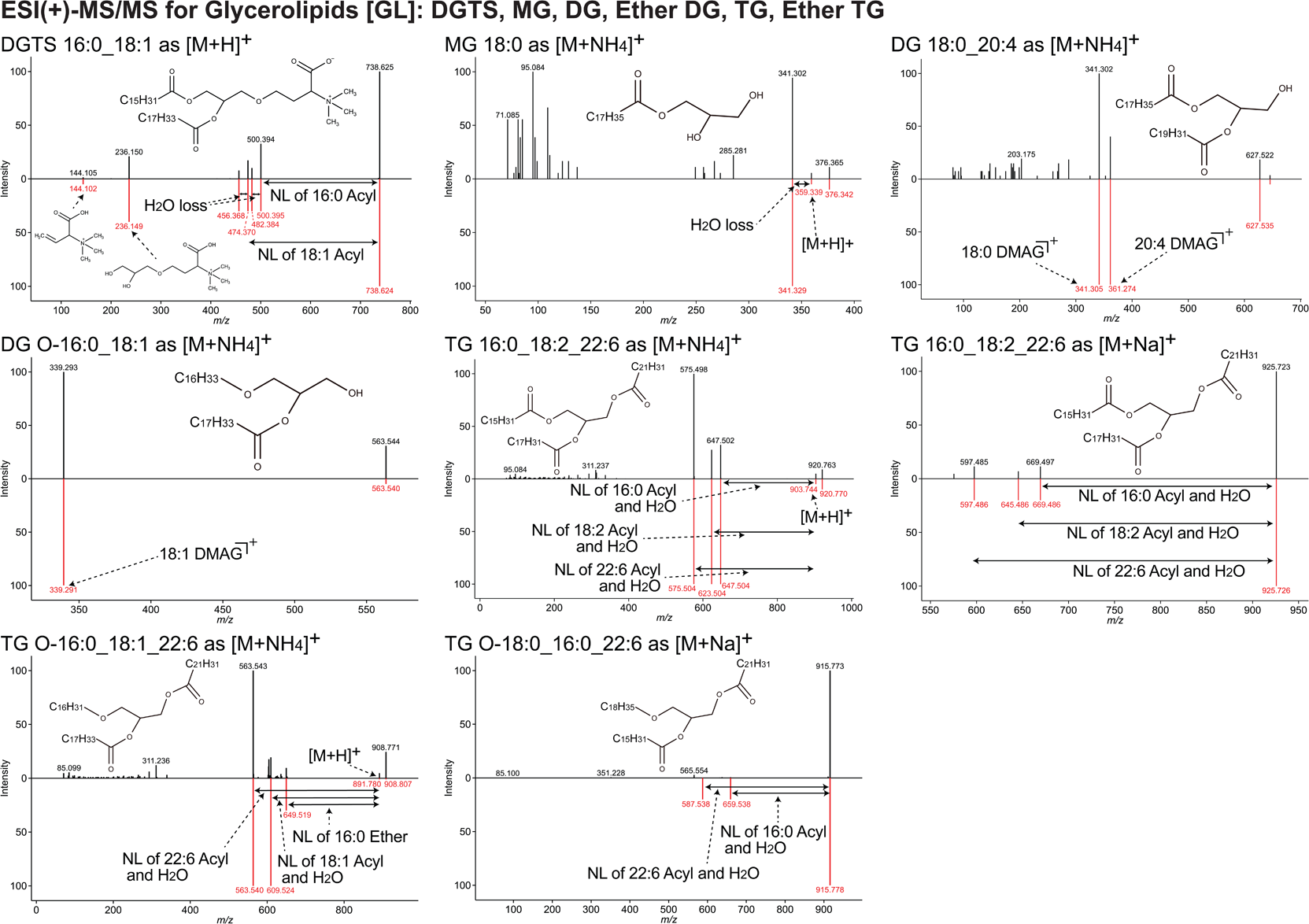

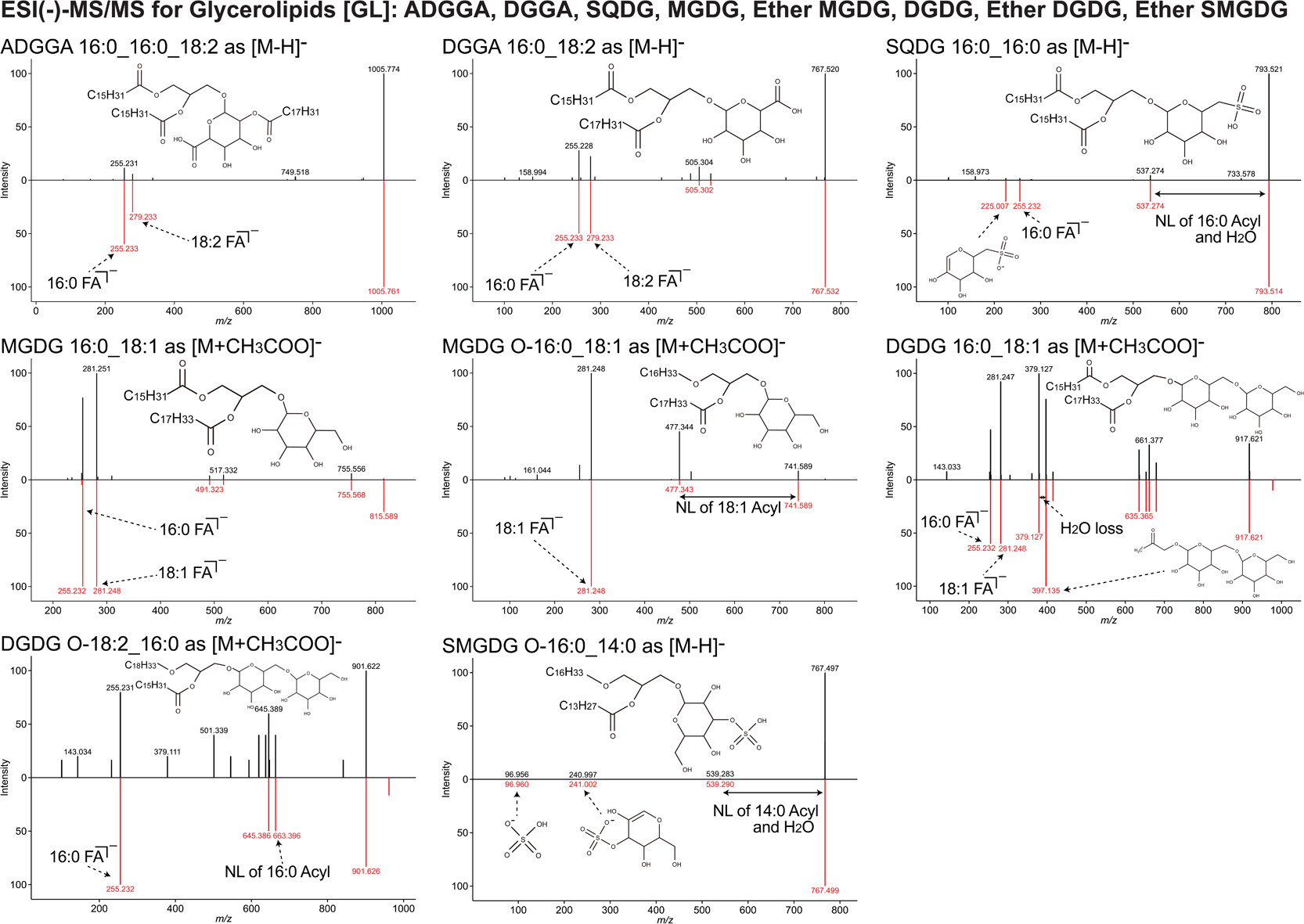

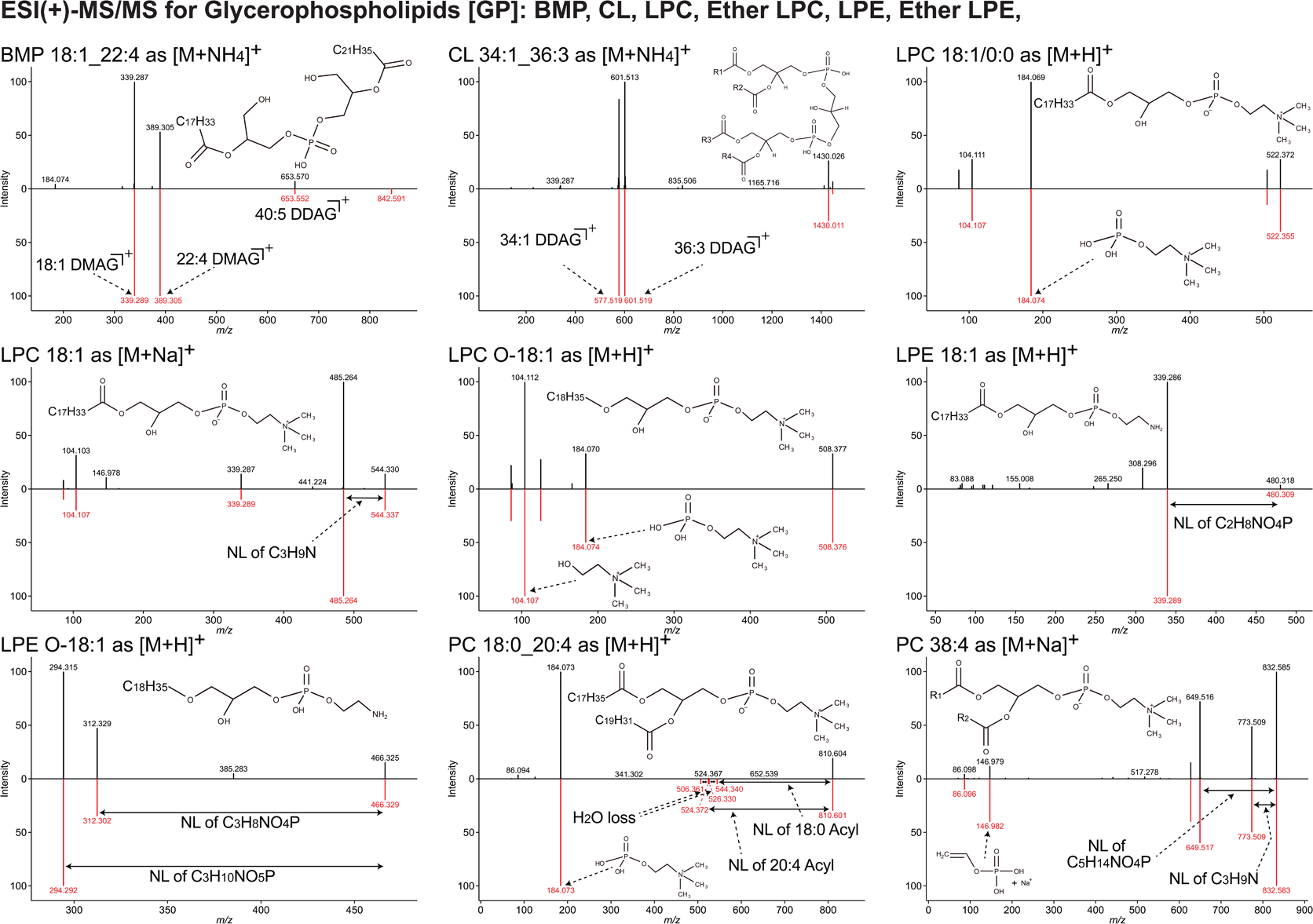

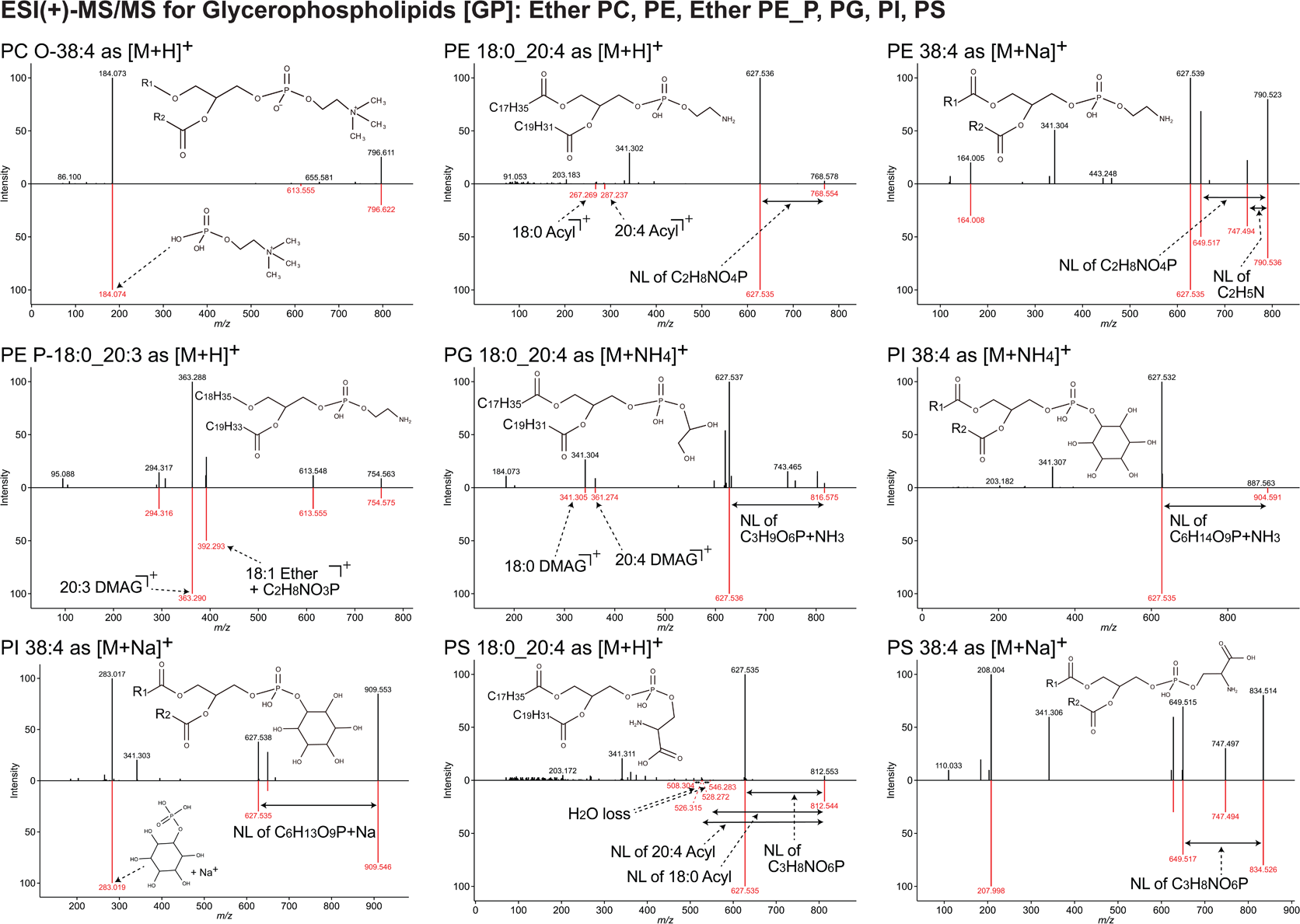

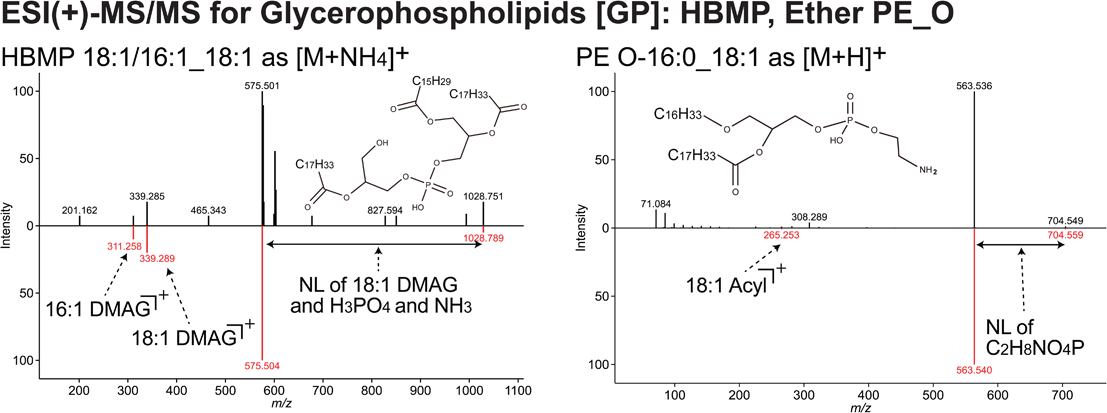

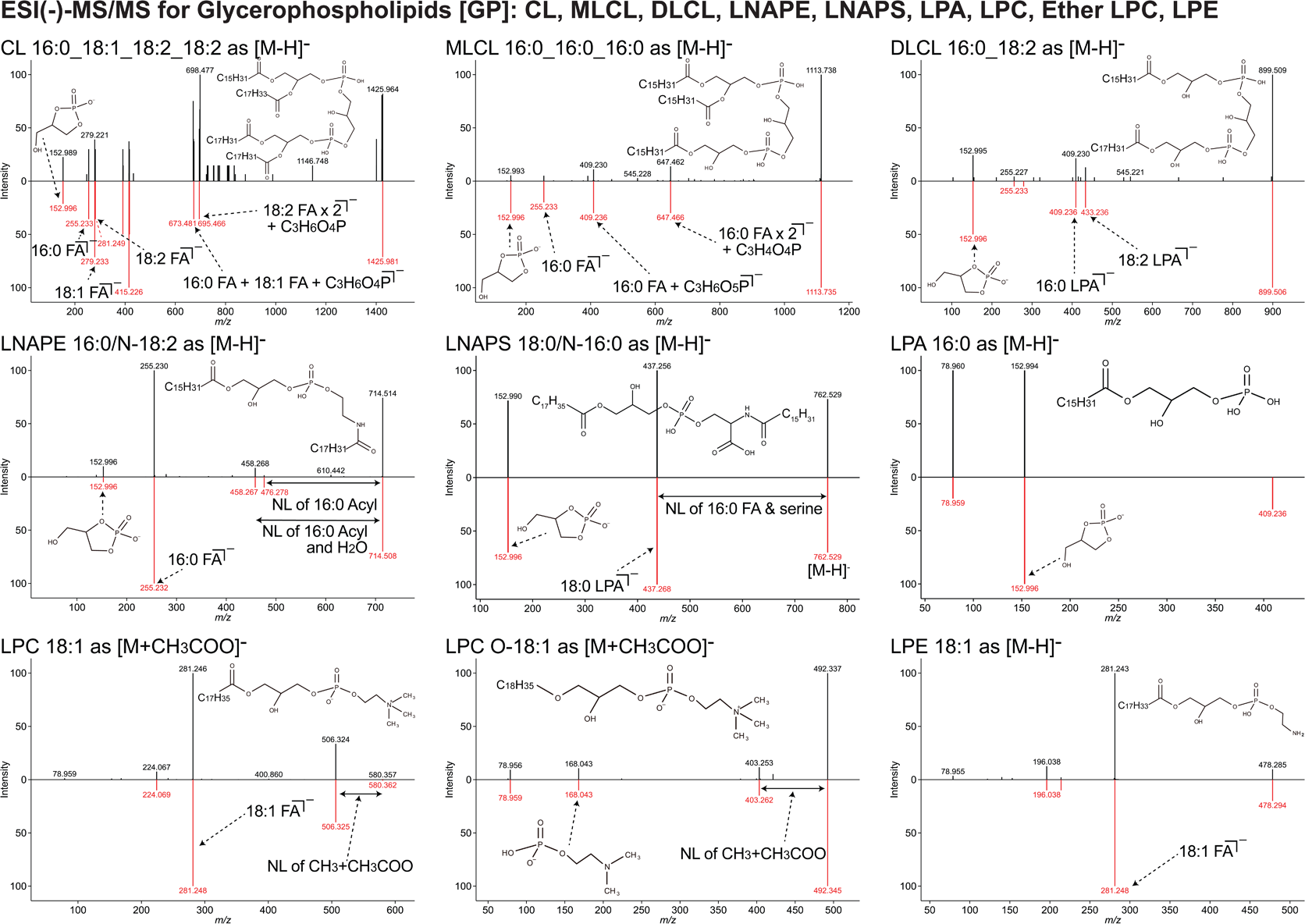

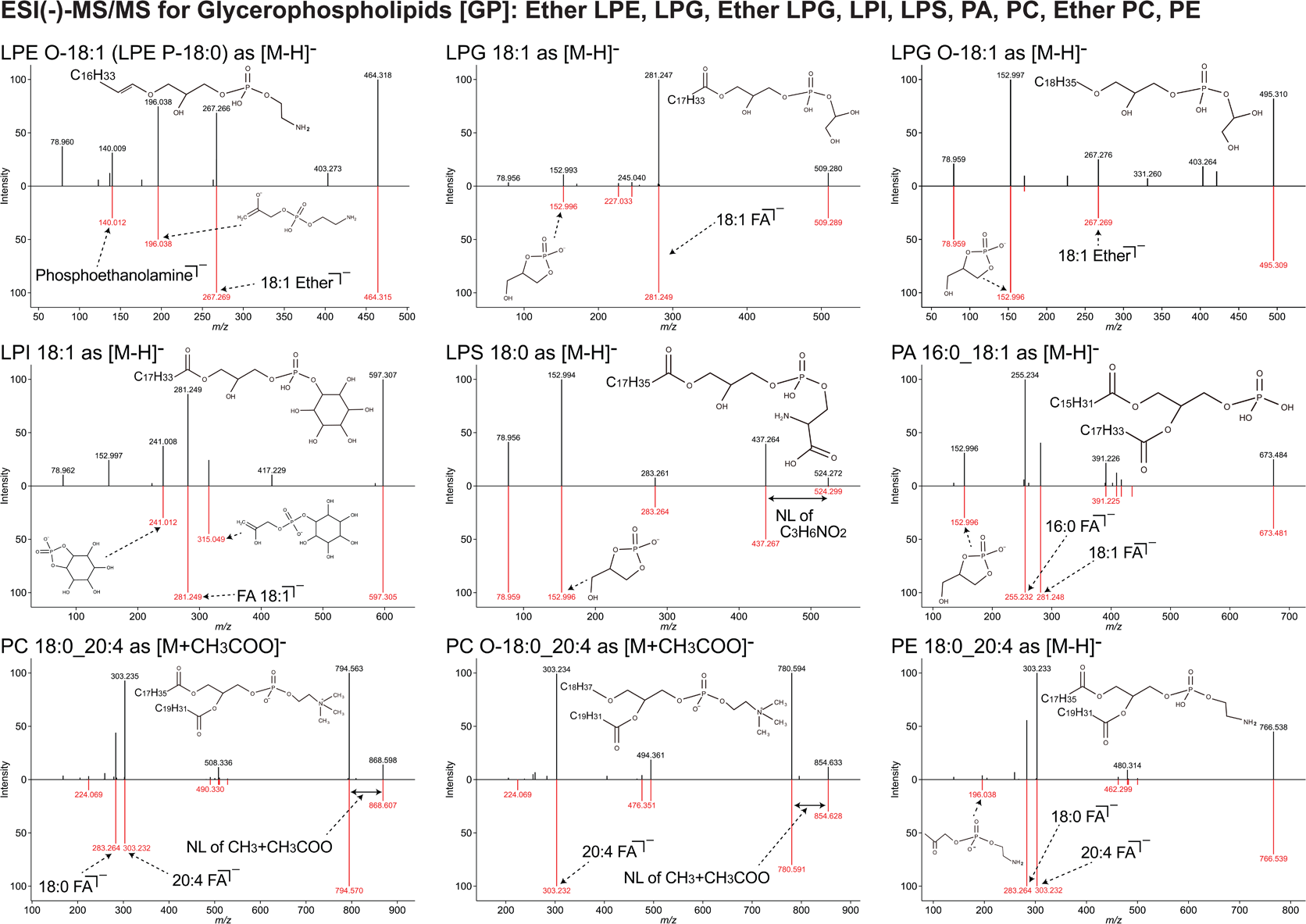

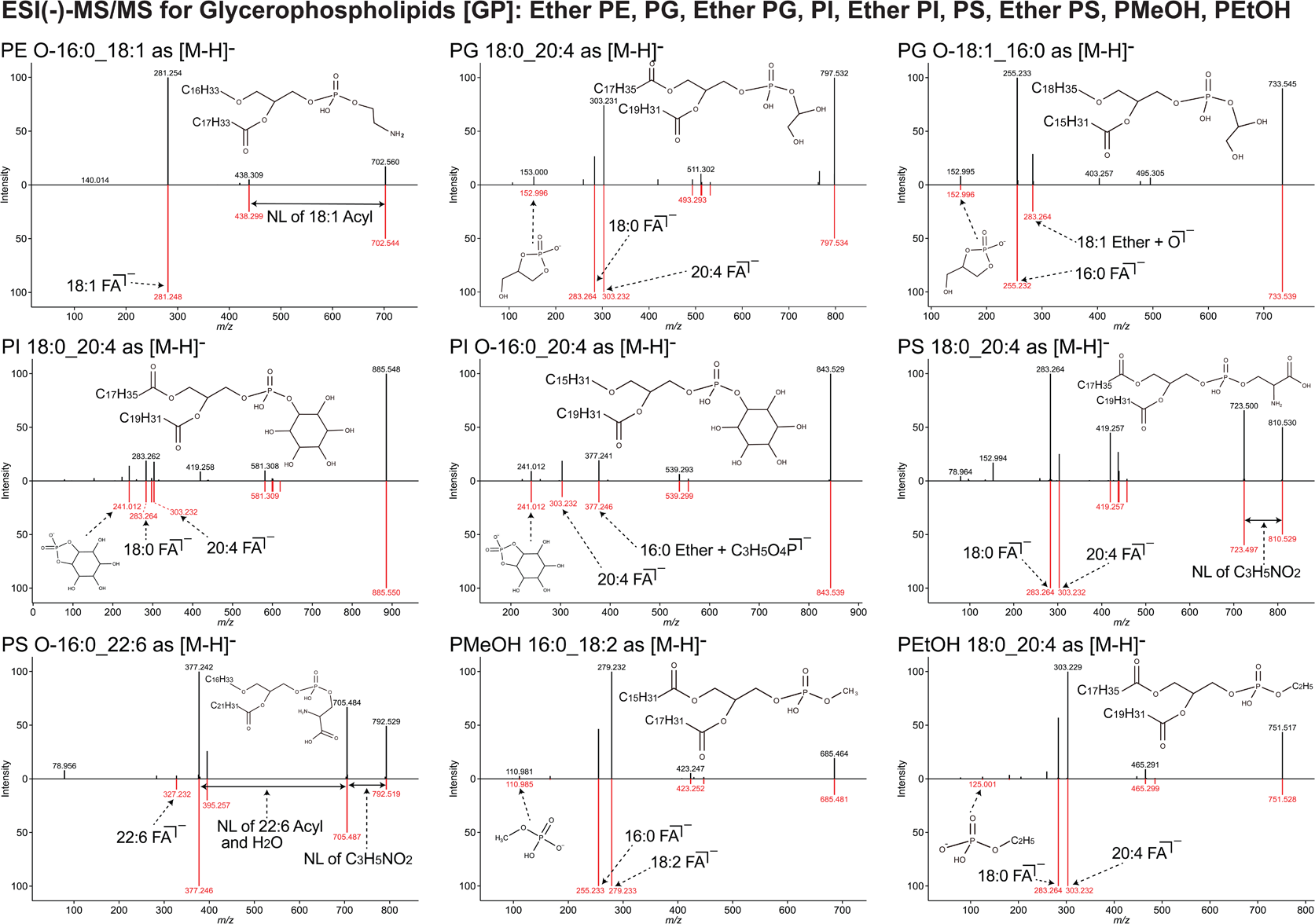

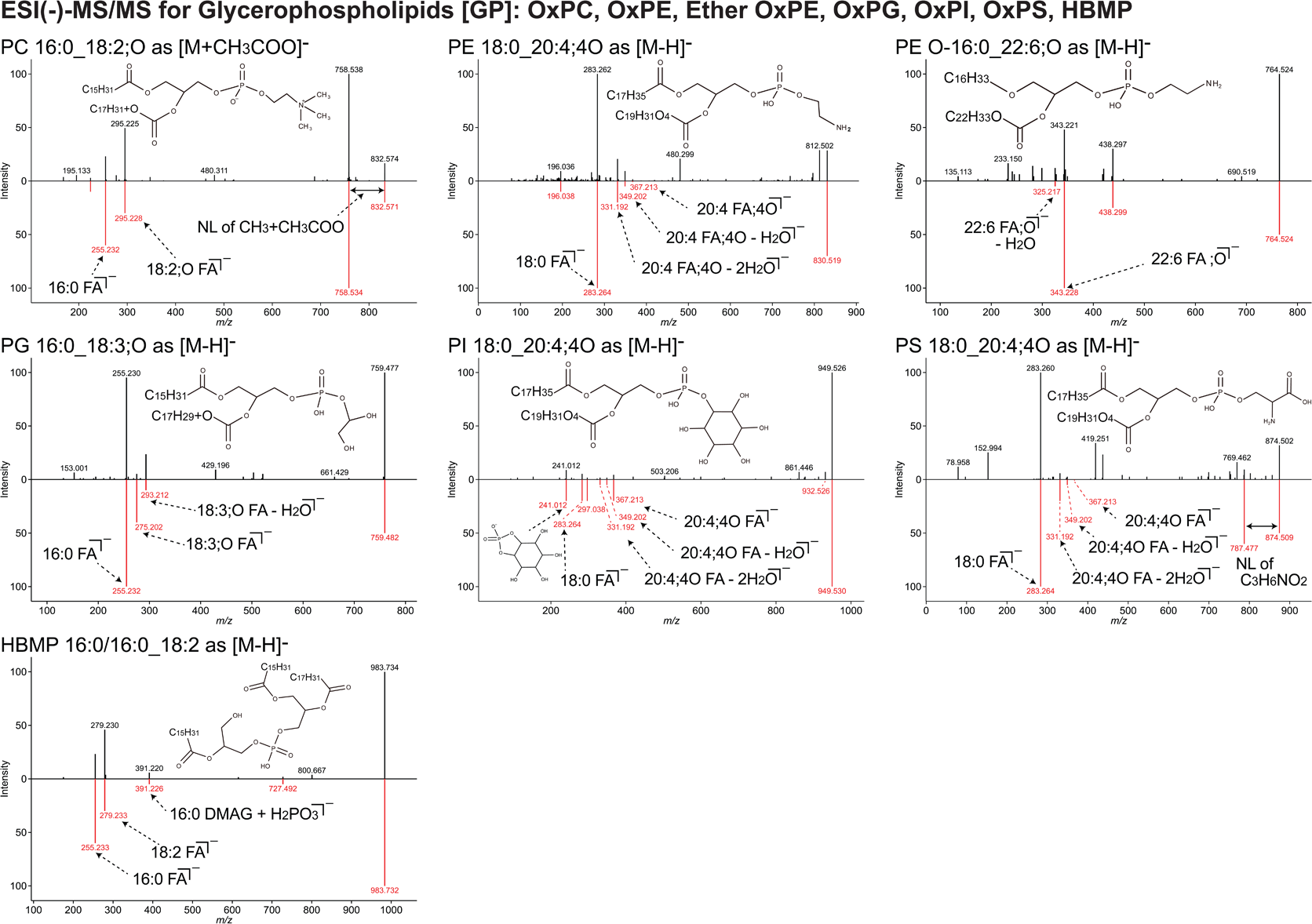

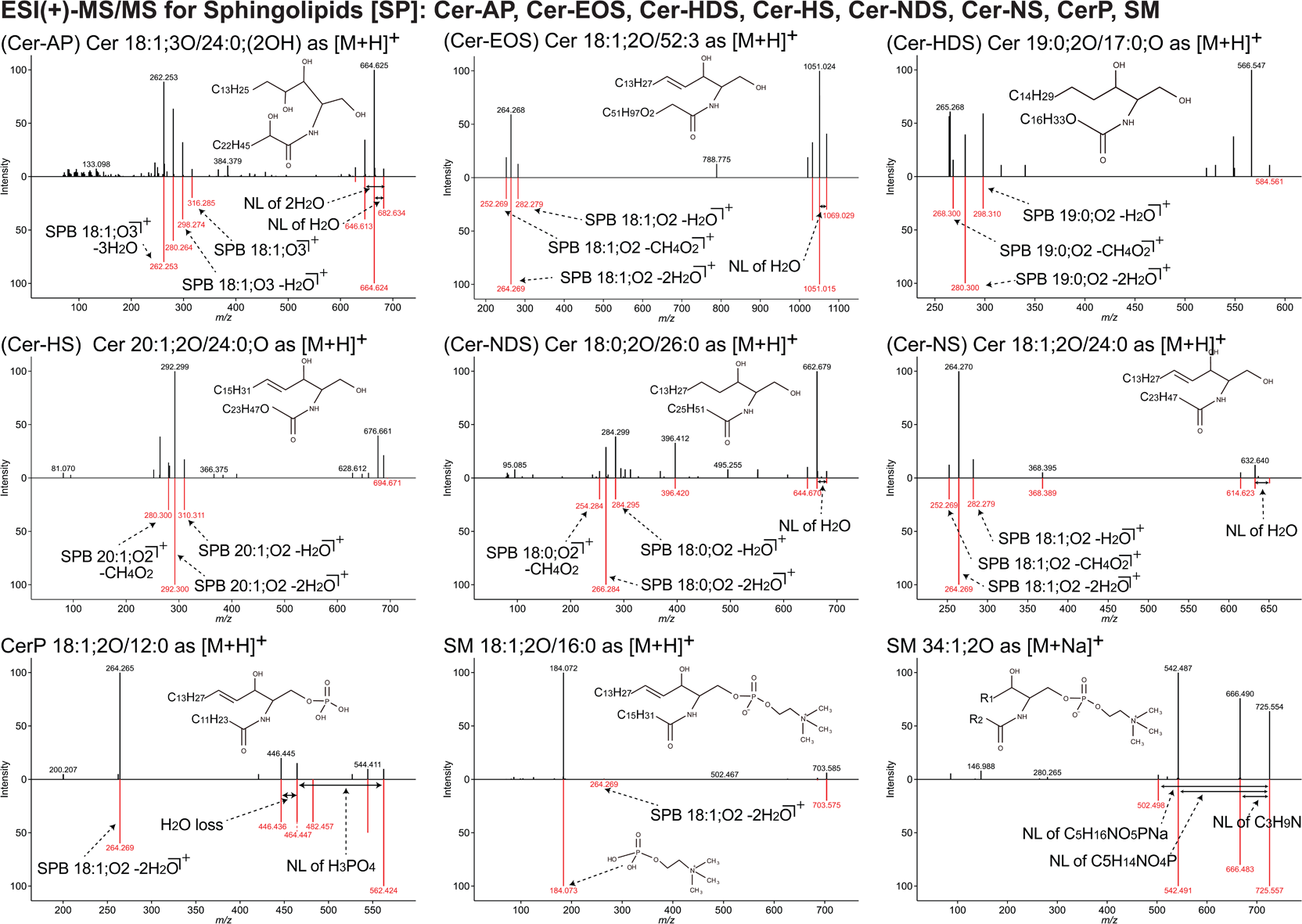

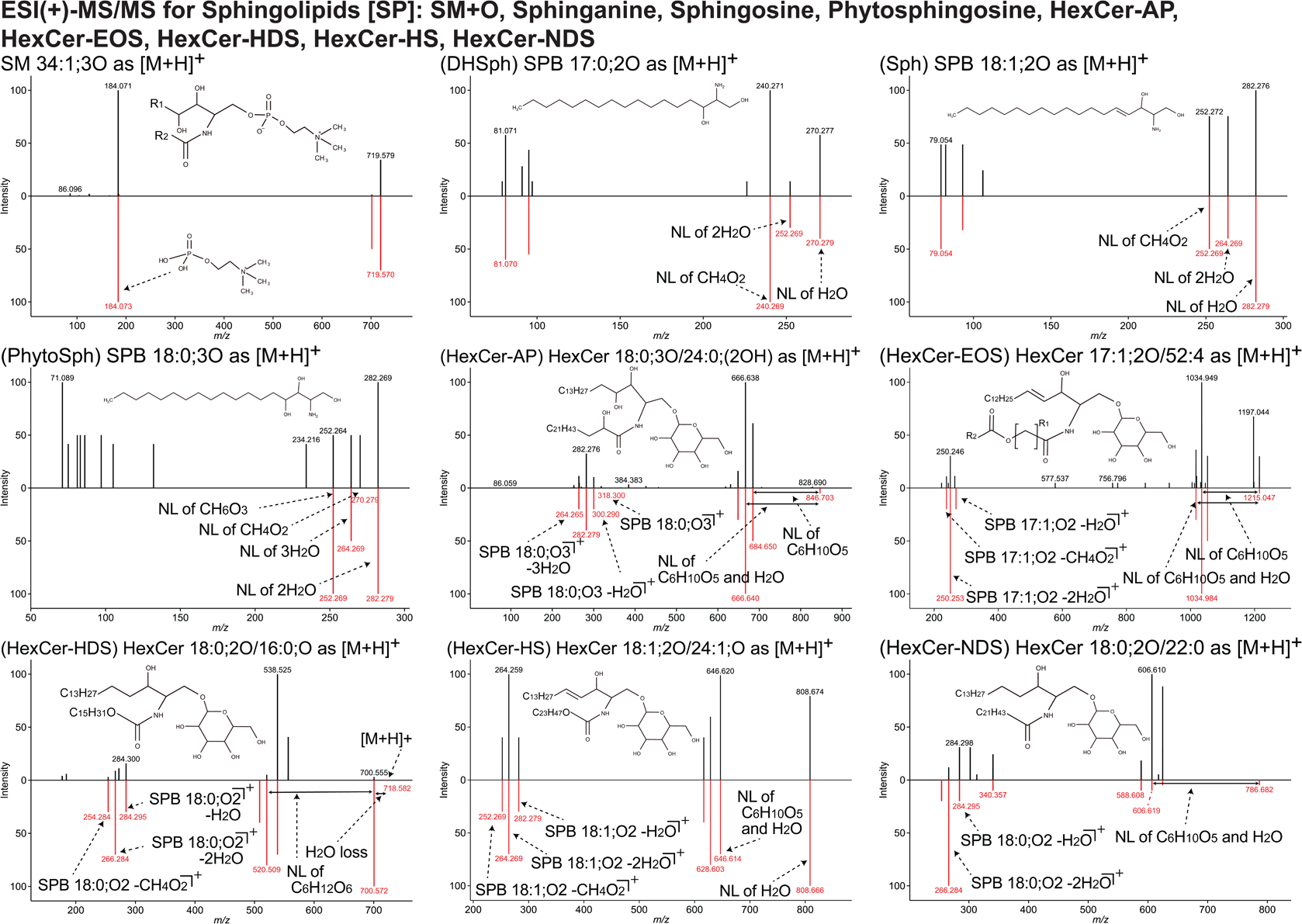

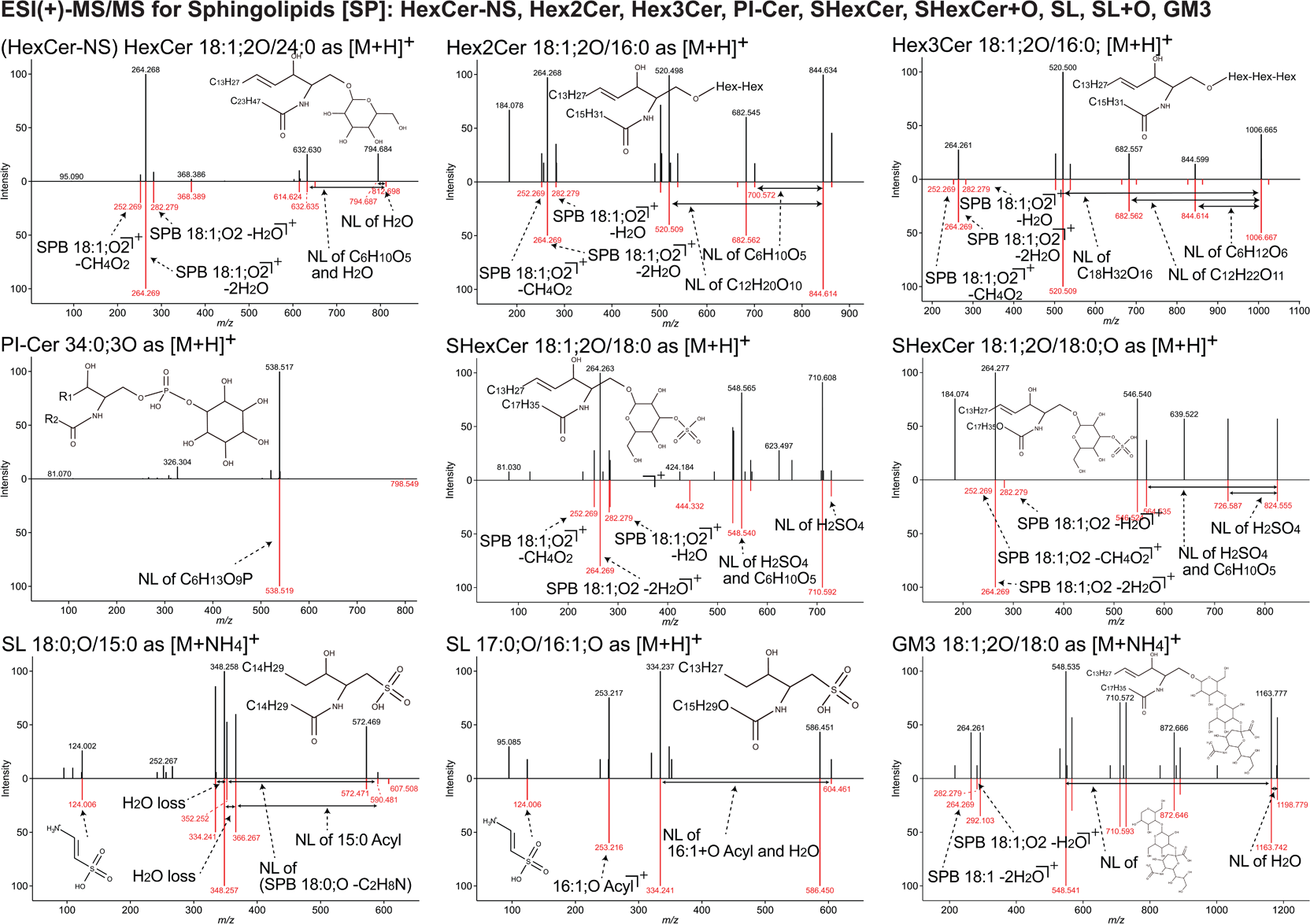

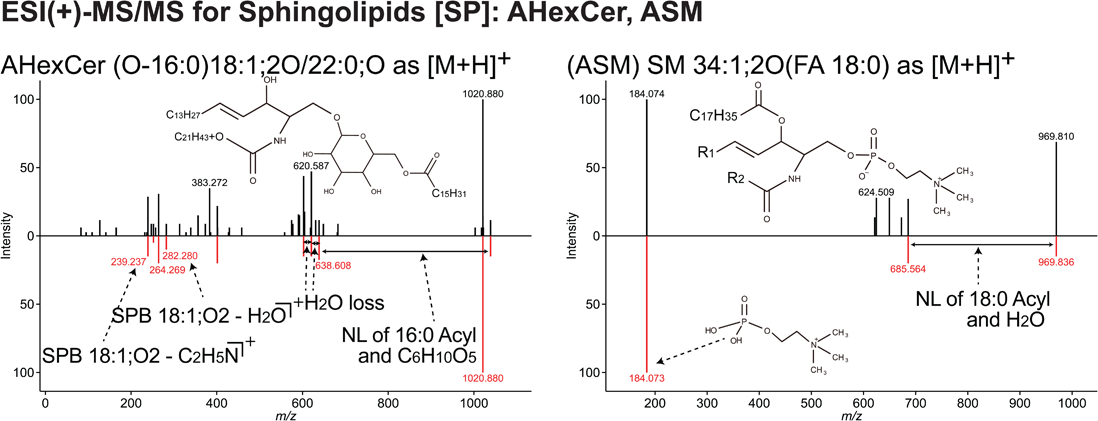

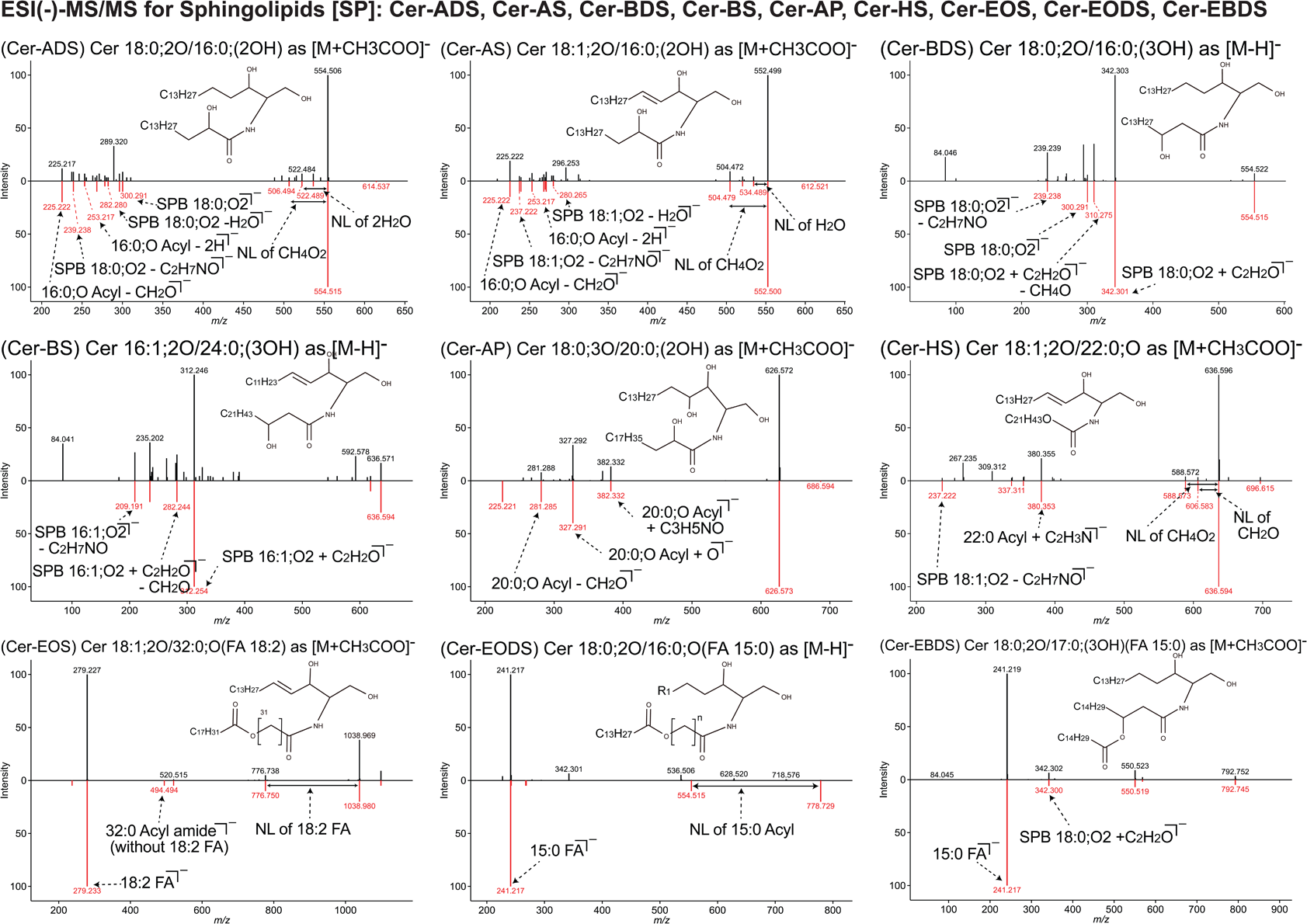

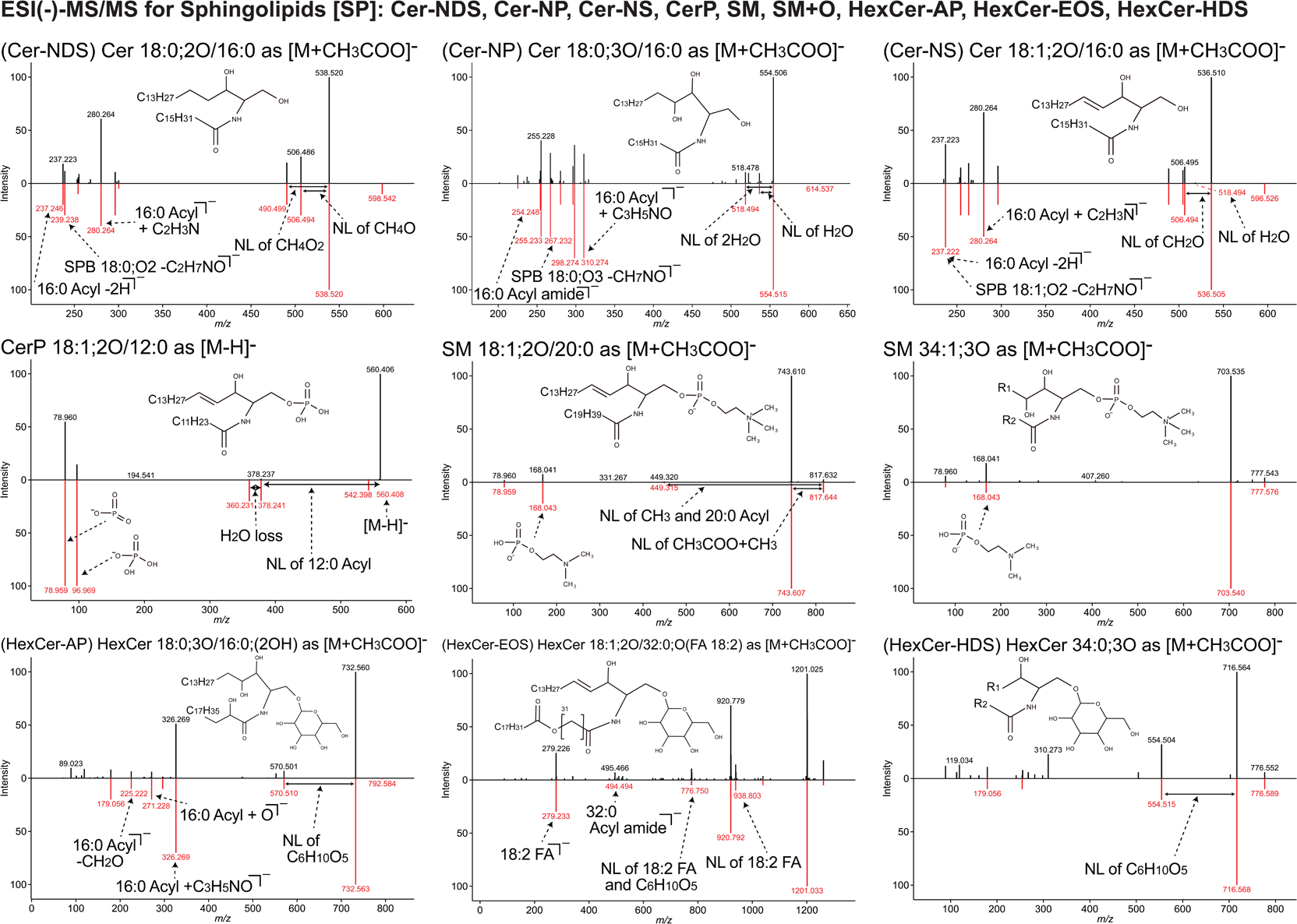

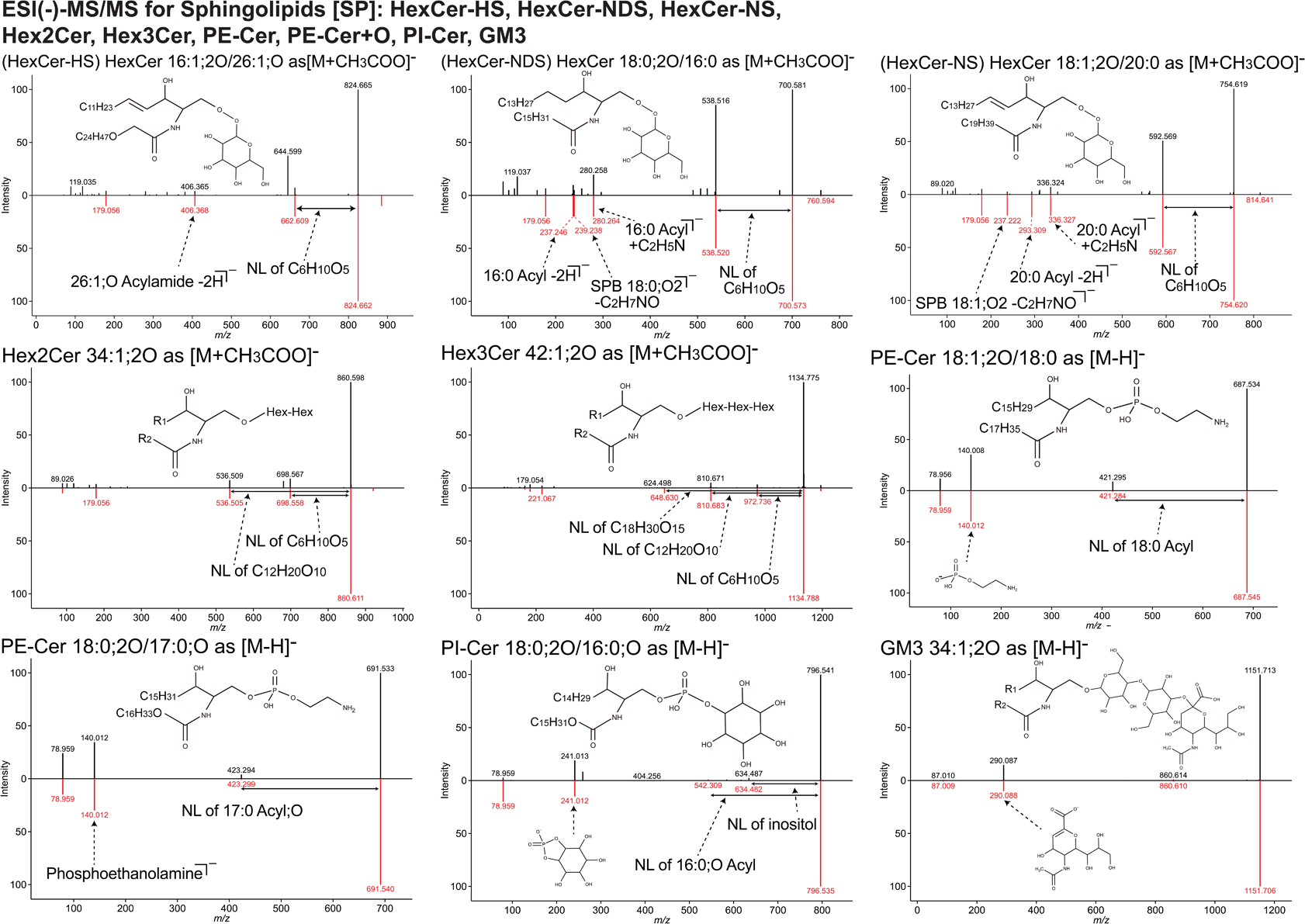

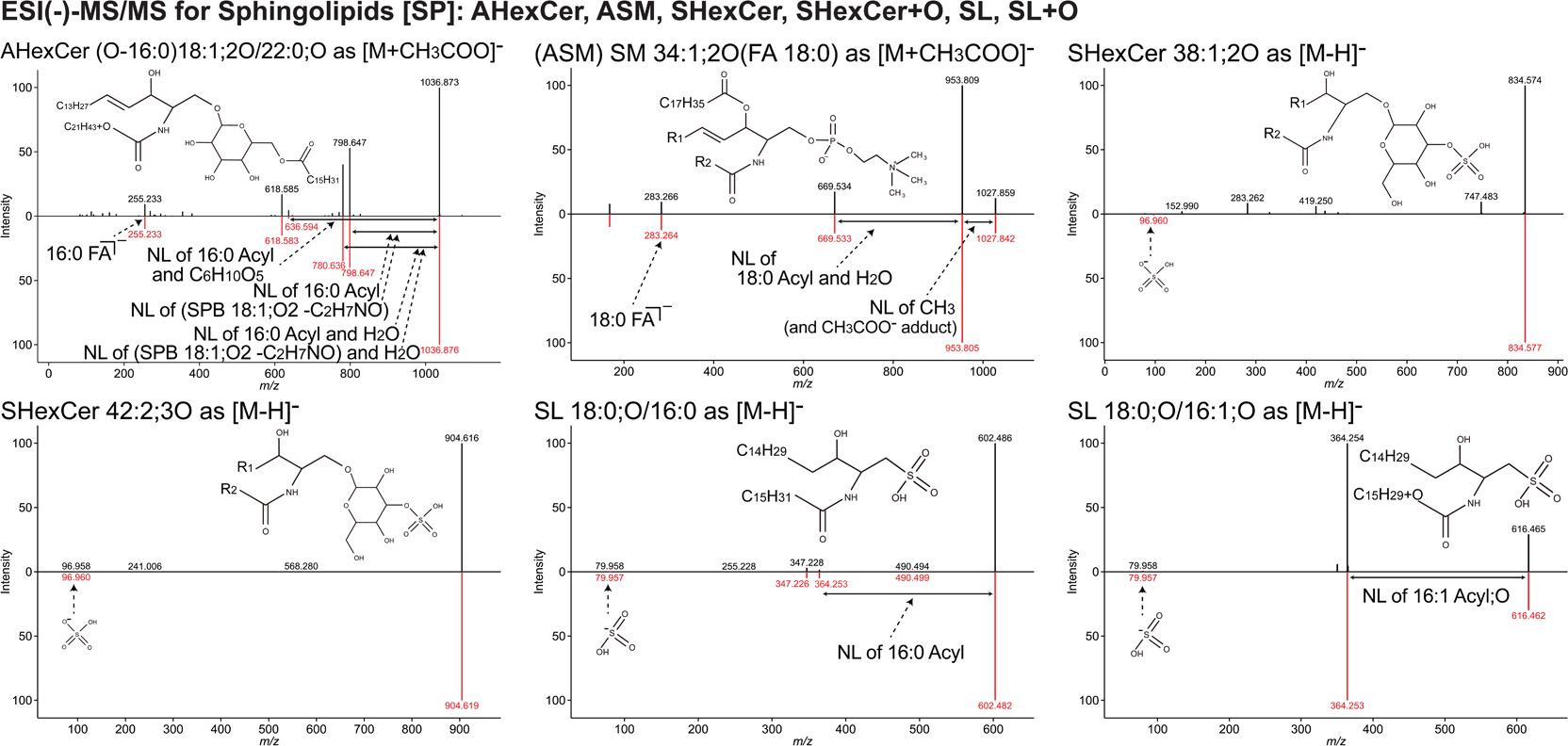

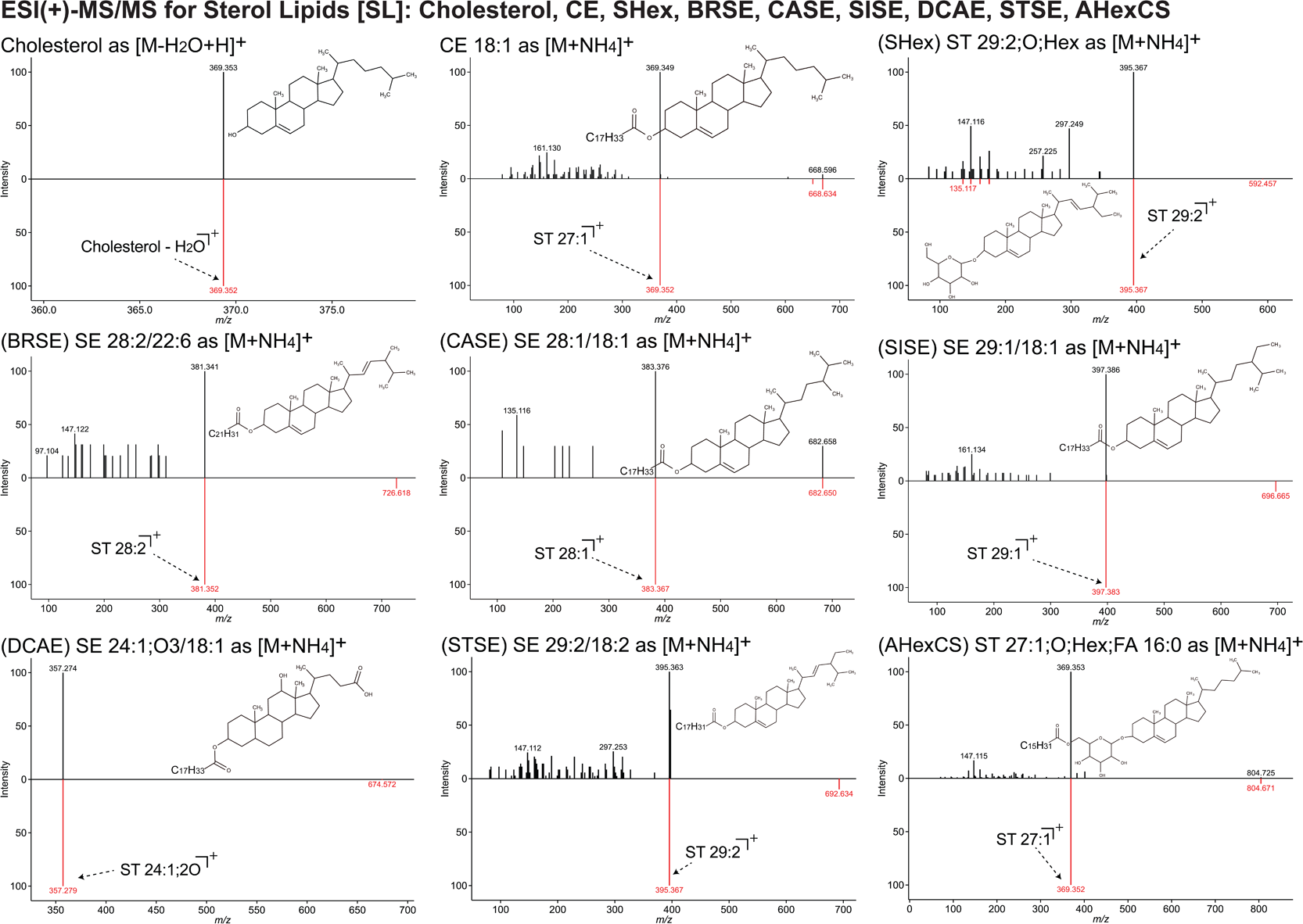

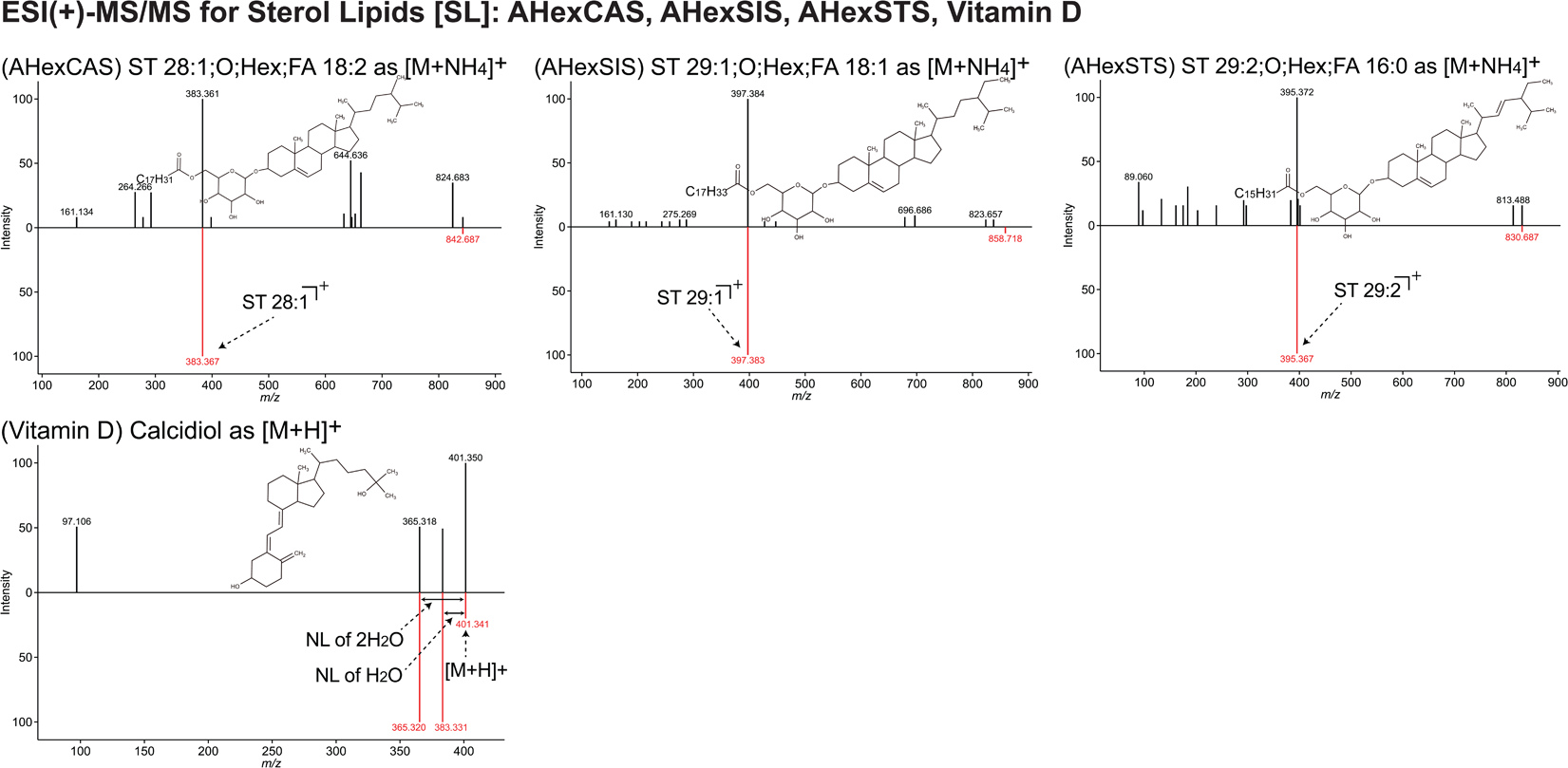

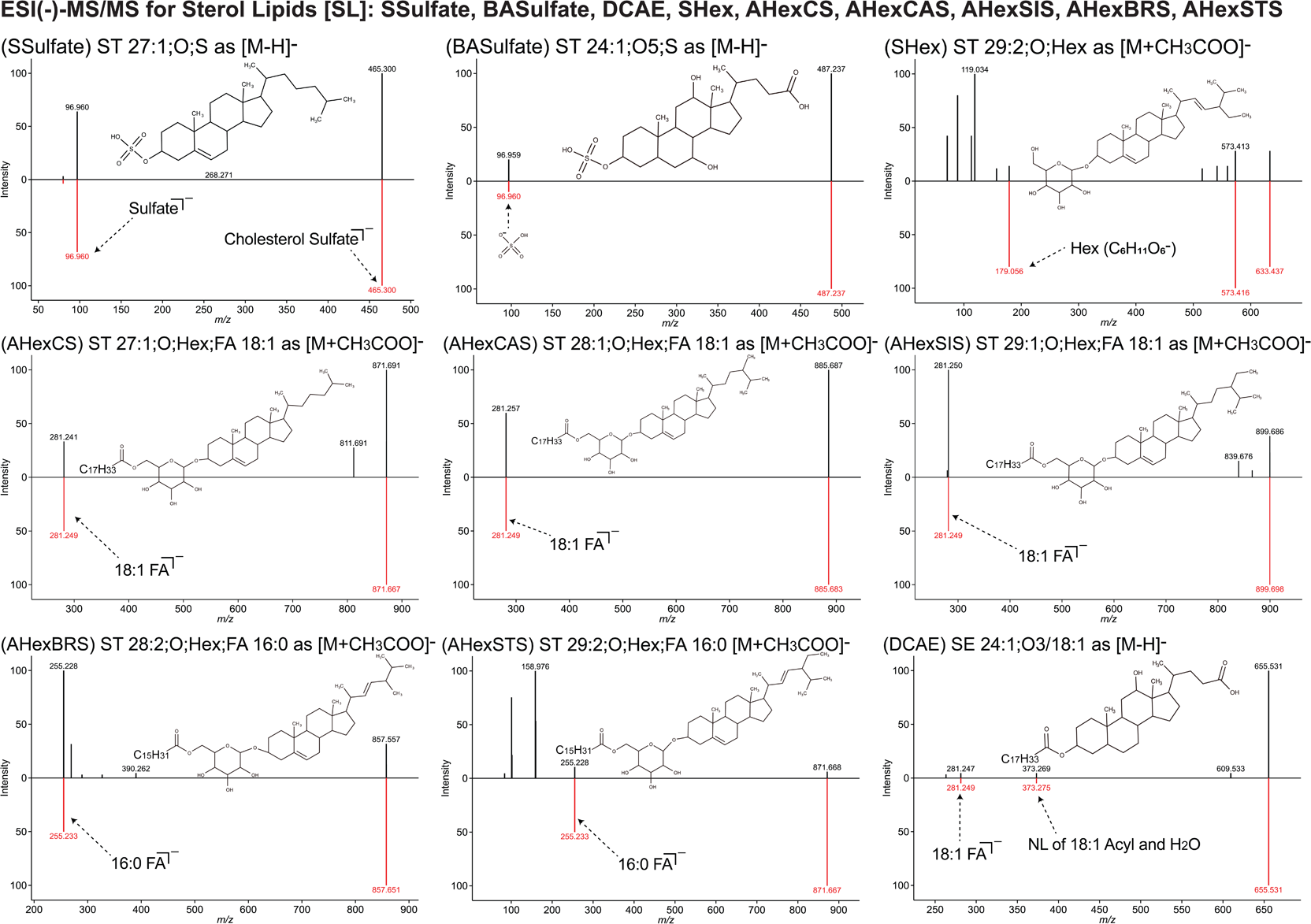

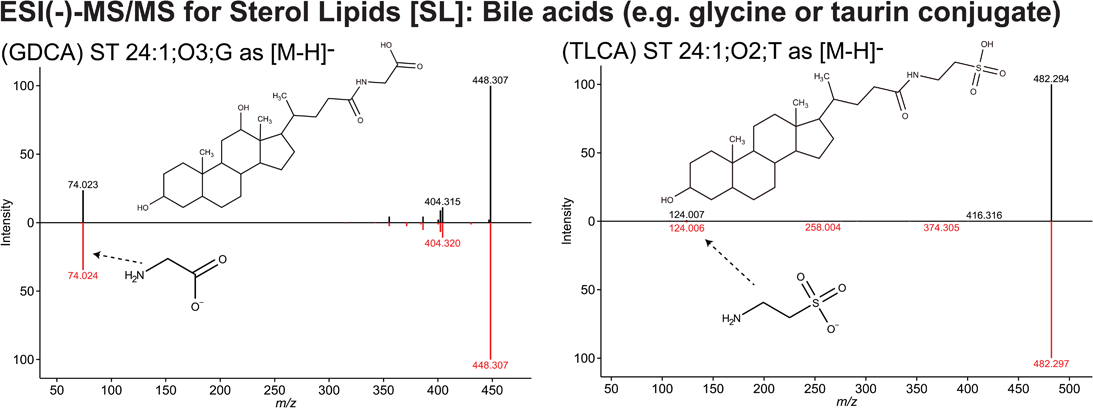

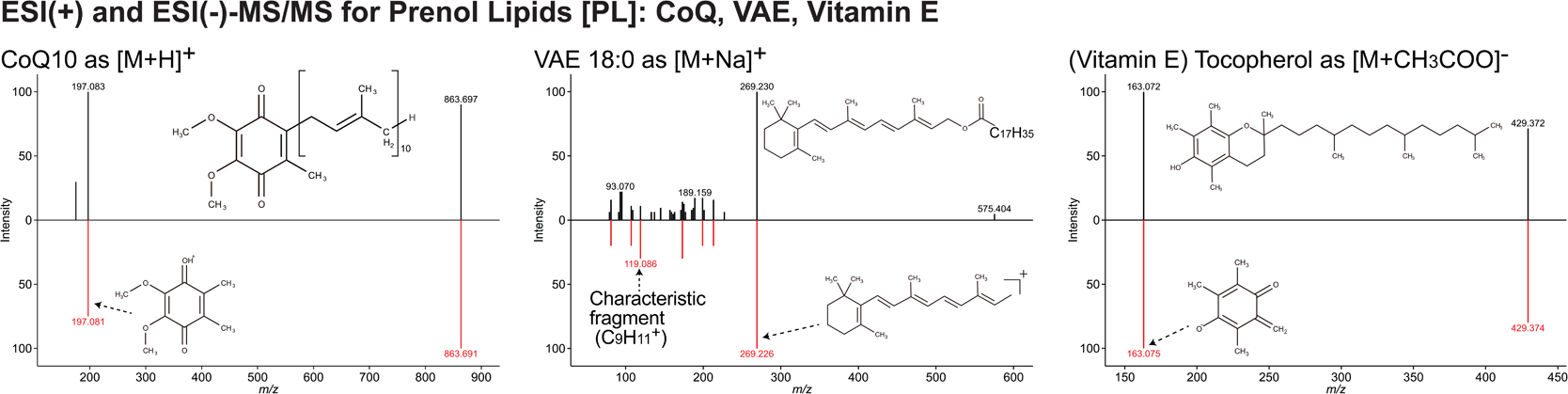

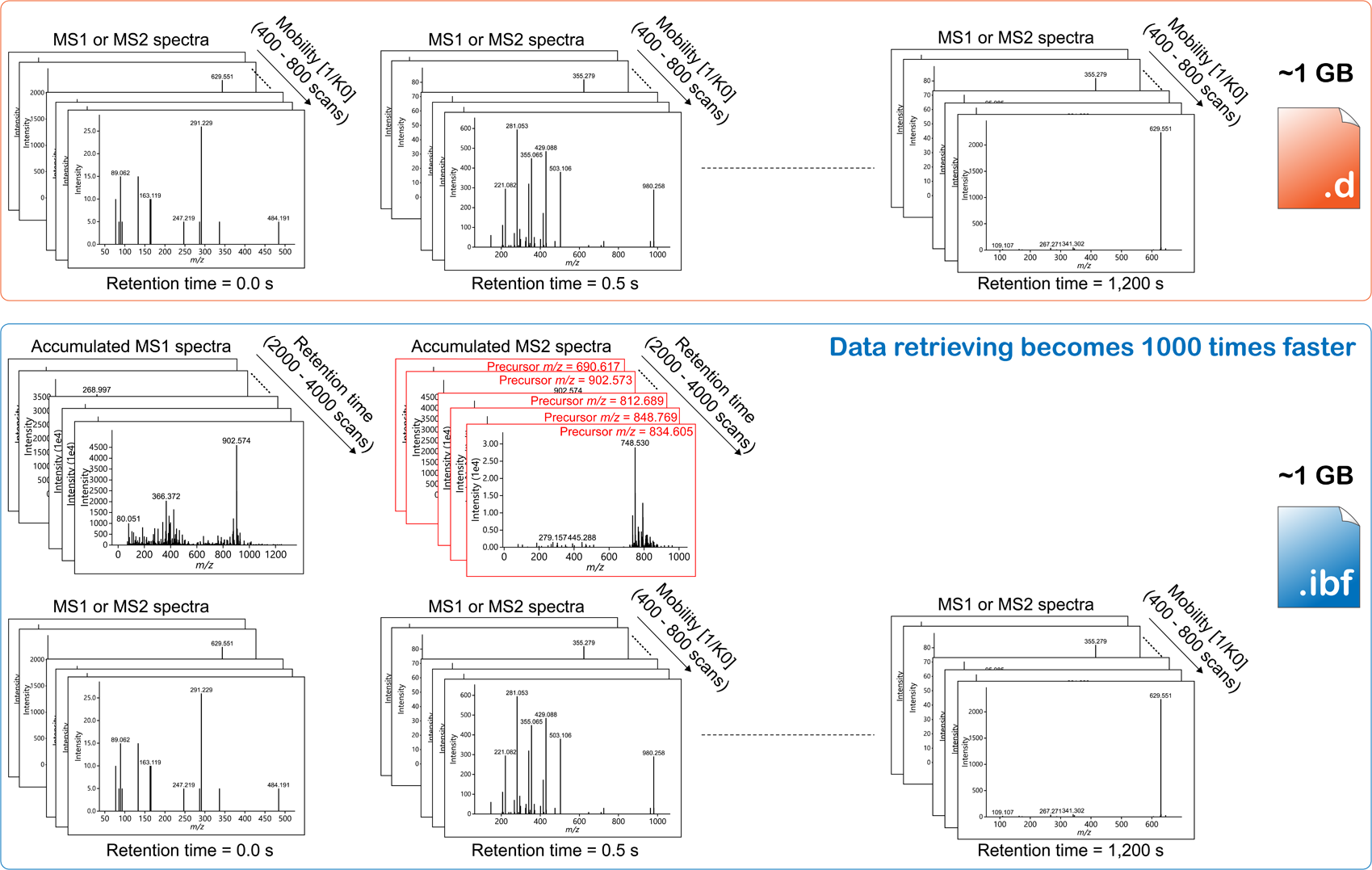

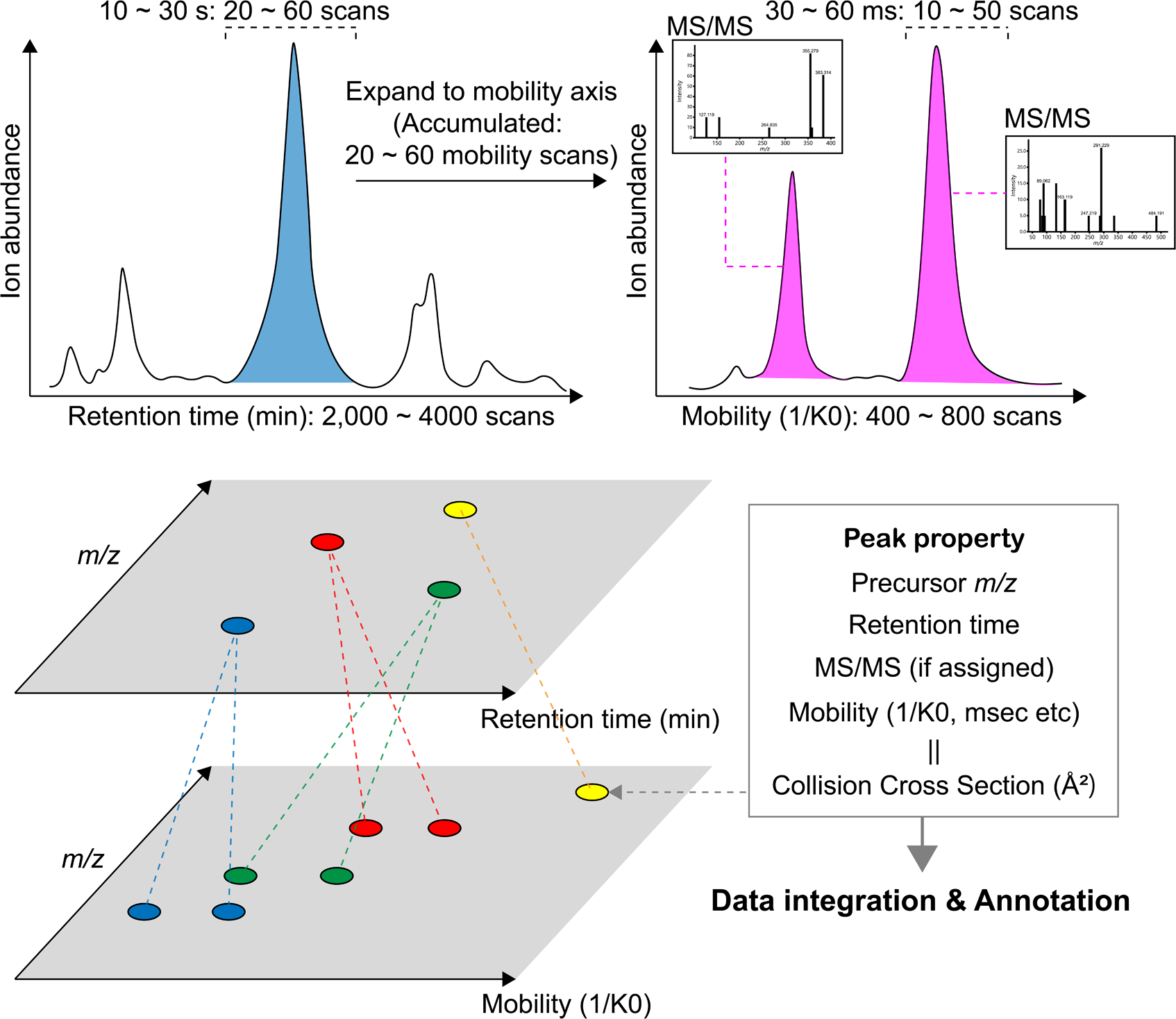

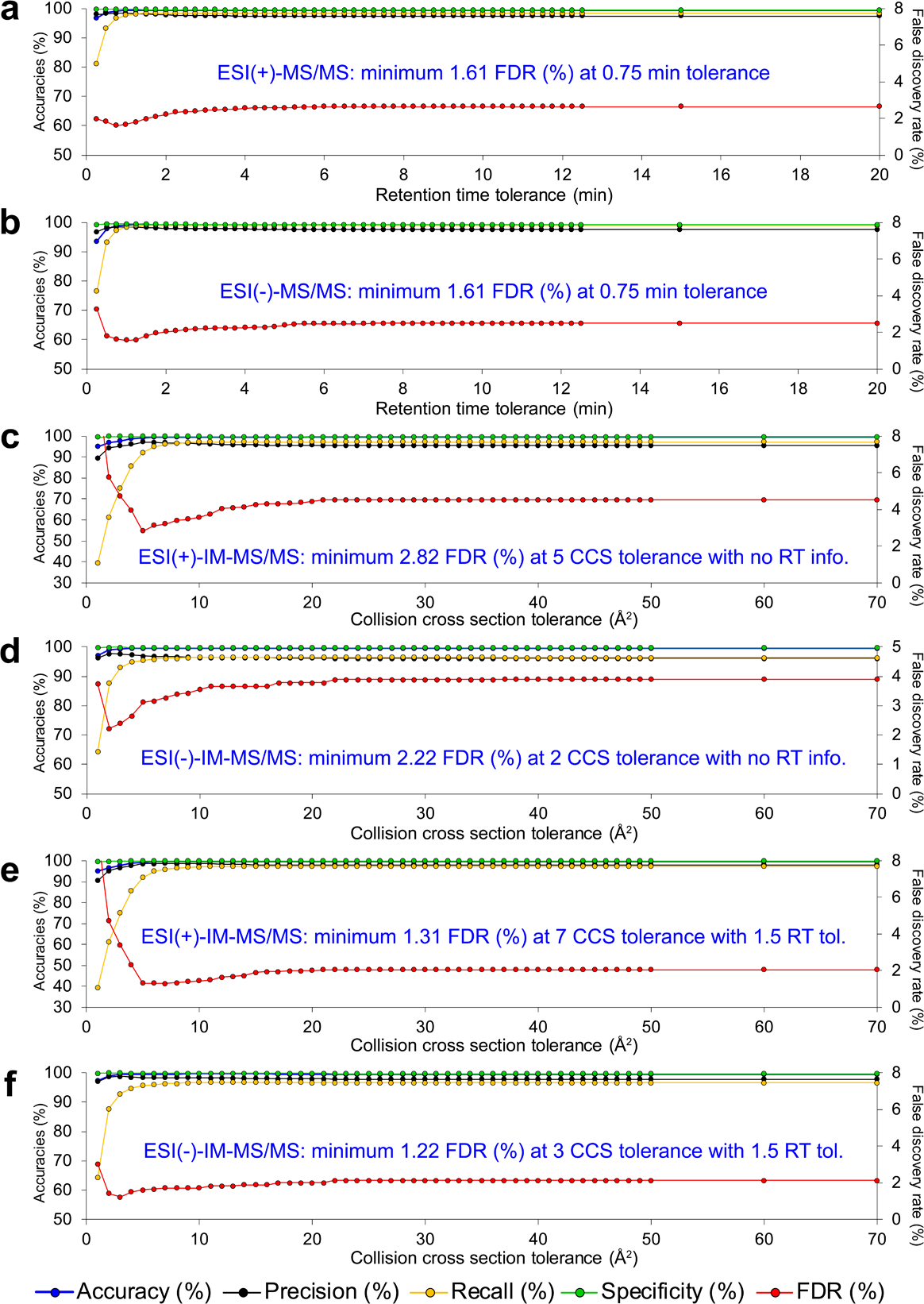

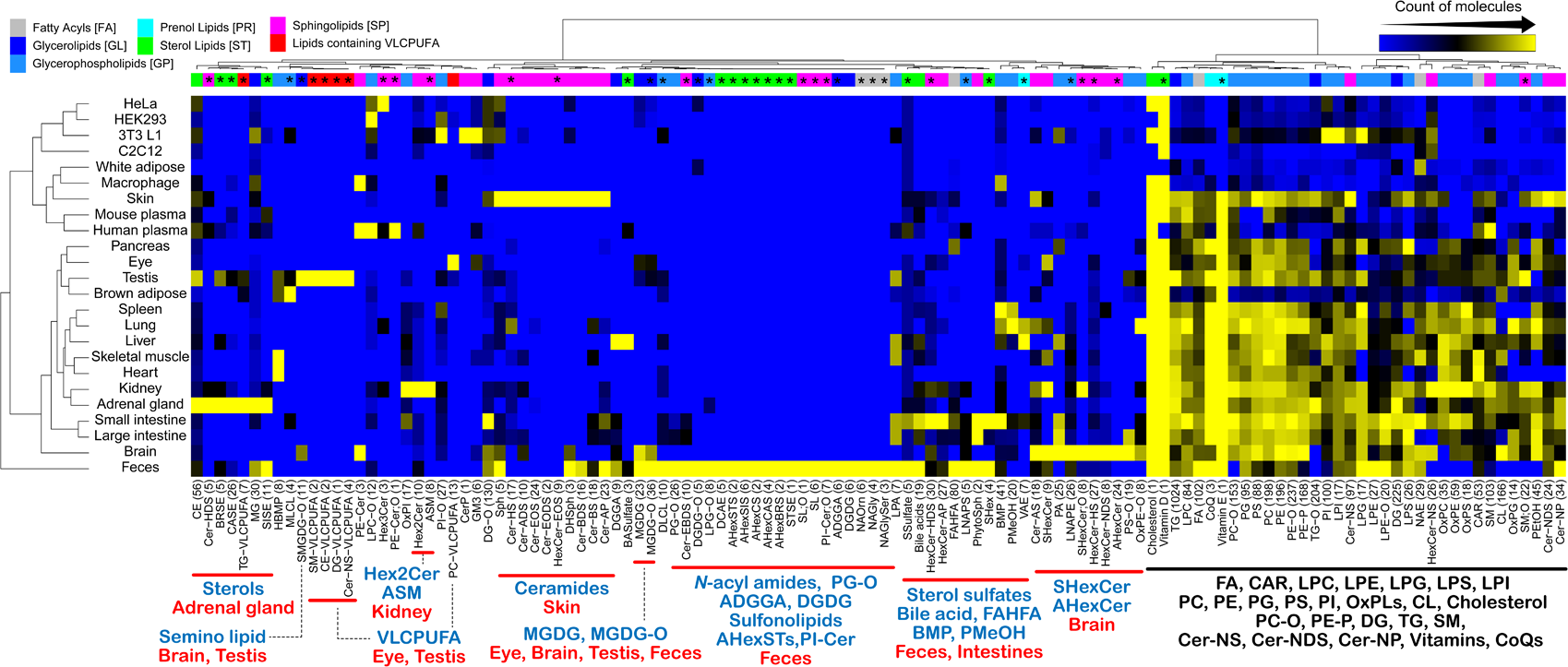

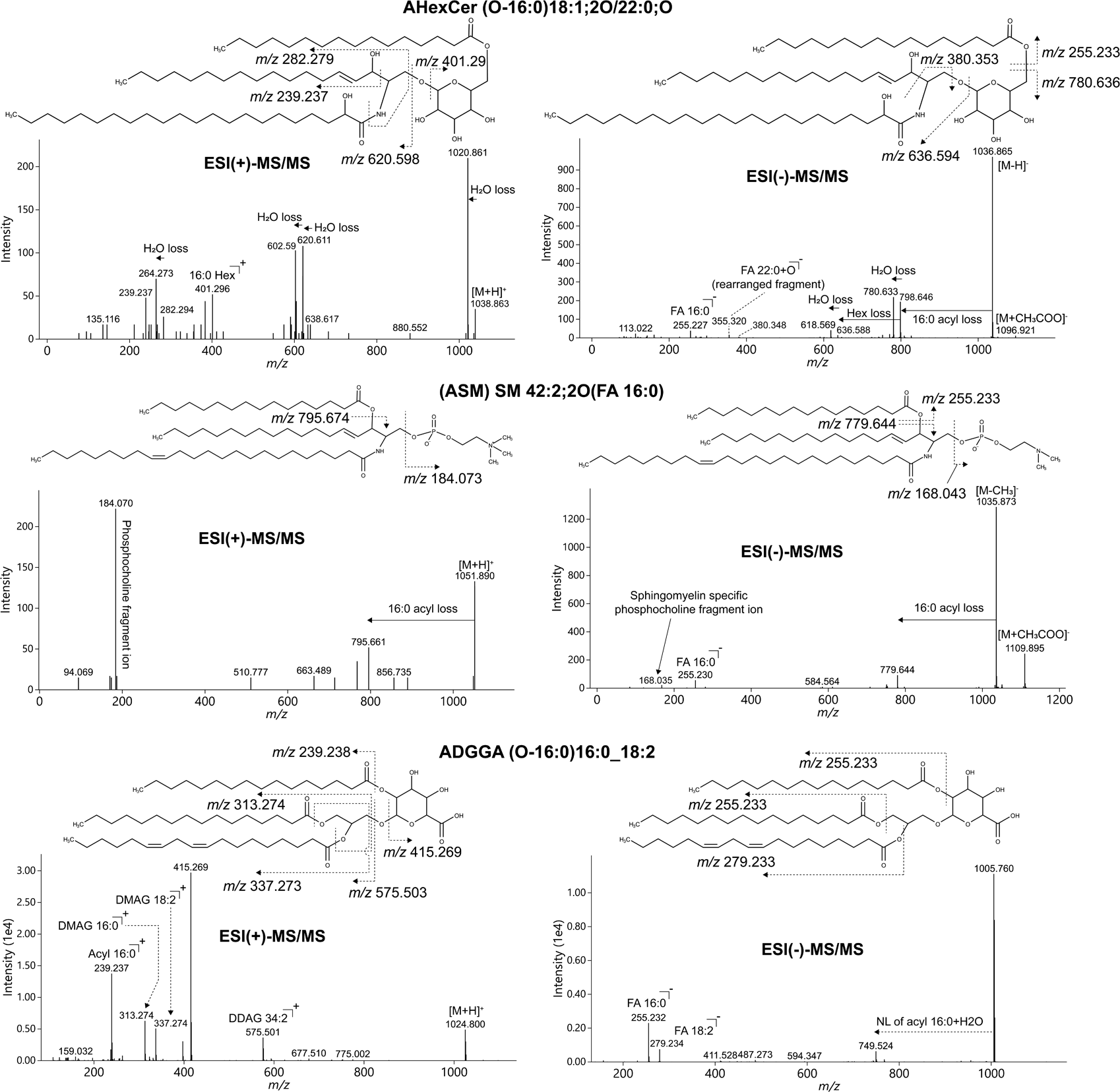

