## Supplementary Methods, Note 1, and Note 2 for "MS-DIAL 4: accelerating lipidomics using an MS/MS, CCS, and retention time atlas"

### Supplementary Figures

**Fig. S1 | Summary of the hybrid scoring system for lipid tandem mass spectrometry (MS/MS) annotations.** The MS/MS spectrum of the acetate adduct form of sphingomyelin (SM) 18:1;2O/16:0 (*m/z* 761.581) is shown as an example. The characteristic fragment ion of *m/z* 168.043 ($C_4H_{11}NO_4P$) and neutral loss of 74.037 Da ($CH_3COO + CH_3$) to specify the SM lipid class were detected whereas the product ion of *m/z* 449.315 as the neutral loss of acyl 16:0 to characterize the *N*-acyl chain moiety was observed in the MS/MS spectrum. The classical spectral similarity calculation using dot- and reverse dot product scores is executed to filter out noisy spectra: the MS/MS spectrum is recognized as unknown if the dot product is <0.1 or the reverse dot product is <0.5. For the decision tree algorithm for the acetate adduct form of SM lipid, the fragment existence of SM-specific fragment ions is evaluated, in which the relative abundance cut off is also utilized. If the neutral loss of *N*-acyl chain (as observed by *m/z* 449.315) is detected and the relative abundance is >0.1% when compared with the base peak of *m/z* 687.550 (recognized as 100%), the lipid structure of "molecular species level" is represented as SM 18:1;2O/16:0. If no acyl chain fragment exists, the description of "species level" is exported as SM 34:1;2O.

**Fig. S2 | Details of tandem mass spectrometry (MS/MS) characterization of seven lipid subclasses categorized as LipidMAPS Fatty Acyls [FA].** The important fragment ions and neutral losses for characterizing the lipid subclass at the molecular species level are described. The details of fragment descriptions are provided in **Supplementary Note 2**.

**Fig. S3 | Details of tandem mass spectrometry (MS/MS) characterization of 17 lipid subclasses categorized as LipidMAPS Glycerolipids [GL].** The important fragment ions and neutral losses for characterizing the lipid subclass at the molecular species level are described. The details of fragment descriptions are provided in **Supplementary Note 2**.

**Fig. S4 | Details of tandem mass spectrometry (MS/MS) characterization of 36 lipid subclasses categorized as LipidMAPS Glycerophospholipids [GP].** The important fragment ions and neutral losses for characterizing the lipid subclass at the molecular species level are described. The details of fragment descriptions are provided in **Supplementary Note 2**.

**Fig. S5 | Details of tandem mass spectrometry (MS/MS) characterization of 37 lipid subclasses categorized as LipidMAPS Sphingolipids [SP].** The important fragment ions and neutral losses for characterizing the lipid subclass at the molecular species level are described. The details of fragment descriptions are provided in **Supplementary Note 2**.

**Fig. S6 | Details of tandem mass spectrometry (MS/MS) characterization of 17 lipid subclasses categorized as LipidMAPS Sterol Lipids [ST].** The important fragment ions and neutral losses for characterizing the lipid subclass at the molecular species level are described. The details of fragment descriptions are provided in **Supplementary Note 2**.

**Fig. S7 | Details of tandem mass spectrometry (MS/MS) characterization of three lipid subclasses categorized as LipidMAPS Prenol Lipids [PR].** The important fragment ions and neutral losses for characterizing the lipid subclass at the molecular species level are described. The details of fragment descriptions are provided in **Supplementary Note 2**.

**Fig. S8 | Concept of the ion mobility base framework (IBF) format.** The raw data of Bruker's parallel accumulation-serial fragmentation (PASEF) are shown as an example. In the raw data format (.d), 400–800 spectral records along with the changes of trapped electric field are stored at each retention time (RT) point, and their recording is sequenced by the end of the liquid chromatography (LC) condition. The file size of our 20 min LC condition was approximately 1 GB. The IBF format (.ibf) was designed to rapidly access ion mobility (IM)-MS data and the accumulated ion signals summing the mass spectral data of IM axis at the same RT bin were also stored to achieve a rapid peak picking process in the RT and *m/z* dimensions. The calibrant information such as beta coefficient and intercept values for the Agilent single field method and t0, exponent, and coefficient values for the Waters collision cross section calculation were also stored in the IBF format. The file size was roughly equivalent to the original size, and data retrieval could be accomplished within 10 s.

**Fig. S9 | Scheme of peak picking for liquid chromatography coupled with ion mobility tandem mass spectrometry (LC-IM-MS/MS).** The extracted ion chromatogram (EIC) for a certain *m/z* value with approximately 10 ppm mass tolerance is constructed by using accumulated MS1 spectral panels, which are constructed by summing spectra of the IM axis with three-decimal binning for *m/z* value. After the peak detection method is performed in the retention time (RT) axis, the extracted ion mobilogram (EIM) is constructed in the IM axis by accumulating ions from the left- and right edges of the detected peak in the RT axis, followed by the peak detection procedure applied to the EIM. For example, the average peak width on the RT dimension is 10–30 s whereas the average peak width on the IM dimension is 30-60 ms in our LC-IM-MS/MS condition. The MS/MS spectrum from data-dependent/independent acquisition methods is assigned to each peak on the IM axis. As a result, each peak spot in the RT and *m/z* dimension has more than one peak spot in the mobility and *m/z* dimension, and the peak contains peak height, peak area, RT, mobility value, *m/z*, and MS/MS as the peak properties. The collision cross-section (CCS) is calculated using the Mason–Schamp equation for Bruker trapped IM, the single field CCS method for Agilent drift tube IM, and the IM calibration function for Waters travelling wave IM.

**Fig. S10 | Evaluations for the MS-DIAL 4 annotation pipeline.** The accuracy, precision, recall, specificity, and false discovery rate (FDR; %) were calculated by using the set of true positives (TPs) and true negatives (TNs), which are available as **Supplementary Data 2**. **a**, The set of 12,798 TPs and 64,121 TNs from liquid chromatography-electrospray ionization-positive-tandem mass spectrometry (LC-ESI(+)-MS/MS) (data dependent acquisition, DDA) data was processed where the minimum FDR value was 1.61% at 0.75 min retention time (RT) tolerance. **b**, The set of 10,600 TPs and 30,290 TNs from LC-ESI(−)-MS/MS-DDA was processed, where the minimum FDR value was 1.61% at 0.75 min RT tolerance. **c**, The set of 2,598 TPs and 30,131 TNs from LC-ESI(+)-IM-MS/MS (PASEF) was processed with various collision cross-section (CCS) tolerances without RT information, where the minimum FDR was 2.82% at 5 $Å^2$ CCS tolerance. **d**, The set of 1,670 TPs and 20,737 TNs from LC-ESI(−)-IM-MS/MS (PASEF) was processed with no RT, where the minimum FDR was 2.22% at 2 $Å^2$ CCS tolerance. **e**, The same set as used in (**c**) was processed with 1.5 RT tolerance, where the minimum FDR was 1.31% at 7 $Å^2$ CCS tolerance. **f**, The same set as in (**d**) was processed with 1.5 RT tolerance, where the minimum FDR was 1.22% at 3 $Å^2$ CCS tolerance.

**Fig. S11 │ Mapping lipid diversity mammalian tissues and cells.** Hierarchical clustering analysis was performed using the data matrix containing the count of molecules categorized to each lipid subclass; the count was scaled from −1 to 1. Overall, 112 lipid subclasses in addition to PC, CE, DG, TG, Cer-NS and SM subclasses containing very-long-chain poly unsaturated fatty acid (VLCPUFA) are described in the x-axis; total counts of molecular species annotated in each lipid subclass are in brackets. The LipidMaps category and a specialized lipid class containing VLCPUFA are given. Asterisks indicate the lipid subclasses that only MS-DIAL 4 can characterized when compared with existing lipidomics software tools evaluated in this study. The lipid nomenclature is detailed in **Supplementary Table 2**. Note that although potentially quantitative, no detection does not evince the nonexistence of certain lipid subclasses because of technical limitations.

**Fig. S12 | Prediction of putative lipid structures by untangling the tandem mass spectrometry (MS/MS) spectra. a**, Fragment assignments for acylated hexosyl ceramide (AHexCer) containing sphingosine (18:1;2O), *N*-acyl chain (22:0;O), and *O*-acyl chain (O-16:0) are shown. Compared to the *O*-acyl chain positional isomer (HexCer-EOS), the fragment ion of Hex-*O*-acyl 16:0 (*m/z* 401.29) is observed in positive ion mode, and no hexosyl loss (162 Da) is found before the *O*-acyl 16:0 loss (238 Da) in negative ion mode. The behaviors were reproducible in all of the 24 different molecular species of AHexCer annotated in this study. **b**, Fragment assignments for acylated sphingomyelin (ASM) containing sphingosine (18:1;2O), *N*-acyl chain (24:1), and *O*-acyl chain (O-16:0) are shown although the sphingobase and *N*-acyl chain moieties cannot be determined at the molecular species level at least in our experimental condition. The SM specific fragment ions can be observed in both positive and negative ion MS/MS spectra. In addition, the product ion derived from the fatty acid moiety (*O*-acyl 16:0) was detected in both ion modes although they cannot be observed in usual SM lipid species. These behaviors were also true in all of the eight different molecules of ASM lipid subclass found in this study. **c**, An example of acylated uronosyldiacylglycerol (ADGGA) containing 16:0 and 18:2 acyl chains in the glycerol moiety and *O*-16:0 acyl chain in the uronosyl moiety is described. The fragment ions of HexA-*O*-acyl 16:0 (*m/z* 415.269) and acyl 16:0 are observed in positive ion mode where HexA denotes the uronic acid moiety.

These behaviors were true in all of the 28 different molecules of the ADGGA lipid subclass found in this study.

### Supplementary Tables

**Table S1 | Metadata of biological samples for Supplementary Data 1. a,** The sample category (i.e., human, mouse, human cultured cell, mouse cultured cell, plant, and algae), sample name (e.g., mouse tissue, cell type, and species), analysis method number, institute where the analysis was performed, mass spectrometry (MS) machine name, injection volume, sample volume, and internal standard information incorporated in this study were described. The analysis method is detailed in **Supplementary Note 1**. **b,** Information regarding the internal standards is described. From sample number 14 to 17, the peak abundance was normalized using the concentration of phosphorus in the lipid extraction solvent. For plant samples, all lipids were normalized using phosphatidylcholine 10:0_10:0 because of the internal standard content. For the other samples, the lipid quantifications were calculated as described in **Supplementary Table S2**.

**Table S2 | Details of lipid subclass characterization. a,** The LipidMAPS classification, lipid ontology, example nomenclature for species level- and molecular species-level annotations, formula, SMILES, and InChIKey were described. For each example lipid species, the diagnostic fragments for characterizing the lipid subclass and acyl chain compositions are described, with the example structure being consistent with the structure described in **Supplementary Figs. 2–7**. The confirmation level is also described: level 1, confirmed by the authentic standard compound; level 2, elucidated by the fragment evidence along with the reported literature; and level 3, putatively assigned novel structures by accumulating indirect evidence of the fragment ions in positive and negative ion modes. The internal standard was dissolved using methanol, chloroform, or the mixture, and analyzed by method 1 or 2 as described in **Supplementary Note 1**. **b,** The adduct form for lipid quantification that is registered in our lipidomics quant database was shown. Although the acetate adduct form is detected in our experimental condition, the information can be replaced by that for the formate adduct when ammonium formate is used as the modifier for the liquid chromatography (LC) solvent. The internal standard for each lipid subclass is described for all internal standard solutions described in **Supplementary Table S1**. If the abbreviation of lipid subclass is the same as that of the internal standard component, the quantification level becomes 2, otherwise 3.

**Table S3 | Comparison with other lipidomics software tools. a,** The evaluated tool name, version, available format, and the function/utility existence for peak detection/picking, peak alignment/data integration, and tandem mass spectrometry (MS/MS) annotation method are described. The count of annotatable lipid subclasses by the default library is also described. **b,** The general comparison between MS-DIAL 4 and others including MS-DIAL 3, LipiDex, LipidMatch, LIQUID, LipidHunter2, Greazy, and MzMine2 was performed. The lipid subclasses which can be characterized in MS-DIAL 4 are described, and the term 'Yes' is added if the lipid subclass was characterized in this study. Moreover, the references are described if the lipid class has already been reported elsewhere as the part of MS-DIAL 3 (or other versions). **c,** The result obtained by analyzing the liquid chromatography (LC)-MS/MS data of the standard mixture is described. The origin of standard compound is also described. **d,** The count of annotated lipids in the LC-MS/MS data analyzing NIST SRM 1950 plasma is shown. The parameters used for each software program are described in (**e,f,g,h,i,j,k**).

**Table S4 | Summary of named lipids from 1,056 lipidomics data in total.** All lipids were annotated as species level (class resolved) or molecular species level (chain resolved). The term of "lipid assigned" was used for species level annotation of vitamins, bile acids, sterols, and their conjugates, Because both results of liquid chromatography-tandem mass spectrometry (LC-MS/MS) and LC-ion motility (IM)-MS/MS were integrated, the "−1" value was inserted for drift time and collision cross-section (CCS) information in the results of conventional LC-MS/MS methods. **a,** The summary of our lipidomics results is shown. **b,** Information for the unique lipid "ion" in which the duplicates from different adduct forms of the same lipid molecule were not excluded is described. **c,** The unique lipid molecules without duplicate structure descriptions are described.

**Table S5 | Validation of NIST SRM 1950 lipidomics data using LipidQC.** The LipidQC program was downloaded at http://secim.ufl.edu/secim-tools/lipidqc/. The lipid nomenclature of our lipidomics results was reformatted to be used in LipidQC for evaluating semi-quantitative values of human plasma lipid concentrations. **a,** The median of platform means (MDMEs) among eight platforms was validated. The validations for plasma lipidomics results from method 1 ($n$ = 5), method 2 ($n$ = 5), method 5 ($n$ = 5), method 6 ($n$ = 5), method 7 ($n$ = 4), method 8 ($n$ = 3), method 9 ($n$ = 4), and method 10 ($n$ = 6) were described in **b, c, d, e, f, g, h,** and **i,** respectively, where $n$ denotes for the number of technical replicates.

**Table S6 | Training set for retention time (RT) prediction.** A total of 4,303 lipids of 108 lipid subclasses annotated at the molecular species level are described. The pairs of RT and representative structure description using SMILES and InChIKey are shown in addition to 152 chemical descriptors generated by PaDEL.

**Table S7 | Training set for collision cross-section (CCS) prediction.** A total of 3,570 lipids of 101 lipid subclasses annotated at the molecular species level are described. The pairs of CCS and representative structure description using SMILES and InChIKey are shown in addition to 328 chemical descriptors generated by PaDEL.

**Supplementary Data**

**Data S1 | Results of lipid profiling for each biological study.** For ion mobility (IM) data (nos. 47–62), the quantification table was divided into two tables. One is based on the quantification by using the IM axis, and the other is based on quantification in the retention time (RT) axis, where ions in the mobility dimension are summed.

**Data S2 | Set of true positives and true negatives for evaluating the MS-DIAL annotation pipeline.** Four data files exist, which include the set for electrospray ionization-positive-tandem mass spectrometry ESI(+)-MS/MS (DDA_Positive_True positive and negative spectral kit.txt), ESI(−)-MS/MS (DDA_Negative_True positive and negative spectral kit.txt), ESI(+)-ion mobility (IM)-MS/MS (PASEF_Positive_True positive and negative spectral kit.txt), and ESI(−)-IM-MS/MS (PASEF_Negative_True positive and negative spectral kit.txt).

### Supplementary Methods

**Consensus measurements in NIST SRM 1950 lipidomics data analysis**. The lipid extraction and LC-MS/MS analysis for NIST SRM 1950 plasma were performed for eight independent conditions (**Supplementary Note 1**). Briefly, a single-phase lipid extraction method (chloroform:methanol:water, 1:2:0.2, v/v/v)[21] was utilized in five conditions. The modified Matyash method using methanol and methyl *tert*-butyl ether was used in two conditions[22], and the modified Bligh and Dyer method (chloroform:methanol:water, 80:45:29, v/v/v) was used in one condition. TripleTOF 6600 SWATH-MS (SCIEX), TripleTOF 6600 DDA (SCIEX), timsTOF PASEF (Bruker), timsTOF DDA (Bruker), Q Exactive Plus DDA (Thermo Fisher), XevoG2 QTOF DDA (Waters), and 6546 QTOF DDA (Agilent) instruments were incorporated in this study. Various types of internal standards (ISs) such as SPLASH LIPIDOMICS I (Avanti), EquiSPLASH (Avanti), or a mixture of odd-chain and deuterium lipids were incorporated.

All data were processed using MS-DIAL 4, and the concentrations (pmol/μL plasma) were calculated by using ISs where the tag of level 2 was assigned if the lipid was quantified by an IS of the same lipid polar head class, and the tag of level 3 was assigned if the lipid was quantified by an IS of a similar lipid class or representative standard compound (https://lipidomics-standards-initiative.org/). The consensus values for lipid concentrations quantified as level 2 were measured using the sample coefficient of dispersion (COD) method as utilized in a previous lipidomics harmonizing study[11]. After the median value of laboratory mean (MEDM) was calculated for each lipid species, the COD value was calculated according to:

$$100 \times u/\mathrm{MEDM} \quad (1)$$

where $u$ denotes an associated standard uncertainty calculated as follows:

$$u = \sqrt{(\pi/2m)} \times 1.483 \times \mathrm{MAD} \quad (2)$$

where MAD and $m$ denote the median absolute deviation of the laboratory means and the number of laboratories, respectively. For evaluation purposes, the MEDMs were deemed acceptable for quality control activities if they had a COD value <40%. The estimations were only performed for lipids annotated by at least five laboratories.

**Prediction for retention time (RT) and collision cross-section (CCS) using machine learning**. The machine learning and evaluation for RT and CCS predictions were performed for lipid structures in which molecular species-level annotation was achieved. Because the double bond positions, *E/Z* isomers, and acyl chain positions cannot be determined in the majority of lipids, the SMILES code for each acyl chain moiety was represented by a single fatty acid-, alkyl chain-, or sphingobase structure which is considered as the majority in mammalian cells. The RT and lipid structure pairs of a total of 4,303 molecules of 108 lipid subclasses acquired with the Waters UPLC system coupled with SCIEX TripleTOF were utilized as the training (80%) or test set (20%) (**Supplementary Table 6**). The pairs of 3,079 total lipids acquired with Bruker Elute UHPLC coupled with timsTOF were used as an external validation set. The CCS and lipid structure pairs of the 3,570 total lipids acquired with Bruker timsTOF Pro were utilized as a training (80%) or test set (20%) (**Supplementary Table 7**). The chemical descriptor was generated using PaDEL version 2.21[23]. Overall, 152 and 328 of the descriptors exhibiting correlation coefficients >0.6 with the experimental RTs and CCSs in the training set were used for subsequent machine learning techniques.

RT and CCS prediction for lipidomics data were calculated using the Retip program, freely available at https://github.com/PaoloBnn/Retip/. The experimental dataset was divided into training and testing sets with a ratio of 80/20 based on the XLogP value. Retip provides five different machine learning methods:

XGBoost, Keras, LightGBM, Random Forest, and Bayesian Regularized Neural Network (BRNN). XGBoost machine included in Retip uses the grid search parameters with Caret (http://topepo.github.io/caret/index.html) with 10-fold cross validation resampling. Grid search allows searching the best performing tuning parameters, trying different combination of values. In this case, Retip uses nrounds, max_depth, eta, gamma, colsample_bytree, subsample, and min_child_weight to build 56 different machine learning models. For BRNN, our code automatically searches the optimal number of neurons to build the neural network from 1 to 7, in order to achieve the best results. All other tuning parameters were kept as default values. Our code for LightGBM included a 10-times cross-validation to select the best interaction to build the final model. Early stopping rounds, max_depth, and maximum leaf number were tuned to achieve the best results. Keras was optimized in several ways. In the first instance, dense_units were settled on the same value of column number of input data. Early stopping rounds were defined to avoid overfitting. Learning rate was tuned to a fixed value that yielded the best results in our tests. Our code built three different Keras models with different dropout values and automatically selected the best performing model based on the mean absolute error on the testing dataset. For random forest, our algorithm randomly assigned ten different values of mtry (number of variables available for splitting at each tree node). The best model after ten-time cross-validation was stored.

For RT prediction, the best machine learning was obtained using XGBoost with a root mean squared error (RMSE) of 0.19 min on training and 0.20 min on testing datasets. The most predictive chemical descriptor was XLogP, which is consistent with previously reported data[24]. CCS prediction was achieved by modifying the Retip source code to recognize the CCS column in the experimental data rather than RT. XGBoost also constituted a best-performing algorithm, with an RMSE of 1.4 $Å^2$ on training and 2.8 $Å^2$ on testing datasets. Two chemical descriptors, ATS3v and ATS2v, were almost equally important to build our prediction models.

Notably, although this study focused on lipidomics workflow, it also provided a CCS library for hydrophilic metabolites to render our MS-DIAL 4 universal (http://prime.psc.riken.jp/). The 2,646 structure and CCS pairs were utilized in machine learning, and a support vector regression algorithm was utilized for prediction with the automatically optimized parameters incorporating a recursive feature elimination algorithm. The prediction function is open-access within the AllCCS web server (http://allccs.zhulab.cn/). Predicted RT and CCS values for all small molecules were available at the MS-DIAL section of RIKEN PRIMe website (http://prime.psc.riken.jp/).

**Data format construction for ion mobility tandem mass spectrometry (IM-MS/MS) data**

The ion mobility base framework (IBF) was designed for the rapid retrieval for IM-MS/MS data because (A) the file reader using the Bruker software development kit (SDK) for ~1 GB file size of PASEF data requires several hours for retrieving all mass spectral data, which it was also true in ProteoWizard msconvert[25] (**Supplementary Fig. 8**), and (B) the file size of mzML becomes 10–100 times larger than the original file size. In addition to the original raw spectral data, the IBF format prepares the accumulated MS1 ion signals, summing the mass spectral data of the IM axis at the same RT bin, which is used for peak picking in the extracted ion chromatogram in the RT dimension. The data-dependent MS2 spectra in the same PASEF cycle are also accumulated and stored as the accumulated MS2 ion signals to be used in the usual spectral annotation. The *m/z* binning for the accumulation was set to three decimals for QTOF instrumental setting. All spectral data were indexed, as seek pointers, at the header space of IBF so that the application could rapidly retrieve the MS1 spectra for the peak detections in both RT and IM dimensions and assign the MS2 spectrum for detected peaks in the IM axis. The IBF file converter and source code are available at RIKEN PRIMe (http://prime.psc.riken.jp/), which currently supports the conversion of mzML, Waters (.raw), Agilent (.d), and Bruker (.tdf) format files. For 1 GB PASEF data, the IBF file size becomes 1 GB with 30 min conversion time. The IBF format file requires 10 s to retrieve all spectral data, which should be reasonable for the usual data processing workflow such as peak picking and alignment and for browsing raw data files in the application.

**Peak picking for IM-MS data**. The basic concept of peak picking, also termed as peak detection and feature detection, for IM data is to firstly detect the peaks in the RT and *m/z* dimension followed by peak picking in the IM axis after the spectra in the IM axis are accumulated within the RT peak width (**Supplementary Fig. 9**). Namely, each peak in the RT axis at a certain *m/z* value should have more than one peak in the IM axis. In this study, the peak feature in the RT and IM axis is termed as the "RT peak" and "IM peak", respectively.

The basic peak detection procedure of MS-DIAL[26] was used for the accumulated MS1 spectra in the IBF format file. In our reverse-phase LC-MS workflow, the RT peak width basically ranged from 10 to 30 s, and 20–60 data points were obtained in the PASEF acquisition because the cycle time was approximately 0.5 s. For each RT peak, the extracted ion "mobilogram" (EIM) was constructed in the IM axis by accumulating ions within the RT peak width, and the same peak detection procedure is applied to the EIM with the optimal parameter sets for the IM peak width scan points: in our condition, each IM peak had 10–50 data points, indicating a peak width of several dozen milliseconds. The MS/MS spectrum having the largest ion abundance within the IM peak width was assigned. Note that the accumulated MS/MS spectrum described in the previous section was assigned according to the information of RT, mobility unit (1/K0, K0, or milliseconds), and precursor *m/z* value. Here, the peak characteristics such as monoisotopic ion, isotopic ions, adduct type, and in-source fragment ions were determined in the peak feature information of the RT and *m/z* axis as described previously[26], and the IM peak characteristics followed the characteristics of the parent RT peak.

For data independent MS/MS acquisition (IM-DIA) data currently used in Waters and Agilent IM instruments, our MS2Dec algorithm[22,26] was renovated for chromatogram deconvolution for IM-DIA. After the centroided DIA-MS2 spectrum at a certain IM peak is extracted, the MS/MS mobilograms for the existing *m/z* values are constructed within the IM peak top ± 1.5 × IM peak width. The mobilograms are dedicated to the MS2Dec algorithm, and the deconvoluted MS/MS spectrum is obtained for each IM peak spot. The procedure and practical algorithm were optimized by testing several Agilent and Waters LC-IM-DIA-MS data sets from our colleagues.

**Peak alignment for IM-MS data**. The concept of the MS-DIAL peak alignment algorithm described in previous studies[22,26] was extended to the IM data architecture. Namely, the procedure contains four steps (1) creating a master peak list, (2) aligning each peak of each sample to the master peak list, (3) gap-filling for missing value interpolation, and (4) filtering based on the peak information in blank samples and the variation in the same sample group.

The initial template for the master peak list was constructed using the RT and IM peak features of the user-defined reference file. To encompass all RT and IM peak features involved to all analysis files, the master peak list was extended by the following procedures: (1) the RT peak feature of a biological sample is added to the master list if the feature does not exist in the current master peak list; and (2) the IM peak feature of a biological sample is added to the master list if the feature does not exist in the current master peak list. The existence was evaluated according to the following simple criteria: if $|RT_{master} – RT_{sample}| <$ RT tolerance is satisfied, the $RT_{sample}$ peak is recognized as "existing"; and if $|RT_{master} – RT_{sample}| <$ RT tolerance and $|IM_{master} – IM_{sample}| <$ IM tolerance are satisfied, the $IM_{sample}$ peak is recognized as "existing", where $RT_{master}$ and $RT_{sample}$ indicate the retention times of RT peaks of the master peak list and the biological sample, respectively, $IM_{master}$ and $IM_{sample}$ indicate the drift times of IM peaks of the master peak list and the biological sample, respectively, and the RT and IM tolerances indicate the user-defined tolerances of RT and drift time, respectively. The above procedure was repeated for all analysis files except for the analysis file used as the reference file; the parameter sets of 0.1 min and 0.05 1/k0 were used in this study for RT- and IM tolerances.

The peak features of each analysis file were aligned to the master peak list. The alignment was based on the matched score according to the following equation.

$$\text{Score} = m/z_{\text{factor}} \times |m/z_{\text{master}} - m/z_{\text{sample}}| + \text{RT}_{\text{factor}} \times |\text{RT}_{\text{master}} - \text{RT}_{\text{sample}}| + \text{IM}_{\text{factor}} \times |\text{IM}_{\text{master}} - \text{IM}_{\text{sample}}| \quad (3)$$

The coefficient values of $m/z_{\text{factor}}$, $\text{RT}_{\text{factor}}$, and $\text{IM}_{\text{factor}}$ were set to 1 in this study. $m/z_{\text{master}}$ and $m/z_{\text{sample}}$ indicate the *m/z* values of the master peak list and the biological sample, and the others are the same as described above. Note that the MS-DIAL program retains the information of RT peak properties; therefore, users can browse both results in RT- and IM dimensions.

The basic process for the gap-filling function to interpolate missing values in alignment results is the same as a previously described algorithm for LC-MS data processing[22,26]. A concern in IM-MS analysis is that the IM peak height and area values depend on the RT peak width: the RT peak width for accumulating ions to construct the extracted ion mobilogram can differ among samples, resulting in the over/under-estimation of IM peak height/area in an alignment result. This issue is especially important when some of the samples have a pure singlet peak and others exhibit a shoulder peak from co-eluted metabolites. Therefore, the current MS-DIAL program provides a user-defined parameter, “accumulated RT width”, to fix the RT peak range for constructing the extracted ion mobilogram. The RT tolerance, *m/z* tolerance, IM tolerance, and accumulated RT width for peak alignment in this study were set to ± 0.05 min, ± 0.015 Da, ± 0.05 1/K0, and ± 0.1 min, respectively.

No additional filtering option was used in this study although the current MS-DIAL program provides several filtering methods for IM data in addition to conventional LC-MS data to curate the alignment results automatically: details are provided in our tutorial (https://mtbinfo-team.github.io/mtbinfo.github.io/).

**Lipid tandem mass spectral atlas**. Although numerous software programs support lipid MS/MS annotations with the unique scoring systems, all programs require the knowledge of mass fragmentations for each lipid subclass. Through the analysis of our five-year untargeted lipidomics studies (1,056 LC-MS/MS data files from various biological samples), the mass fragmentations for a total of 117 lipid subclasses were formulated for their major adduct types including $[M+H]^+$, $[M+Na]^+$, $[M+NH_4]^+$, $[M-H_2O+H]^+$, $[M-H]^-$, $[M+HCOO]^-$, and $[M+CH_3COO]^-$ and a rule-based annotation system (decision tree algorithm) was constructed for each lipid ionization form (**Supplementary Table 2**): Note that the previous MS-DIAL series did not use a rule-based system for lipid annotations; this is thus the first study to provide a decision tree algorithm for annotating MS/MS spectra of 117 lipid subclasses. Although MS-DIAL contains the *in silico* spectral information for an additional 26 lipid subclasses such as LipidA, bile acid esters, and acylglycosylsterols based on the literatures, the present study describes lipid subclasses that were formulated using the experimental MS/MS spectral evidence from our biological studies in which the peak abundance of product ions was also incorporated into our annotation pipeline. The lipid nomenclature was proposed as the previously study[10] and in agreement with lipidomics standard initiative (LSI) committee, and is summarized in **Supplementary Table 2**. The lipid classification followed the classification system in LipidMAPS[9]. Notably, several lipid subclasses including mono- and digalactosylalkylacylglycerol (EtherMGDG, EtherDGDG), acyldiacylglycerylglucuronide (ADGGA), phosphatidylmethanol (PMeOH), phosphatidylethanol (PEtOH), ceramides incorporating sphinganine/sphingosine and β-hydroxy fatty acid (Cer-BDS, Cer-BS), acylhexosylceramide (AHexCer), hexosylceramide incorporating esterified fatty acid (HexCer-EOS), acylated sphingomyelin (ASM), sulfonolipid (SL), bile acid esters such as deoxycholic acid esters (DCAE), and several acylsterylglycosides (AHexST) such as AHexBRS (acylhexosylbrassicasterol) are not registered in the current LipidMAPS database (as of December, 2019). In addition, it includes several lipid subclasses such as *N*-acyl glycine (NAGly), *N*-acyl glycyl serine (NAGlySer), *N*-acyl ornithine (NAOrn), SL, AHexST, and various types of ether-linked phospholipids (Ether phosphatidylglycerol, EtherPS, EtherPI) that have never been incorporated in untargeted lipidomics studies. The classification and nomenclature of these lipid subclasses will be further discussed in the LSI consortium in the future.

Our workflow annotates the pair of lipid subclass and acyl chain compositions where positional isomers (*sn1*/*sn2*/*sn3*/*sn4*), *E/Z* isomers, and double bond positions are not determined basically: the ion abundance rules to distinguish positional isomers as described in previous studies[27,28] can be added in our open source code if needed although we believe that the universal rule for the stereoisomer clarifications is unavailable in conventional low-energy collision induced dissociation (CID)-based mass fragmentations. In fact, several acyl chain positional isomers e.g. in HBMP and ADGGA can be determined by the low energy-CID, and the position is specified by the virgule "/" character instead of underline "_". The MS raw data for sharing the lipid MS/MS spectra is available at RIKEN DropMet (http://prime.psc.riken.jp/menta.cgi/prime/drop_index) by the indexes of DM0022, DM0030, and DM0031 and the details are also described in **Supplementary Table 1** and **Supplementary Note 1**.

Three confirmation levels exist for lipid structure and MS/MS annotations: level 1, for which 51 lipid subclasses were confirmed using the authentic standard compounds; level 2, with 60 lipid subclasses elucidated according to the fragment evidence combined with literature showing the lipid structure and MS/MS spectra (total 20 articles are cited in **Supplementary Note 2**); and level 3, comprising six lipid subclass structures that have not been reported elsewhere and were putatively assigned by untangling the fragment ions in positive and negative ion modes. For level 3 annotations, several forms of indirect evidence were accumulated. First, the lipid ions were confirmed in both positive and negative ion modes to exclude the possibility of in source fragment ions. Second, the fragment ions were reasonably formulated according to the hydrogen rearrangement rules[29] and the confirmation of polar head-specific ions in both positive and negative ion modes. Finally, several acyl chain varieties of the lipid subclass were detected in the biological samples. The mass fragmentation of 117 lipid subclasses was formulated in our annotation pipeline; the summary is described in **Supplementary Note 2**, **Supplementary Figs. 2–7**, and **Supplementary Table 2**.

**Quantification, nomenclature, and shorthand notation for lipids**. The lipid quantification followed the definition of the LSI: level 2 quantification was achieved if the lipid molecule was quantified by the IS compound of the same lipid subclass and level 3 quantification was achieved if the lipid molecule was quantified by the IS compound of a similar or representative lipid class category. The peak quantification of lipids to create our lipidomics database was performed according to the representative adduct type in each lipid subclass. For example, the ceramide containing sphingosine and nonhydrolyzed acyl chain, termed as Cer-NS, was quantified by $[M+CH_3COO]^-$ in negative ion mode although the molecule can also be characterized as $[M+H]^+$ and $[M-H]^-$ in MS-DIAL 4. The adduct type for lipid quantification was determined by the ion sensitivity and the practical annotation handiness. Note that the description for MS/MS spectrum annotation for the acetate adduct form could be replaced by the formate adduct form. The only difference of mass spectra between acetate and formate adduct forms is the *m/z* value of the precursor ion derived from the $CH_2$ (14.016 Da) difference, whereas the MS/MS pattern of product ions when using ammonium acetate as the LC solvent is almost equivalent as that obtained using ammonium formate according to our experience.

The acyl chain moiety is characterized by the carbon and double bond numbers, such as 18:2 indicating 18 carbon and 2 double bonds in the acyl chain. The number of oxygen atoms involved with acyl chain incorporation; e.g., in oxidized phospholipids and ceramides, is separated by a semi-colon: e.g., Cer 18:1;O2/18:0;O, which is usually described in mammalian studies as Cer d18:1/18:0h, incorporating sphingosine and hydroxy fatty acid in the acyl chain moiety. When the hydroxy position can be determined from the MS/MS information, the description becomes Cer 18:1;(1OH, 3OH)/18:0;(2OH) which is known as Cer-AS in mammalian ceramides[18]. The ether- and vinyl ether linkages are described by O-18:0 and P-18:0, respectively. The *N*-acyl linkage is described by N-18:0 except for ceramide species. Underlining "_" is used to describe acyl chain compositions if the *sn1*, *sn2*, *sn3*, and *sn4* positional isomers are uncharacterized whereas the virgule (/) character is used if the acyl chain position is determined in a specific position such as *N*-acyl chain, ether chain, or acyl sugar. The details are described in **Supplementary Table 2 and Supplementary Note 2** and all lipidomics results can be downloaded as **Supplementary Data 1** and at the RIKEN PRIMe website (http://prime.psc.riken.jp/).

**Program comparisons**. The various lipidomics software programs including the previous version of MS-DIAL (version 3) were evaluated as the benchmark for MS-DIAL 4 (**Table 1** and **Supplementary Table S3**). We selected the benchmark programs according to the following two criteria: the software tools were (1) designed for untargeted lipidomics data using LC-MS/MS and (2) introduced in LipidMAPS or LSI websites. Among these, we excluded LPPtiger[30], LIMSA[31], LipidQA[32], LipidMS[33], and LDA[27] for the evaluations because LPPtiger supports only oxidized lipids, LIMSA has not been updated since 2006, LipidQA is not maintained, LipidMS is only for data independent MS/MS acquisition, and LDA is designed for targeted lipidomics.

MS-DIAL 4, MS-DIAL 3[6], LipiDex[34], LipidMatch[35] (2.0.2), LIQUID[36] (7.4.6988), LipidHunter2[37], Greazy[38], and MzMine 2[39] (2.51) were evaluated. First, general utility features including the function support for (1) peak detection/quantification, (2) peak alignment/data integration, (3) MS/MS-based annotation, (4) decision tree annotation for providing appropriate structure representation of lipids by fragment evidences, and (5) supported lipid subclasses by the default library were evaluated. Secondly, the LC-MS/MS data of a dilution series of standard mixtures containing 22 different lipid subclass molecules including FA 20:4, CAR 18:1, LPA 18:1, LPC 18:1, LPE 18:1, LPG 18:1, LPI 18:1, LPS 18:1, PA 18:1_18:1, PC 18:1_18:1, PC O-18:1_18:1, PE 18:1_18:1, PE P-18:0_18:1, PG 18:1_18:1, PS 18:1_18:1, BMP 18:1_18:1, SPB 18:1;2O, Cer 18:1;2O/18:1, MG 16:1, DG 18:1_18:1, TG 18:1_18:1_18:1, and CE 17:0 were evaluated. The concentration of all standards was adjusted equally in the mixture, and the dilution series was established as 5 nM, 10 nM, 25 nM, 50 nM, 100 nM, 250 nM, 500 nM, 1 μM, 2.5 μM, 5 μM, 10 μM, 25 μM, and 50 μM in an LC-MS vial. The injection volume was set to 1 μL for both positive and negative ion modes, and the LC-MS/MS setting of method 2 described in **Supplementary Note 1** was used for data acquisition. The parameter setting for each software program was optimized so that the lipid subclasses were annotated to the greatest degree possible; the parameter details are described in **Supplementary Table 3**. If the lipid molecule was annotated, the term of "ok (with a brief summary of the parameter condition)" was assigned. If not, the term of "NA" (not assigned) was added. Finally, the annotation results for NIST SRM 1950 plasma analyzed by method 2 were compared, and the FDR value of our method can be estimated as less than 2.5% although they could not be evaluated for other programs. To count named lipids, the duplicate names and duplicate peak indexes were excluded where the peak information having the highest annotation score among lipid molecules assigned to the same peak index was kept.

To investigate the coverage of lipids containing VLC-PUFAs (**Supplementary Fig. 11**), our lipidomics data of eye and testis tissues were processed by their programs with the default MS/MS libraries. Because such software programs can import customized rules or spectral libraries, the result does not directory reflect the program functionalities for annotation.

#### Reference (Supplementary Methods)

### Supplementary Note 1 | Details of the biological samples and analytical platforms

A total of 41 unique tissues/species which contain human plasma, 19 different mouse tissues, 9 algal species, and 3 plant species from 10 different analytical platforms were processed by MS-DIAL 4.0. The detail of biological samples and internal standards were described in **Supplementary Table S1** and **S2**.

#### LC-MS/MS and LC-IM-MS/MS methods

*Method 1: SCIEX TripleTOF DDA high mass range method*

Methanol, isopropanol, and acetonitrile of LC-MS grade were purchased from Wako. Ammonium acetate and EDTA were purchased from Wako and Dojindo, respectively. Milli-Q water was purchased from Millipore. SPLASH Lipidomics I or EquiSPLASH used as internal standards were purchased from Avanti Polar Lipids.

The LC system consisted of a Waters Acquity UPLC system. Lipids were separated on an Acquity UPLC Peptide BEH C18 column (50 × 2.1 mm; 1.7 µm) (Waters, Milford, MA, USA). The column was maintained at 45°C at a flow-rate of 0.3 mL/min. The mobile phases consisted of (A) 1:1:3 (v/v/v) acetonitrile:methanol:water with ammonium formate (5 mM) and 10 nM EDTA and (B) 100% isopropanol with ammonium formate (5 mM) and 10 nM EDTA. A sample volume of 0.5−3 µL, which depended on biological samples, was used for the injection. The separation was conducted under the following gradient: 0 min 0% (B); 1 min 0% (B); 5 min 40% (B); 7.5 min 64% (B); 12 min 64% (B); 12.5 min 82.5% (B); 19 min 85% (B); 20 min 95% (B); 20.1 min 0% (B); and 25 min 0% (B). Sample temperature was maintained at 4°C.

Mass spectrometric detection of lipids was performed on a quadrupole/time-of-flight mass spectrometer TripleTOF 5600 or 6600 (SCIEX, Framingham, MA, USA). All analyses were performed at the high resolution mode in MS1 (~35,000 full width at half maximum (FWHM)) and at the high sensitivity mode (~20,000 FWHM) in MS2. Data dependent MS/MS acquisition (DDA) was used. The parameters were MS1 and MS2 mass ranges, *m*/*z* 70–1750; MS1 accumulation time, 200 ms; MS2 accumulation time, 70 ms; collision energy, +40/–42 eV; collision energy spread, 15 eV; cycle time, 1370 ms; curtain gas, 30; ion source gas 1, 40(+)/50(–); ion source gas 2, 80(+)/50(–); temperature, 250°C(+)/300°C(-); ion spray voltage floating, +5.5/–4.5 kV; declustering potential, 80 V. The other DDA parameters were dependent product ion scan number, 16; intensity threshold, 100 cps; exclusion time of precursor ion, 0s; mass tolerance, 20 ppm; ignore peaks, within *m/z* 200; and dynamic background subtraction, True. The mass calibration was automatically performed using an APCI positive/negative calibration solution via a calibration delivery system (CDS).

*Method 2: SCIEX TripleTOF DDA normal mass range method*

The LC-MS machine platform was the same as Method 1, and the LC condition was also the same as Method 1. The mass spectrometer settings were changed for scanning *m*/*z* 70–1250 mass range accordingly: MS1 and MS2 mass ranges, *m*/*z* 70–1250; MS1 accumulation time, 250 ms; MS2 accumulation time, 100 ms; cycle time, 1300 ms. The other parameters were the same as Method 1.

*Method 3: Bruker timsTOF Pro DDA for plant lipidomics*

Methanol, isopropanol, and acetonitrile of LC-MS grade were purchased from Wako. Ammonium formate and formic acid were purchased from Wako. Water was purchased from Millipore. The LC system consisted of a Bruker Elute UHPLC system. Lipids were separated on an Acquity UPLC HSS T3 C18 column (50 × 1.0 mm; 1.8 µm) (Waters, Milford, MA, USA). The column was maintained at 55°C at a flow-rate of 0.15 mL/min. The mobile phases consisted of (A) 200:800:10:1 (v/v/v/v) acetonitrile:water:1M ammonium formate:formic acid and (B) 100:900:10:1 (v/v/v/v) acetonitrile:isopropanol:1M ammonium formate:formic acid. A sample volume of 2 μL was used for the injection. The separation was conducted under the following gradient: 0 min 35% (B); 3 min 70% (B); 7 min 85% (B); 10 min 90% (B); 12 min 90% (B); 12.5 min 35% (B); and 15 min 35% (B). Sample temperature was maintained at 10°C.

Mass spectrometric detection of lipids was performed on a hybrid trapped ion mobility-quadrupole time-of-flight mass spectrometer (timsTOF Pro, Bruker Daltonics, Bremen, Germany). Data dependent MS/MS acquisition (DDA) was used. The parameters were MS1 mass ranges, *m*/*z* 200–1600 and MS2 mass ranges, *m*/*z* 50–1600; MS1 cycle time time, 0.5 sec; MS2 accumulation, 14 Hz ; collision energy, 30 eV; end plate offset, 500 V; capillary voltage, +4.2/–4.2 kV; nebulizer pressure, 2 bar; dry gas, 8.0 l/min; dry temperature, 200°C; funnel 1 RF (radio frequency), 300 Vpp; funnel 2 RF, 250 Vpp; multipole RF, 200 Vpp; deflection delta, 70 V; quadrupole ion energy, 5 eV; collision transfer energy, 8 eV; collision RF, 1100 Vpp; transfer time, 54 μs; pre-pulse Storage, 5 μs; intensity threshold, 31cts.; exclusion time of precursor ion, 0.2 min. The mass calibration was automatically performed using 5 mM sodium formate calibration solution.

*Method 4: Bruker timsTOF Pro PASEF for plant lipidomics*

The LC-MS machine platform was the same as Method 3, and the regents and LC conditions were the same as Method 3. The PASEF mode acquisition, i.e. data dependent MS/MS acquisition with the ion mobility separation, was performed by the following parameters. The parameters were MS1 mass ranges, *m/z* 200–1600 and MS2 mass ranges, *m/z* 50–1600; MS1 and MS2 accumulation time, 148.5 ms; mobility range 0.55–1.9; collision energy 30 eV; end plate offset, 500 V; capillary voltage, +4.2/–4.2 kV; nebulizer pressure, 2 bar; dry gas, 8.0 l/min; dry temperature, 200°C; funnel 1 RF (radio frequency), 300 Vpp; funnel 2 RF, 250 Vpp; multipole RF, 200 Vpp; deflection delta, 70 V; quadrupole ion energy, 5 eV; collision transfer energy, 15 eV; collision RF, 1100 Vpp; transfer time 54 μs; pre-pulse storage, 5 μs; tims transfer Δ1, –20 V/+20 V; tims transfer Δ2, -120 V/+120 V; tims transfer Δ3, +70 V/–70 V; tims transfer Δ4, +100 V/-100 V; tims transfer Δ5, 0 V; tims transfer Δ6, +100 V/–100 V; intensity threshold, 2500 cts; target intensity, 20000 cts; exclusion time of precursor ion, 0.1 min. The mass calibration was automatically performed using 5 mM sodium formate calibration solution.

*Method 5: Bruker timsTOF Pro DDA for mouse tissue lipidomics*

Methanol, isopropanol, and acetonitrile of LC-MS grade were purchased from Wako. Ammonium acetate and EDTA were purchased from Wako and Dojindo, respectively. Milli-Q water was purchased from Millipore. The LC-MS machine system was the same as Method 3 and 4, and the LC condition was the same as Method 1. Data dependent MS/MS acquisition (DDA) was used. The parameters were MS1 mass ranges, *m/z* 200–2500 and MS2 mass ranges, *m/z* 50–2500; MS1 cycle time, 0.5 sec; MS2 accumulation 14 Hz ; collision energy, 30 eV; end plate offset, 500 V; capillary voltage, +4.5 kV/–4.2 kV; nebulizer pressure, 2 bar; dry gas, 10.0 l/min; dry temperature, 220°C/200°C; funnel 1 RF, 300 Vpp; funnel 2 RF, 200 Vpp/250 Vpp; multipole RF, 200 Vpp; deflection delta, +70 V/–70 V; quadrupole ion energy, 6 eV/5 eV; collision transfer energy, 14 eV/15 eV; collision RF, 1100 to 1800 Vpp stepping; transfer time 45 to 75 μs stepping; pre-pulse storage, 10 μs/5 μs; intensity threshold, 31cts.; exclusion time of precursor ion, 0.2 min. The mass calibration was automatically performed using 5 mM sodium acetate calibration solution.

*Method 6: Bruker timsTOF Pro PASEF for mouse tissue lipidomics*

The regents and LC conditions were the same as Method 5. Data dependent MS/MS acquisition (DDA) was used. The parameters were MS1 and MS2 mass ranges, *m/z* 50–2500; MS1 and MS2 accumulation time, 100 ms; mobility range, 0.55–1.90; collision energy 30 eV; end plate offset, 500 V; capillary voltage, +4.5 kV/–4.2 kV; nebulizer pressure, 2 bar; dry gas, 10.0 L/min; dry temperature, 220°C; funnel 1 RF, 300 Vpp; funnel 2 RF, 200 Vpp/250 Vpp; multipole RF, 200 Vpp; deflection delta, 70 V/–70 V; quadrupole ion energy, 6 eV/5 eV; collision transfer energy, 14 eV/15 eV; collision RF, 1500 Vpp/1100 Vpp; transfer time 75 μs/54 μs; pre-pulse storage, 10 μs/5 μs; tims transfer $\Delta$1, –20 V/20 V; tims transfer $\Delta$2, –120 V/120 V; tims transfer $\Delta$3, 70 V/–70 V; tims transfer $\Delta$4, 100 V/–100 V; tims transfer $\Delta$5, 0 V; tims transfer $\Delta$6, 100 V/–100 V; intensity threshold, 250 cts.; target intensity, 4000 cts; exclusion time of precursor ion, 0.1 min. The mass calibration was automatically performed using 5 mM sodium acetate calibration solution. For the confirmation of ESI(+)-MS/MS spectra for phosphatidylcholine, the positive ion mode data was acquired with the following setting. The parameters were dry temperature, 200°C; funnel 2 RF, 250 Vpp; quadrupole ion energy, 5 eV; collision transfer energy, 15 eV; collision RF, 1100 Vpp; and transfer time 54 μs; pre-pulse storage, 5 μs. The other parameters were the same as above.

*Method 7: Waters XevoG2 QTOF DDA for plant lipidomics*

Methanol, isopropanol, and acetonitrile of LC-MS grade were purchased from Wako. Ammonium acetate, formic acid, methyl *tert*-butyl ether (MTBE), and water were also purchased from Wako. 1,2-Didecanoyl-*sn*-glycero-3-phosphocholine used for the internal standard was purchased from Sigma-Aldrich.

The LC system consisted of a Waters Acquity UPLC system. Lipids were separated on the mostly same condition as described in Method 3 except for using ammonium acetate instead of ammonium formate. The LC system consisted of a Waters Acquity UPLC system. Lipids were separated on an Acquity UPLC HSS T3 C18 column (50 × 1.0 mm; 1.8 μm) (Waters, Milford, MA, USA). The column was maintained at 55°C at a flow-rate of 0.15 mL/min. The mobile phases consisted of (A) 200:800:10:1 (v/v/v/v) acetonitrile:water:1 M ammonium acetate:formic acid and (B) 100:900:10:1 (v/v/v/v) acetonitrile:isopropanol:1 M ammonium acetate:formic acid. A sample volume of 1 µL was used for the injection. The separation was conducted under the following gradient: 0 min 35% (B); 3 min 70% (B); 7 min 85% (B); 10 min 90% (B); 12 min 90% (B); 12.5 min 35% (B); and 15 min 35% (B). Sample temperature was maintained at 10°C.

Mass spectrometric detection of lipids was performed on a quadrupole/time-of-flight mass spectrometer Xevo G2 QTOF MS (Waters, Milford, MA, USA). MS2 analyses were performed at the sensitivity mode. Data dependent MS/MS acquisition (DDA) was used for MS2. The conditions for recording were as follows: source capillary, 3.0 (positive ion mode) and 2.5 kV (negative ion mode); sampling cone, 20 (positive) and 40 (negative); extraction cone, 4.0; source temperature, 120°C; desolvation temperature, 450°C; cone gas flow, 50 L/h; desolvation gas flow, 600 L/h; scan ranges, *m*/*z* 100–1600; MS1 scan time, 200 ms (centroid); MS2 scan time, 100 ms (centroid); collision energy, 20–50 eV (ramp mode). The other DDA parameters were event number, 6; scan repeat, 3; peak selection mode to trigger MS/MS, intensity-based detection; deisotope peak detection, yes; ionization mode, ESI; and correction by lock mass function, yes.

*Method 8: SCIEX TripleTOF 6600 SWATH for NIST SRM 1950 human plasma*

Water, isopropanol, and acetonitrile were purchased from Fisher Optima. Methanol was purchased from J.T. Baker. Ammonium formate, ammonium acetate, formic acid, and methyl *tert*-butyl ether (MTBE) were purchased from Sigma–Aldrich. Odd chain and deuterated lipids used as internal standards were purchased from Avanti Polar Lipids, CDN Isotopes, Cayman Chemical, and Sigma–Aldrich.

The LC system consisted of an Agilent 1290 system (Agilent Technologies Inc.) with a pump (G4220A), a column oven (G1316C), and an autosampler (G4226A). Lipids were separated on an Acquity UPLC CSH C18 column (100 × 2.1 mm; 1.7 µm) coupled to an Acquity UPLC CSH C18 VanGuard precolumn (5 × 2.1 mm; 1.7 µm) (Waters, Milford, MA, USA). The column was maintained at 65°C at a flow-rate of 0.6 mL/min. For LC–ESI(+)-MS analysis the mobile phases consisted of (A) 60:40 (v/v) acetonitrile:water with ammonium formate (10 mM) and formic acid (0.1%) and (B) 90:10:0.1 (v/v/v) isopropanol:acetonitrile:water with ammonium formate (10 mM) and formic acid (0.1%). For LC–ESI(–)-MS analysis the organic solvents for mobile phases were the same with the exception of using ammonium acetate (10 mM) as mobile-phase modifier. A sample volume of 3 µL was used for the injection in both ESI(+) and ESI(–). The separation was conducted under the following gradient in ESI(+): 0 min 15% (B); 0−2 min 30% (B); 2−2.5 min 48% (B); 2.5−11 min 82% (B); 11−11.5 min 99% (B); 11.5−12 min 99% (B); 12−12.1 min 15% (B); and 12.1−15 min 15% (B). The separation was conducted under the following gradient in ESI(–): 0 min 15% (B); 0−2 min 30% (B); 2−2.5 min 48% (B); 2.5−9.5 min 76%; 9.5−9.6 min 99% (B); 9.6−10.5 min 99% (B); 10.5−10.6 min 15% (B); 10.6−13.5 min 15% (B). Sample temperature was maintained at 4°C.

Mass spectrometric detection of lipids was performed on a quadrupole/time-of-flight mass spectrometer TripleTOF 6600 (SCIEX, Framingham, MA, USA). All analyses were performed at the high resolution mode in MS1 (~35,000 full width at half maximum (FWHM)) and at the high sensitivity mode (~15,000 FWHM) in MS2. For ESI(+), the SWATH parameters were MS1 accumulation time, 100 ms; MS1 mass range, *m*/*z* 100–1700; MS2 accumulation time, 10 ms; collision energy, 45 eV; collision energy spread, 15 eV; cycle time, 550 ms; Q1 window, 20 Da; SWATH mass range, *m*/*z* 300–1100; number of SWATH experiments, 40; MS2 mass range: *m*/*z* 80–1100. Other parameters were curtain gas, 35; ion source gas 1, 60; ion source gas 2, 60; temperature, 350°C; ion spray voltage floating, 4.5 kV; declustering potential, 80 V. For ESI(–), the SWATH parameters were MS1 accumulation time, 100 ms; MS1 mass range, *m*/*z* 100–1700; MS2 accumulation time, 10 ms; collision energy, −50 eV; collision energy spread, 10 eV; cycle time, 550 ms; Q1 window, 15 Da; SWATH mass range, *m*/*z* 400–1000; number of SWATH experiments, 40; MS2 mass range: *m*/*z* 80–1000. Other parameters were curtain gas, 35; ion source gas 1, 60; ion source gas 2, 60; temperature, 350°C; ion spray voltage floating, −4.5 kV; declustering potential, −80 V. The mass calibration was automatically performed using an APCI positive/negative calibration solution via a calibration delivery system (CDS).

*Method 9: Thermo Q Exactive Plus DDA for NIST SRM 1950 human plasma*

Water was purchased from VWR International, isopropanol from Merck, and acetonitrile from Honeywell. MTBE, ammonium formate, ammonium acetate, formic acid, and acetic acid were purchased from Sigma–Aldrich. Odd chain and deuterated lipids used as internal standards were purchased from Avanti Polar Lipids, Chromsystems, Cayman Chemical, and Sigma–Aldrich.

The LC system consisted of a Vanquish UHPLC system (Thermo Fisher Scientific, Bremen, Germany) with a pump, a column oven and an autosampler. Lipids were separated on an Acquity UPLC BEH C18 column (50 × 2.1 mm; 1.7 µm) coupled to an Acquity UPLC BEH C18 VanGuard precolumn (5 × 2.1 mm; 1.7 µm) (Waters, Milford, MA, USA). The column was maintained at 65°C at a flow-rate of 0.6 mL/min. For LC–ESI(+)-MS analysis the mobile phases consisted of (A) 60:40 (v/v) acetonitrile:water with ammonium formate (10 mM) and formic acid (0.1%) and (B) 90:10:0.1 (v/v/v) isopropanol:acetonitrile:water with ammonium formate (10 mM) and formic acid (0.1%). For LC–ESI(–)-MS analysis the organic solvents for mobile phases were the same with the exception of using ammonium acetate (10 mM) and acetic acid (0.1%) as mobile-phase modifiers. A sample volume of 2 µL and 3 µL was used for the injection in ESI(+) and ESI(–), respectively. Separation was conducted under the following gradient for LC–ESI(+): 0 min 15% (B); 0–1 min 30% (B); 1–1.3 min from 30% to 48% (B); 1.3–5.5 min from 48% to 82% (B); 5.5–5.8 min from 82% to 99% (B); 5.8–6 min 99% (B); 6–6.1 min from 99% to 15% (B); 6.1–7.5 min 15% (B). For LC–ESI(–), the following gradient was used: 0 min 15%

(B); 0–1 min 30% (B); 1–1.3 min from 30% to 48% (B); 1.3–4.8 min from 48% to 76% (B); 4.8–4.9 min from 76% to 99% (B); 4.9–5.3 min 99% (B); 5.3–5.4 min from 99% to 15% (B); 5.4–6.8 min 15% (B). Sample temperature was maintained at 4°C.

Mass spectrometric detection of lipids was performed on a quadrupole/orbital ion trap mass spectrometer Q Exactive Plus with a HESI-II ion source (Thermo Fisher Scientific, Bremen, Germany). Simultaneous MS1 and MS/MS (data-dependent MS/MS) acquisition was used. The parameters were as follows: sheath gas pressure, 60; aux gas flow, 25; sweep gas flow, 2; spray voltage, 3.6 kV and −3.0 kV for ESI(+) and ESI(–), respectively; capillary temperature, 300°C; aux gas heater temperature, 370°C; MS1 mass range, *m*/*z* 200–1700; MS1 resolving power, 35,000 FWHM (*m*/*z* 200); number of data-dependent scans per cycle: 3; MS/MS resolving power, 17,500 FWHM (*m*/*z* 200); MS1 ion time: 100 ms; MS2 ion time: 50 ms; normalized collision energy, 20% for ESI(+) and 10, 20, 30% for ESI(–). The instrument was tuned using a Thermo positive and negative ion mode calibration solutions.

*Method 10: Agilent 6546 LC-QTOF/MS DDA for NIST SRM 1950 human plasma*

Ultrapure water was produced with a Milli-Q Integral system equipped with a LC-Pak Polisher and a 0.22 µm point-of-use membrane filter cartridge (EMD Millipore, Billerica, MA, USA). Ammonium fluoride and LC/MS grade ammonium acetate were purchased from Millipore Sigma (St. Louis, MO, USA). Isopropanol of LC/MS grade “LiChrosolv” was purchased from Sigma Aldrich. HPLC-grade acetonitrile and methanol were purchased from Honeywell (Morristown, NJ, USA). SPLASH lipidomics I used as internal standards were purchased from Avanti Polar Lipids. Chloroform was purchased from Honeywell (Charlotte, North Carolina).

The LC system consisted of an Agilent 1290 Infinity II (Agilent Technologies, Santa Clara, CA, USA) with a pump, a column oven and an autosampler. Lipids were separated on an Agilent InfinityLab

Poroshell 120 EC-C18 column (100 × 3.0 mm; 2.7 µm) coupled to an Agilent InfinityLab Poroshell 120 EC-C18 column precolumn (5 × 3.0 mm; 2.7 µm). The column was maintained at 50°C at a flow-rate of 0.6 mL/min. The mobile phases consisted of (A) 9:1 (v/v) water:methanol with ammonium acetate (10 mM) and ammonium fluoride (0.2 mM) and (B) 2:3:5 (v/v/v) acetonitrile:methanol/isopropanol with ammonium formate (10 mM) and ammonium fluoride (0.2 mM). A sample volume of 5 µL was used for tinjections in ESI(+) and ESI(–). Separation was conducted under the following gradient: 0 min 70% (B); 1 min 70% (B); 3.5 min 86% (B); 10.0 min 86% (B); 11.0 min 100% (B); 17.0 min 100% (B); 17.1 min 70% (B); 19.0 min 70% (B). Sample temperature was maintained at 4°C.

Mass spectrometric detection of lipids was performed on a quadrupole/time-of-flight mass spectrometer 6546 Q-TOF (Agilent Technologies, Santa Clara, CA, USA). Simultaneous MS1 and MS/MS (data-dependent MS/MS) acquisition was used. The parameters were as follows: gas temperature, 200°C; gas flow, 10 L/min; nebulizer (psig), 50; sheath gas temperature, 300°C; sheath gas flow, 12 L/min; VCap, 3.5 kV and −3.0 kV for ESI(+) and ESI(–), respectively; nozzle voltage, 0 V; fragmentor, 150 V; skimmer, 65 V; octupole RF Vpp, 750 V; MS1 and MS2 mass range, *m*/*z* 40–1700; min MS and MS/MS acquisition rate: 3 spectra/s; isolation width, narrow (~1.3 *m/z*); max precursors per cycle, 3; precursor abundance-based scan speed, target 25,000 counts/spectrum; use MS/MS accumulation time limit, yes; reject precursors that cannot reach target TIC, no; threshold for MS/MS, 5,000 counts and 0.001%; active exclusion enabled, one repeat, then exclude for 0.05 minutes; purity, stringency 70%, cut off 0%; isotope model, common organic molecules; sort precursors, 1,2,unknown; static exclusion ranges, *m/z* 40 to 151 for ESI(+) and *m/z* 40 to 210 for ESI(–); collision energy, 20 eV for ESI(+) and 25 eV for ESI(–). The instrument was tuned using a reference mass *m/z* 121.050873, *m/z* 1221.990637 (+) and *m/z* 119.03632, *m/z* 980.016375 (–).

**Biological resources**

*NIST SRM 1950 human plasma*

As the benchmark to prone the scalability of MS-DIAL 4.0, NIST SRM 1950 human plasma sample was analyzed by eight platforms (Method 1, 2, 5, 6, 7, 8, 9, 10), and the experimental protocols for each platform were described as follows.

For method 1, 2, 5, and 6, an aliquot of 20 μL of NIST SRM 1950 human plasma sample (National Institute of Standards and Technology) was added to 100 μL of ice cold chloroform and vortexed for 10 s. After 1-h incubation on ice, 200 μL of ice cold MeOH containing 5 μL of EquiSPLASH, 10 μM palmitic acid-$d_3$, and 10 μM stearic acid-$d_3$ was added and vortexed for 10 s. After 2-h incubation on ice, the solvent tube was centrifuged at 2000×*g* for 10 min at 4 °C. 200 μL of supernatant was transferred to LC-MS vial. For method 7, after the same procedure as described above is performed, the supernatant was dried with a vacuum dryer and resuspended in 200 μL of ethanol. The solvent was transferred to LC-MS vial.

For method 8, an aliquot of 20 μL of NIST SRM 1950 human plasma sample (National Institute of Standards and Technology) was added to 225 μL of ice cold MeOH and vortexed for 10 s. Then, 750 μL of ice cold MTBE was added and vortexed for 10 s. After shaking for 6 min at 4°C in the orbital mixer, 188 μL LC-MS-grade water was added and vortexed for 20 s. After centrifugation for 2 min at 14,000 rcf, 350 μL of the supernatant was transferred to a new 1.5 mL Eppendorf tube and evaporated to dryness in the Labconco Centrivap cold trap concentrator. The dried sample was resuspended in 110 μL MeOH:toluene 9:1 (v/v) containing CUDA (12-[[(cyclohexylamino)carbonyl]amino]-dodecanoic acid) internal standard (50 ng/mL). After vortexing for 20 s the samples were centrifuged for 2 min at 14,000 rcf and 100 μL of the supernatant was transferred to a glass amber vial with micro-insert.

For method 9, an aliquot of 25 μL of NIST SRM 1950 human plasma sample was added to a mixture of 165 μL of ice cold MeOH and 600 μL of ice cold MTBE. After shaking for 30 s in a cold block,

165 µL of 10% MeOH/90% water (v/v) mixture was added followed by shaking for 30 s. After centrifugation for 5 min at 16,000 rcf, 100 µL of the supernatant was transferred to a new 1.5 mL Eppendorf tube and evaporated to dryness in the Labconco Centrivap cold trap concentrator. The dried sample was resuspended in 100 µL MeOH containing CUDA internal standard (200 ng/mL). After shaking for 30 s the samples were centrifuged for 5 min at 16,000 rcf and 90 µL of the supernatant was transferred to a glass amber vial with micro-insert.

For method 10, 50 µL of NIST SRM 1950 human plasma sample was added to 50 µL SPLASH LIPIDOMIX I and 400 µL of ice cold MeOH. After vortexed briefly, then sonicated for five minutes, 800 µL of chloroform was added followed by shaking for 1 min. 240 µL of water was added for phase partitioning followed by shaking for 1 min. After centrifugation for 4 min at 16,000 g, the lower layer was carefully removed with a gas-tight glass syringe, and transferred to a 2 mL Agilent A-Line amber glass vial. To re-extract the remaining interphase and upper phase layers, 900 µL chloroform/methanol/water (86:14:1, v/v/v) was added, and the mixture was vortexed for one minute and centrifuged again. The combined lower layers from ten 50 µL extractions (ESI-) were combined, and the lower layers from a single 50 µL extraction (ESI+) were combined and dried by a vacuum concentrator. The dried samples were resuspended in 100 µL of a MeOH/chloroform mixture (9:1, v/v). After shaking for 1 min, the samples were centrifuged for 4 min at 16,000 g and 250 µL of the supernatants were transferred to glass amber vials with micro-inserts.

*Mouse tissues*

The adrenal gland, brain, brown adipose tissue, eye, feces, heart, kidney, large intestine, liver, lung, pancreas, skeletal muscle, skin (ear), small intestine, spleen, testis, and white adipose tissues of 8–10 weeks male mice (C57BL/6 J background, CLEA Japan, Tokyo, Japan) were harvested according to the ethical protocol approved by the RIKEN Center for Integrative Medical Sciences (2019-015(2)). The detail of diet and genetics background was described in **Supplementary Data 1**. Tissues were frozen immediately after dissection and stored at −80 °C until lipid extraction. The lipid extraction was performed according to the previously reported method[21,24] where the mixed solvent containing methanol, chloroform, and water (MeOH:$CHCl_3$:$H_2O$, 1:2:0.2, v/v/v) was utilized. The detail of solvent volumes and internal standards used in this study was described in **Supplementary Table S1 and S2**. The tissues were homogenized by a multi-beads shocker (YASUI KIKAI, Japan) with a metal cone (YASUI KIKAI, Japan) at 1500×*g* for 15 s × 2, and MeOH was added to the homogenate according to the tissue weight. After the solvent was homogenized again by the same condition, the appropriate amount of MeOH (<100 μL) solving 10 mg tissue weight was transferred to a 2-mL glass tube. After the solvent was messed up to 175 μL by MeOH, 25 μL of MeOH containing internal standards was added to the solvent, and vortexed for 10 s. After the solvent was incubated for 2 hours on ice, 100 μL of chloroform was added, and vortexed for 10 s. After the solvent was incubated for 1 hour on ice, 20 μL of Milli-Q water was added, and vortexed for 10 s. After 10 min incubation on ice, the solvent was centrifugated at 2000×*g* for 10 min at 4 °C, and the supernatant was transferred to the LC-MS vial (Agilent Technologies). The biological samples were analyzed by Method 1, 5, or 6.

*Cultured cells*

3T3-L1 cell line was seeded by $5\times10^4$ cells/well, cultured in 1 mL D-MEM (high glucose) with L-glutamine and phenol red (Wako, Japan) and incubated in 5% $CO_2$ at 37 °C in 12-well cell culture multiwell plates (Greiner bio-one). After 2 days where cells were reached to a confluency of ~70%, the medium was removed and cells were cultured in 1 mL MDI composing of 0.5 mM IBMX (Sigma–Aldrich), 1 μM dexamethasone (Sigma–Aldrich), and 10 μg/mL insulin (Sigma–Aldrich). After 2 days, the MDI induction medium was removed and replaced with 1 mL insulin medium. After 2 days, the insulin medium was removed and replaced with fresh D-MEM. After 4 days where the medium was changed every 2 days, the medium was aspirated, and cells were washed on the plates with PBS buffer. The solution was transferred to a new glass tube, and then dried with a vacuum dryer.

C2C12 cell line was seeded by $1\times10^5$ cells/well, cultured in 1 mL D-MEM (high glucose) with L-glutamine and phenol red (Wako, Japan) containing 10% fetal bovine serum (FBS) and Pen Strep glutamine (PSG), and incubated in 5% $CO_2$ at 37 °C in 24-well cell culture multiwell plates (Greiner bio-one). After over-night cultivation, the medium was aspirated, and cells were washed on the plates with PBS. After the PBS buffer was aspirated, 600 μL of MeOH was added and the cells were detached by cell scraper. The solution was transferred to a new glass tube, and then dried with a vacuum dryer.

The HeLa and HEK293 cells were cultivated as described previously[24]. HEK293- and HeLa cell lines were cultured in D-MEM (high-glucose) with L-glutamine (Wako) containing 10% fetal bovine serum (FBS) and Pen Strep glutamine (PSG), and incubated in 5% $CO_2$ at 37 °C in six-well cell culture multiwell plates (Greiner bio-one). After 1 hour cultivation, the medium was aspirated, and cells were washed on the plates with 10 mM Tris–HCl (pH 8.0) buffered saline.

The lipid extraction was performed by the same mixed solvent protocol ($MeOH:CHCl_3:H_2O$, 1:2:0.2, v/v/v) as described in human plasma (for method 1 and 2) section. First, chloroform was added to dried cells in a tube, followed by 30-s sonication. After 60-min incubation at room temperature, methanol was added and vortexed for 10 s. After 120-min incubation, Milli-Q water was added, vortexed for 10 s, and left the tube to stand for 10 min. The cells were then centrifuged at $2000\times g$ for 10 min at 20 °C. Supernatant was transferred to LC/MS vials to determine the phosphorous contents of the lipid fraction. The phosphorus content of the extracted lipids was quantified by the method of Bartlett[40], and then the extracted lipids were gently dried with $N_2$ and reconstituted with the mixed solvent of chloroform and methanol (1:2, v/v) to 0.5 mM, 0.8 mM, 1.0 mM, and 1.5 mM phosphorus for C2C12, HEK293, HeLa, and 3T3-L1, respectively, and transferred to the LC-MS vial. The biological samples were analyzed by method 2.

*Plant materials*

The heat-stressed/non-stressed Arabidopsis leaves were prepared as the benchmark for plant lipidomics using liquid chromatography ion mobility-tandem mass spectrometry (LC-IM-MS/MS). In addition, the lipid extracts from Arabidopsis (*Arabidopsis thaliana*), rice (*Oryza sativa*), and eggplant (*Solanum melongena*) were analyzed as the benchmark in using a conventional LC-MS/MS (DDA) method.

For method 3 and 4, plant growth condition was described previously[41]. *Arabidopsis thaliana* ecotype Columbia (Col-0) was used. Arabidopsis seeds were surface-sterilised and sown on an agar-solidified Murashige and Skoog medium containing 0.5% (w/v) sucrose. Plants were grown at 22°C under a 16-h-light/8-h-dark cycle (40 to 75 μmol photons $s^{-1}$ $m^{-2}$ of fluorescent lamp). Seventeen-day-old Col-0 plants at about 3 h after the onset of the light phase were subjected to heat stress at 38°C for one day (biological replicate $n$ = 3). Plants grown at 22°C were examined as the normal growth condition. Whole upper-ground parts from two different plants were harvested and pooled as a biological replicate. The plant material was immediately weighed in a 2.0-mL microcentrifuge tube (Sarstedt), frozen in liquid nitrogen and stored at −80°C until use.

The method for lipid extraction was described previously[42]. Under cryogenic conditions with liquid nitrogen, the plant material in the 2.0-mL microcentrifuge tube was milled by shaking at 900 rpm for 2 min on the Shake Master Neo (BMS, Tokyo, Japan) using a zirconia bead. From the frozen powdered plant material, total lipids were extracted by adding a 20-fold volume of extraction solvent (20 μL/mg F.W. plant sample) (chloroform/methanol/water, 50/100/31.45, v/v/v) with 1 μM 1,2-didecanoyl-*sn*-

glycero-3-phosphocholine, Sigma-Aldrich) as the internal standard (20 pmol $mg^{-1}$ F.W. plant sample). After stirring on a vortex mixer vigorously, the homogenate was incubated for 5 min at room temperature and centrifuged at 10,000×*g* for 10 min at room temperature. Supernatant (200 μL) was transferred to a new 1.5-mL microcentrifuge tube and mixed with chloroform (52.6 μL) and water (52.6 μL). After stirring on a vortex mixer vigorously, the mixture in the 1.5-mL microcentrifuge tube was incubated on ice for 10 min in darkness and centrifuged at 1,000×*g* for 10 min. The lower organic layer (85 μL) was transferred to a new 1.5-mL microcentrifuge tube and dried by the centrifugal concentrator (ThermoSavant SPD2010, Thermo Fisher Scientific) at room temperature. The dried lipid extract was dissolved in 160 μL of ethanol, centrifuged at 14,000×*g* for 10 min and transferred to a vial with a glass insert for LC-MS analysis.

For method 7, Arabidopsis (*Arabidopsis thaliana* ecotype Columbia (Col-0)), rice (*Oryza sativa* cv. Nipponbare), and eggplant (*Solanum melongena*) were used in this study. Arabidopsis and eggplant were grown on soil (PROMIX BX (Premier Horticulture Inc., Quakertown, PA): vermiculite = 2:1) supplemented with fertilizer in a greenhouse at 22°C under fluorescent light (16 h light/8 h dark cycle, 55 μmol photons $s^{-1}$ $m^{-2}$) for 7 weeks and 37 days, respectively. Rice was grown on wet commercial soil (Bonsol II (Sumitomo Chemical, Tokyo, Japan)) in a greenhouse with sunlight under a 16-h fluorescent light (28°C)/8-h dark (20°C) cycle[43]. Plants were kept under constant subirrigation conditions with tap water and fertilizer for 5 weeks. The entire aboveground (aerial) parts were harvested, immediately frozen in liquid nitrogen and lyophilized by the freeze dryer FDU-2200 (EYELA, Tokyo, Japan).

The method for lipid extraction was described previously[44]. The freeze-dried plant material was milled by a blender (IFM-800DG, Iwatani, Tokyo, Japan), filtered with a screen (0.5 mm) and transferred into a 2.0-mL microcentrifuge tube containing a zirconia bead. Lipids were extracted from the freeze-dried powdered plant material with a 160-fold volume of extraction solvent (160 μL $mg^{-1}$ D.W. plant sample) (methyl *tert*-butyl ether/methanol = 3/1, (v/v) with 1 μM 1,2-didecanoyl-*sn*-glycero-3-

phosphocholine, Sigma-Aldrich) as the internal standard (160 pmol mg$^{-1}$ D.W. plant sample). After vigorous stirring on a vortex mixer, a 50-fold volume of water (50 μL mg$^{-1}$ D.W. plant sample) was added to the homogenate. After vigorous stirring on a vortex mixer and dark incubation for 15 min on ice, the homogenate was centrifuged at 3,000×*g* for 10 min. The upper layer (160 μL) was transferred to a new 1.5-mL microcentrifuge tube. The organic phase was dried by the centrifugal concentrator (ThermoSavant SPD2010, Thermo Fisher Scientific) at room temperature. The dried lipid extract was dissolved in 200 μL of ethanol, centrifuged at 14,000×*g* for 10 min and transferred to a vial with a glass insert for LC-MS analysis.

*Published data resources*

At RIKEN DropMet website (http://prime.psc.riken.jp), the ID-DM0022[22] for nine algal lipidomics data using SCIEX TripleTOF 5600 DDA and the ID-DM0030[21] for 11 mouse tissues data using SCIEX TripleTOF 5600 DD were downloaded and reanalyzed by MS-DIAL 4.0.

### Reference (Supplementary Note 1)

**Supplementary Note 2 | Details of tandem mass spectrometry (MS/MS) characterization for 117 lipid subclasses**

We describe the details of mass fragmentation for 117 lipid subclasses. Note that the peak quantification of lipids to create our lipidomics database was performed according to the representative adduct type in each lipid subclass. For example, the ceramide containing sphingosine and nonhydrolyzed acyl chain, termed as Cer-NS, was quantified by $[M+CH_3COO]^-$ in negative ion mode although the molecule can also be characterized as $[M+H]^+$ and $[M-H]^-$ in MS-DIAL 4. The adduct type for lipid quantification was determined by the ion sensitivity and the practical annotation handiness. Moreover, the description for MS/MS spectrum annotation for the acetate adduct form could be replaced by the formate adduct form. The only difference of mass spectra between acetate and formate adduct forms is the *m/z* value of the precursor ion derived from the $CH_2$ (14.016 Da) difference, whereas the MS/MS pattern of product ions when using ammonium acetate as the LC solvent is almost equivalent as that obtained using ammonium formate according to our experience. The acyl chain moiety is characterized by the carbon and double bond numbers, such as 18:2 indicating 18 carbon and 2 double bonds in the acyl chain. The number of oxygen atoms involved with acyl chain incorporation; e.g., in oxidized phospholipids and ceramides, is separated by a semi-colon: e.g., Cer 18:1;O2/18:0;O, which is usually described in mammalian studies as Cer d18:1/18:0h, incorporating sphingosine and hydroxy fatty acid in the acyl chain moiety. When the hydroxy position can be determined from the MS/MS information, the description becomes Cer 18:1;(1OH, 3OH)/18:0;(2OH) which is known as Cer-AS in mammalian ceramides[18]. The ether- and vinyl ether linkages are described by O-18:0 and P-18:0, respectively. The *N*-acyl linkage is described by N-18:0 except for ceramide species. Underlining "_" is used to describe acyl chain compositions if the *sn1*, *sn2*, *sn3*, and *sn4* positional isomers are uncharacterized whereas the virgule (/) character is used if the acyl chain position is determined in a specific position such as *N*-acyl chain, ether chain, or acyl sugar.

The annotation is mostly performed according to the fragment ion of the alkyl ketone (termed as "Acyl ion"), dehydromonoacylglycerol (termed as "DMAG ion"), dehydrodiacylglycerol (termed as "DDAG ion"), or the neutral loss of alkyl ketone (termed as "Acyl loss") in positive ion mode and by the fragment ion of fatty acid (termed as "FA ion") in negative ion mode, respectively, in addition to lipid subclass specific diagnostic ions. The ontology, nomenclature for species level and molecular species level, and confirmation level are exemplified as {ontology, species level, molecular species level (if needed), confirmation level}.

**Annotations in LipidMAPS Fatty Acyls [FA]: Supplementary Fig. 2.** Acyl carnitine (CAR, CAR 18:1, level 1) was confirmed as the $[M]^+$ form by the unique product ion of *m/z* 85.028 ($C_4H_5O_2^+$) in positive ion mode. *N*-acyl ethanolamine (NAE, NAE 20:4, level 1) and fatty acid (FA, FA 18:1, level 1) were only confirmed by the consistency with the experimental or predicted retention times and precursor *m/z* values in positive and negative ion mode, respectively. The fatty acid ester of hydroxyl fatty acid (FAHFA, FAHFA 34:1, FAHFA 16:0/18:1, level 1) was confirmed as the $[M-H]^-$ form by the product ion and the neutral loss of esterified fatty acid in negative ion mode. *N*-acyl amides including *N*-acyl glycine[17,45] (NAGly, NAGly 31:0, NAGly 15:0/16:0, level 2), *N*-acyl glycyl serine[17,45] (NAGlySer, NAGlySer 31:0, NAGlySer 15:0/16:0, level 2), and *N*-acyl ornithine[17,45] (NAOrn, NAOrn 31:0, NAOrn 15:0/16:0, level 2) were mainly assigned and quantified as the $[M+H]^+$ form, and were double-checked in negative ion mode data where the metabolites were observed as $[M+CH_3COO]^-$ or $[M-H]^-$. The *m/z* 76.039, 106.05, 145.062, and 115.087 in positive ion mode constituted the characteristic ions for glycine, serine, glycyl serine, and ornithine moieties, respectively. The fatty acid ester form was characterized in this study, and the acyl chain moiety of esterified fatty acid was characterized by the acyl plus $H_2O$ neutral losses in both positive and negative ion mode data.

**Annotations in LipidMAPS Glycerolipids [GL]: Supplementary Fig. 3.** For monoacylglycerolipids, monoacylglycerol (MG, MG 18:0, level 1) was assigned as the $[M+NH_4]^+$ form, and the neutral loss (35 Da) of $NH_3$ and $H_2O$ from the precursor *m/z* was used for the lipid class confirmation. The lysodiacylglyceryl-3-O-carboxyhydroxymethylcholine[46] (LDGCC, LDGCC 16:0, level 2) was characterized as the $[M+H]^+$ form using the product ions of *m/z* 104.107 ($C_5H_{14}NO^+$) and 132.102 ($C_6H_{14}NO_2^+$) associated with the polar head moiety. The lysodiacylglyceryltrimethylhomoserine (LDGTS, LDGTS 18:1, level 2) and lysodiacylglyceryl hydroxymethyl-*N*,*N*,*N*-trimethyl-β-alanine (LDGTA, LDGTA 18:1, level 2) lipid subclasses[22] could not be distinguished at least in our low-energy collision induced dissociation (CID) condition, and the annotation was determined as LDGTS/A through the confirmation of *m/z* 144.102 ($C_7H_{14}NO_2^+$) and 236.149 ($C_{10}H_{22}NO_5^+$) values.

For diacylglycerolipids, the diacylglyceryl glucuronide (DGGA, DGGA 34:2, DGGA 16:0_18:2, level 1) was basically characterized and quantified as $[M-H]^-$ in negative ion mode because of higher sensitivity than that in positive ion mode whereas the DGGA lipid subclass could be characterized in positive ion mode by the unique neutral loss of 211.07 Da ($C_6H_{10}O_7$ and $NH_3$). According to our observation, the sensitivity of sulfoquinovosyl diacylglycerol (SQDG, SQDG 34:1, SQDG 16:0_18:1, level 1) is often higher in negative ion mode than that of positive ion mode. However, this depends on the injection volume and the matrix effect in biological samples. Moreover, the characterization for both lipid class and acyl chain moieties for SQDG is easier in positive ion mode than in negative ion mode because of the low sensitivities of fragment ions in negative ion mode. Therefore, the representative adduct type for SQDG lipid quantification was determined in each study independently. The neutral loss of $C_6H_{10}O_7S$ (sulfoquinovosyl moiety) plus $NH_3$ (adduct ion moiety) and the product ion of *m/z* 225.007 ($C_6H_9O_7S^-$) were used to characterize the SQDG lipid subclass in positive and negative ion mode, respectively. Both monogalactosyldiacylglycerol (MGDG, MGDG 34:1, MGDG 16:0_18:1, level 1) and digalactosyldiacylglycerol (DGDG, DGDG 34:1, DGDG 16:0_18:1, level 1) lipid subclasses were characterized and quantified as $[M+CH_3COO]^-$ and the product ions of *m/z* 253.093 (monogalactosyl moiety, $C_9H_{17}O_8^-$) and *m/z* 397.135 (digalactosyl moiety, $C_{15}H_{25}O_{12}^-$) were used to characterize the MGDG and DGDG lipid subclasses, respectively. As the sensitivity of the MGDG-specific product ion, i.e. *m/z* 253.093, is often very low in negative ion mode, the classification of MGDG lipid in negative ion mode was performed only by checking the FA ions from two acyl chain moieties whereas the lipid class could be double-checked in positive ion mode by the neutral loss of $C_6H_{12}O_6$ (monogalactocyl moiety) plus $NH_3$ (adduct moiety). The DGTS[22] (DGTS 34:1, DGTS 16:0_18:1, level 2) and DGCC[46] (DGCC 34:1, DGCC 16:0_18:1, level 2) lipid classes could be characterized by the same characteristic ions as described in the lyso-type of LDGTS and LDGCC. The acyl chains were elucidated according to the acyl losses.

For alkylacylglycerolipids, our platform distinguished the ether type molecule from the ester type one of the same polar head lipid class by the accurate mass and the unique fragmentation associated to ether chains. The alkylacyl form of MGDG[47] (EtherMGDG, MGDG O-34:1, MGDG O-16:0_18:1, level 2) could be characterized and quantified as $[M+CH_3COO]^-$ in negative ion mode by the confirmation of the FA ion and the acyl loss derived from the ester-linked acyl chain moiety. The EtherMGDG lipid could also be annotated as $[M+NH_4]^+$ by the confirmation of the DMAG ion and neutral loss of $C_6H_{12}O_6$ plus $NH_3$ as described in the MGDG subclass. Notably, the same polar head lipid class could also be distinguished by the 30–40 mDa mass difference of precursor *m/z* values: for example, the *m/z* value of MGDG O-16:0_16:0 (*m/z* 775.59409) in $[M+CH_3COO]^-$ adduct type exhibits 36 mDa difference from that of MGDG 15:0_16:0 (*m/z* 775.5577) in the same adduct type, demonstrating that high resolution MS offers a distinct advantage in lipid profiling. The semino lipid[47] (EtherSMGDG, SMGDG O-30:0,

SMGDG O-16:0_14:0, level 2) was characterized as $[M-H]^-$ according to the product ions of *m/z* 241.002 ($C_6H_9O_8S^-$) and *m/z* 96.96 ($HSO_4^-$). The acyl chain moiety was determined by the neutral loss of acyl 14:0 and $H_2O$. The alkylacyl form of DGDG (EtherDGDG, DGDG O-34:2, DGDG O-18:2_16:0, level 3) could only be characterized as $[M+CH_3COO]^-$ in this study, and the annotation could be performed by the presence of the FA ion and the acyl loss from the ester-linked acyl moiety. Note that the product ion of the digalactosyl moiety observed as *m/z* 397.135 in DGDG could not be detected in the alkylacyl form. The alkylacyl type of DG[48] (EtherDG, DG O-34:1, DG O-16:0_18:1, level 2) was characterized as $[M+NH_4]^+$ by confirming the DMAG ion derived from the ester-linked moiety.

For triacylglycerols, the triacylglycerol lipid class (TG, TG 56:8, TG 16:0_18:2_22:6, level 1) was characterized as $[M+NH_4]^+$ and the acyl chains could be annotated by the neutral losses of acyl and $H_2O$. The sodium adduct type of TG ($[M+Na]^+$) was also characterized in our platform to reduce the chances of false positive annotations by the other lipid classes. The alkyldiacyl type of TG[48] (EtherTG, TG O-56:7, TG O-16:0_18:1_22:6, level 2) was characterized as $[M+NH_4]^+$: the EtherTG lipids eluted later than normal TG lipids. The acyl chain moieties could be characterized by the neutral losses of acyl and $H_2O$ where the neutral loss of alkyl chain could also be observed. The acylated DGGA (ADGGA, ADGGA 50:2, ADGGA 16:0_16:1_18:2, level 3) was characterized and quantified as $[M+NH_4]^+$ because the unique product ion of the acylated glucuronosyl moiety could be detected in positive ion mode whereas the other acyl chains were characterized by DMAG ions. Therefore, the acyl chain linked to the glucuronosyl moiety was separated by chain length with the brackets from the other acyl chain descriptions; for example, ADGGA (O-16:0)16:1_18:2 indicated that FA 16:0 is incorporated into the glucuronosyl moiety and the others are linked to the glycerol moiety. Alternatively, all chain compositions were connected by an underline, such as ADGGA 16:0_16:1_18:1 because the acyl chains could only be characterized by FA ions in negative ion mode.

**Annotations in LipidMAPS Glycerophospholipids [GP]: Supplementary Fig. 4.** Among monoacylglycerophospholipids, lysophosphatidylcholine (LPC, LPC 18:1, level 1) and its alkyl type[48] (EtherLPC, LPC O-18:1, level 2) were quantified as $[M+H]^+$ and could also be characterized as $[M+CH_3COO]^-$. The product ion of *m/z* 184.074 ($C_5H_{15}NO_4P^+$) and the neutral loss of 74.037 ($CH_3COOH+CH_2$) were used as the diagnostic fragment ion for LPC and Ether LPC for positive and negative ion mode, respectively. According to a previous report[49], the acyl chain position of LPC is determined as *sn1* by default if the ion abundance of *m/z* 104.107 ($C_5H_{14}NO^+$) exceeds 10% of the base peak. In addition, the ion from the neutral loss of $C_3H_9N$ (59.073) was used as the characteristic ion for the sodium adduct type of LPC. The lysophosphatidylethanolamine (LPE, LPE 18:1, level 1) and its alkyl type[48] (EtherLPE, LPE O-18:1, level 1) were quantified as $[M+H]^+$ and could also be characterized as $[M-H]^-$. The neutral loss of $C_2H_8NO_4P$ (141.02 Da) and the product ion of *m/z* 196.036 ($C_5H_{11}NO_5P^-$) were utilized as the diagnostic criteria for these lipid classes in positive and negative ion mode, respectively. Because of the low sensitivity of *m/z* 196.036, confirmation of the FA ion must be performed in negative ion mode. In addition, the EtherLPE was annotated as the plasmenyl type (plasmalogen) if the "Ether" fragment ion was observed (annotated as LPE P-18:0): the fragment is defined by cleavage of the ether link, resulting in the product ion of $R\text{-}C{=}C\text{–}O^-$ where R denotes the acyl moiety. The lysophosphatidic acid (LPA, LPA 16:0, level 1) was quantified as $[M-H]^-$, and *m/z* 152.996 ($C_3H_6O_5P^-$) was utilized as the diagnostic ion. Note that the FA ion was basically not detected from LPA lipids. The lysophosphatidylglycerol (LPG, LPG 18:1, level 1) and its alkyl type[48] (EtherLPG, LPG O-18:1, level 2) were characterized and quantified as $[M-H]^-$, where *m/z* 152.996 ($C_3H_6O_5P^-$) was utilized for the diagnostic ion. Because of the low sensitivity of *m/z* 152.996, the FA ion was practically utilized for LPG characterization. When the Ether ion was detected, the Ether LPG was annotated as the plasmenyl type as described in EtherLPE. Both lysophosphatidylserine (LPS, LPS 18:0, level 1) and lysophosphatidylinositol (LPI, LPI 18:1, level 1) were quantified as $[M-H]^-$. The product ions of *m/z* 241.008 ($C_6H_{10}O_8P^-$) and *m/z* 315.049 ($C_9H_{16}O_{10}P^-$) were used for LPI and the neutral loss of $C_3H_5NO_2$ (87.032 Da) were used for LPS annotations, respectively: the FA ion especially for LPI was used for the diagnostic criterion because of the low sensitivity of phosphoinositol moiety-derived ions.

For diacylglycerophospholipids, all lipid classes except for bismonoacylglycerophosphate (BMP, BMP 40:5, BMP 18:1_22:4, level 1) and plasmenyl PE (EtherPE_P, PE P-38:3, PE P-18:0_20:3, level 1) were quantified in negative ion mode because both acyl chains and lipid class specific ions could easily be detected, decreasing the number of false positive annotations. Phosphatidic acid (PA, PA 34:1, PA 16:0_18:1, level 1) was only characterized as $[M-H]^-$ in negative ion mode by the confirmation of FA ions from both acyl chains. The phosphatidylcholine (PC, PC 38:4, PC 18:0_20:4, level 1) lipid exhibited the unique product ion of *m/z* 184.074 ($C_5H_{15}NO_4P^+$) and the neutral loss of 74.037 ($CH_3COOH+CH_2$) for $[M+H]^+$ and $[M+CH_3COO]^-$ forms, respectively, as described in the lyso-type form. The acyl chains could easily be determined based on FA ions in negative ion mode whereas they could also be characterized by acyl losses in positive ion mode although the sensitivity was quite low. The sodium adduct form of PC was characterized by two neutral losses of $C_3H_9N$ (59.073 Da) and $C_5H_{14}NO_4P$ (183.066 Da). The plasmenyl or plasmanyl type of PC (EtherPC, PC O-38:4, PC O-18:0_20:4, level 1) could also be confirmed by the same lipid class specific ions of PC whereas the acyl chain was only determined in negative ion mode according to the confirmation of the FA ion derived from the ester-linked moiety. Note that the plasmanyl and plasmenyl types were not distinguished in our annotation pipeline for EtherPC because of the low sensitivity of plasmenyl-specific product ions. Phosphatidylethanolamine (PE, PE 38:4, PE 18:0_20:4, level 1) was characterized as $[M+H]^+$ and $[M-H]^-$ by the neutral loss of $C_2H_8NO_4P$ (141.02 Da) and the product ion of *m/z* 196.038 ($C_5H_{11}NO_5P^-$) in positive- and negative ion mode, respectively.

The product ions of alkyl ketones (termed as “Acyl ion”) and FA ions were utilized to characterize the acyl chain moieties in positive and negative ion mode, respectively, although the sensitivity of the acyl ion was very low. The sodium adduct type of PE was annotated according to the neutral losses of $C_2H_5N$ (43.04 Da) and $C_2H_8NO_4$ (110.045 Da). The plasmanyl (EtherPE_O, PE O-34:1, PE O-16:0_18:1, level 1) and plasmenyl (EtherPE_P, PE P-38:3, PE P-18:0_20:3, level 1) types of PE could be characterized in positive ion mode whereas the difference between vinyl ether and normal ether types was not observed in ESI (−)-MS/MS. The plasmalogen type of PE was characterized by the existence of the DMAG ion derived from the neutral loss of the vinyl ether acyl moiety whereas the fragment ion could not be observed in the plasmanyl type of PE in which only the neutral loss of the PE polar head ($C_2H_8NO_4P$, 141.02 Da) and the acyl ion were detected. Alternatively, the plasmenyl/plasmanyl type of PE (EtherPE) could easily be characterized in negative ion mode by the FA ion and acyl loss derived from the ester chain moiety. Phosphatidylglycerol (PG, PG 38:4, PG 18:0_20:4, level 1) was characterized as $[M+NH_4]^+$ and $[M-H]^-$ by the neutral loss of 171.006 Da ($C_3H_9O_6P+NH_3$) and the product ion of *m/z* 152.996 ($C_3H_6O_5P^-$), respectively. Because of the low ion sensitivity of *m/z* 152.996, the PG lipid was annotated as molecular species level if both FA ions of the two acyl chains were observed in ESI (−)-MS/MS. In comparison, the DMAG ions were used as the diagnostic ions in in ESI (+)-MS/MS: the lipid was annotated as molecular species level when both DMAG ions were detected. The alkylacyl type of PG[48] (EtherPG, PG O-34:1, PG O-18:1_16:0, level 2) was only characterized in negative ion mode by the confirmation of FA ion derived from the ester-linked chain moiety. According to our observation, the ether ion where an oxygen atom was rearranged to the ether chain moiety could be observed in the EtherPG lipid and might occur in the plasmalogen type of EtherPG. As we could not find an EtherPG lipid having no double bond in the alkyl chain moiety in this study, the issue will be resolved in future work. The BMP lipid (BMP 40:5, BMP 18:1_22:4, level 1) was only characterized in positive ion mode by the confirmation of DMAG ions from both acyl chains because the MS/MS spectrum could not be distinguished from PG lipid in negative ion mode. Phosphatidylinositol (PI, PI 38:4, PI 18:0_20:4, level 1) could be characterized as $[M+NH_4]^+$, $[M+Na]^+$, and $[M-H]^-$ where the acyl chains could be resolved in negative ion mode. The neutral loss of 277.056 Da ($C_6H_{14}O_9P+NH_3$) and the product ion of *m/z* 283.019 ($C_6H_{13}O_9PNa^+$) were used as the PI-specific ion in $[M+NH_4]^+$ and $[M+Na]^+$, respectively. The product ions of *m/z* 241.008 ($C_6H_{10}O_8P^-$) and *m/z* 315.049 ($C_9H_{16}O_{10}P^-$) served as the PI class specific ions in $[M-H]^-$ and the annotation was exported as acyl-chain resolved when FA ions from both acyl chains were detected. The alkyl type of PI[48] (EtherPI, PI O-36:4, PI O-16:0_20:4, level 2) was annotated by the FA ion from the ester chain moiety with the confirmation of PI-specific ions. Phosphatidylserine (PS, PS 38:4, PS 18:0_20:4, level 1) was characterized as $[M+H]^+$, $[M+Na]^+$, and $[M-H]^-$ by the neutral losses of $C_3H_8NO_6P$ (185.009 Da) and $C_3H_5NO_2$ (87.032 Da) in positive- and negative ion mode, respectively. The acyl losses and FA ions were utilized for the annotation of acyl chains in $[M+H]^+$ and $[M-H]^-$, respectively. The alkyl type of PS[48] (EtherPS, PS O-38:6, PS O-16:0_22:6, level 2) was characterized as $[M-H]^-$ by the confirmation of acyl loss from the ester chain moiety with the PS-specific neutral loss of 87.032 Da. Dilysocardiolipin[50] (DLCL, DLCL 34:2, DLCL 16:0_18:2, level 2) was characterized as $[M-H]^-$, and the fragment ion matched with the LPA moiety was utilized rather than the FA ion for acyl chain annotations. The product ion of *m/z* 152.996 ($C_3H_6O_5P^-$) was used as the diagnostic ion for the DLCL lipid class. Phosphatidylmethanol (PMeOH, PMeOH 34:2, PMeOH 16:0_18:2, level 1) and phosphatidylethanol (PEtOH, PEtOH 38:4, PEtOH 18:0_20:4, level 1) were characterized by the existence of FA ions from both acyl chain moieties because of the low sensitivity of the product ions of *m/z* 110.985 ($CH_4O_4P^-$) and *m/z* 125.001 ($C_2H_6O_4P^-$), which could be utilized as PMeOH and PEtOH-specific fragment ions, respectively.

The oxidized phospholipids were characterized and quantified in negative ion mode. The oxidized PC, PE, PG, PI, and PS (OxPC, OxPE, OxPG, OxPI, and OxPS, level 2[46]) were first confirmed by their lipid class-specific ions of 74.037 Da neutral loss, *m/z* 196.038 product ion, *m/z* 152.996 product ion, *m/z* 241.008/315.049 product ions, and 87.032 Da neutral loss, respectively. With regard to nomenclature, the oxidized form was described using a semi-colon (;) and the additional oxygen count as in PE 18:0_20:4;4O. Because the current program does not target the oxidized lipid having two oxidized acyl chains, the non-oxidized fatty acid ion was secondly utilized as the diagnostic ion. Because the oxidized fatty acid ion could be detected as the native fatty acid form, one $H_2O$-loss form, or two $H_2O$-loss form, depending on the oxidation number and structural conformation, the annotation was successful if at least one of these three forms was confirmed in the MS/MS spectrum. The alkyl type of OxPE[46] (EtherOxPE, PE O-38:6;O, PE O-16:0_22:6;O, level 2) was annotated by the product ion and neutral loss of the ester-linked oxidized fatty acid moiety.

The lyso-type of *N*-acyl PE[48] (LNAPE, LNAPE 34:2, LNAPE 16:0/N-18:2, level 2) and *N*-acyl PS[51] (LNAPS, LNAPS 34:0, LNAPS 18:0/N-16:0, level 2) were quantified and characterized in negative ion mode. In contrast to PE, the product ion of *m/z* 196.038 was not observed in LNAPE form; instead, the product ion of *m/z* 152.996 was detected. Moreover, the fragment ion of *N*-acyl chain was not detected at least in our experimental condition whereas the FA ion from the ester-link moiety was used for the acyl chain characterization. In comparison to PS, identically, the neutral loss of 74.037 Da was not observed in LNAPS and the product ion of *m/z* 152.996 was detected instead. The product ion matched with the LPA moiety containing an ester-linked fatty acid could be used to characterize the acyl chain moiety.

For triacylglycerophospholipids, hemibismonoacylglycerophosphate (HBMP, HBMP 52:3, HBMP 18:1/16:1_18:1, level 1) was quantified as ether $[M+NH_4]^+$ or $[M-H]^-$, depending on the biological sample and the injection volume. The acyl chain of the LPG moiety was characterized in positive ion mode by the dehydrodiacylglycerol ion (termed as "DDAG" ion) containing two acyl chains of the PG substructure. The DMAG ions derived from the PG moiety were used to determine the other acyl chains. Alternatively, the FA ions from the PG moiety were detected in negative ion mode, and the product ion of DMAG plus the phosphate moiety was used to characterize the acyl chain involved in the LPG moiety. Lysocardiolipin[52] (MLCL, MLCL 56:7, MLCL 16:0_16:0_16:0, level 2) was quantified and characterized in negative ion mode, and the product ion of *m/z* 152.996 was utilized as the diagnostic ion of glycerophospholipid. Three FA ions derived from ester-linked moieties were used for acyl chain characterizations.

For cardiolipin (CL, level 1) incorporating four ester-linked acyl chains, the quantification was performed as $[M-H]^-$ because four acyl chains could be characterized by FA ions with the characteristic ion of *m/z* 152.996 in negative ion mode whereas the acyl chains were semi-resolved in positive ion mode: the DDAG ions incorporating two acyl chains were only available from the $[M+NH_4]^+$ form, resulting in the annotation as e.g. CL 34:0_36:2 for CL 16:0_18:0_18:1_18:1 at least in our experimental condition.

**Annotations in LipidMAPS Sphingolipids [SP]: Supplementary Fig. 5.** The sphingobase (SPB) moiety was described as (carbon number):(double bond);(hydroxy moiety count); for example, SPB 18:1;2O for sphingosine d18:1, indicating that the chain had one nitrogen, two hydroxy moieties, 18 carbons, and 1 double bond. When the hydroxy positions could be determined by the MS/MS spectrum, the sphingobase description became SPB 18:1;(1OH,3OH) for sphingosine d18:1. Moreover, the term of "SPB 18:1;2O ion" was used to describe the characteristic ion for the sphingobase moiety in both positive and negative ion mode. The term "SPB 18:1;2O" ion indicates *m/z* 300.2897 ($C_{18}H_{38}NO_2^+$) and *m/z* 298.2752 ($C_{18}H_{36}NO_2^-$) derived by the hydrogen rearrangement (HR) rule P2 and rule N1 in positive and negative ion mode, respectively. The SPB ion could often be detected in the form of one or two $H_2O$ losses, with the product ion being described as “SPB 18:0;2O-$H_2O$” or “SPB 18:0;2O-2$H_2O$”, respectively. For the intact sphingobase molecules, the sphinganine (DHSph, SPB 17:0;2O, level 1) and sphingosine (Sph, SPB 18:1;2O, level 1) were characterized as $[M+H]^+$ by the neutral losses of $H_2O$, double $H_2O$, and $CH_4O_2$ moieties, and the phytosphingosine (PhytoSph, SPB 18:0;3O, level 1) was also characterized as $[M+H]^+$ by the neutral losses of double and triple $H_2O$ and $CH_6O_3$ moieties.

For sphingolipids incorporating the phosphocholine polar head, the normal sphingomyelin (SM, SM 34:1;2O, SM 18:1;2O/16:0, level 1) was quantified as $[M+CH_3COO]^-$. The SM lipid class could be characterized by the specific product ion of *m/z* 168.043 ($C_4H_{11}NO_4P^+$) and neutral loss of 74.037 Da ($CH_3COOH+CH_2$) in negative ion mode, and the product ion of *m/z* 184.074 ($C_5H_{15}NO_4P^+$) was used in positive ion mode. The acyl chain was annotated by the neutral loss of acyl and methyl moieties and the product ion of “SPB-$H_2O$” in negative and positive ion mode, respectively. Because of the low sensitivity of acyl chain-related ions, the sphingomyelin was often annotated by the summed form, such as SM 34:1;2O for SM 18:1;2O/16:0. The sodium adduct form of SM was characterized by the neutral losses of $C_3H_9N$ (59.073 Da) and $C_5H_{14}NO_4P$ (183.066 Da). The SM lipid incorporating the phytosphingosine or a hydroxy fatty acid[53] (SM+O, SM 34:1;3O, level 2) was quantified as $[M+CH_3COO]^-$, and it could be characterized by the same characteristic ions of normal SM lipid class as $[M+H]^+$ and $[M+CH_3COO]^-$. Notably, the acyl chain specific ion could not be observed in our experimental data set, and therefore the SM+O lipid class was only reported as the summed description. The acylated SM (ASM, SM 52:2;3O, SM 34:1;2O(FA 18:0), level 3) was quantified as $[M+CH_3COO]^-$, and characterized in both positive and negative ion modes. The acylated position was defined at the hydroxy moiety in the sphingobase because it was the only plausible position for the ester reaction. In addition to the SM characteristic ions, the FA ion and the neutral loss derived from the acylated moiety could be detected in negative ion mode, and the neutral loss of the acylated moiety was also used for the characterization in positive ion mode. Moreover, the diagnostic ion to characterize the sphingobase and *N*-acyl chain moieties could not be detected in our experimental data set; therefore, the ceramide moiety was reported as the summed description.

The description and characterization of ceramide lipid classes have been reported previously for negative ion mode data[18]. The ceramide lipids were basically quantified as $[M+CH_3COO]^-$ except for di- and tri-hexosyl ceramides because the hydroxy position of *N*-hydroxy fatty acid could be determined in ESI(−)-MS/MS. In addition, the sensitivity was higher in acetate adduct form than that of proton loss form. Moreover, the example lipid structure was used to explain the diagnostic ions for ceramide characterizations because the mass fragmentation was more complex than that of glycerolipids. The ceramide lipid (Cer-NDS[24], Cer 34:0;2O, Cer 18:0;2O/16:0, level 2) incorporating sphinganine (DS) and fatty acid moieties (N) could be characterized as $[M+H]^+$, $[M-H]^-$, and $[M+CH_3COO]^-$. For the acetate adduct form of Cer 18:0;2O/16:0 in ESI(−)-MS/MS, the product ions of *m/z* 280.264 (Acyl 16:0+$C_2H_3N$), *m/z* 239.238 (SPB 18:0;2O-$C_2H_7NO$), and *m/z* 237.246 (Acyl 16:0-2H) were used to characterize the acyl chain moieties whereas the lipid class was determined by confirming the product ion of $[M-H]^-$ (*m/z* 538.53) and the neutral losses of $CH_4O$ (32.04 Da) and $CH_4O_2$ (48.03 Da) from the $[M-H]^-$ peak. Alternatively, the product ions of *m/z* 284.285 (SPB 18:0;2O-$H_2O$), *m/z* 266.284 (SPB 18:0;2O-2$H_2O$), and *m/z* 254.284 (SPB 18:0;2O-$CH_4O_2$) were used to characterize the acyl chains in positive ion mode. The ceramide lipid (Cer-NS, Cer 34:1;2O, Cer 18:1;2O/16:0, level 1) incorporating sphingosine (S) and fatty acid (N) moieties was also characterized as $[M+H]^+$, $[M-H]^-$, and $[M+CH_3COO]^-$. The scheme of mass fragmentation was the same as described in Cer-NDS where the *m/z* values were changed by the difference of double bond number in the sphingobase. The ceramide lipid (Cer-AP, Cer 38:0;4O, Cer 18:0;3O/20:0;(2OH), level 1) incorporating phytosphingosine (P) and alpha-hydroxy fatty acid (A) was characterized as $[M+H]^+$, $[M-H]^-$, and $[M+CH_3COO]^-$. For the acetate adduct form of Cer 18:0;3O/20:0;(2OH) in ESI(−)-MS/MS, the unique ion characterizing the hydroxyl position was detected as *m/z* 281.285 (Acyl 20:0;O-$CH_2O$, $C_{19}H_{37}O^-$) and the product ions of *m/z* 382.332 (Acyl 20:0;O+$C_3H_5NO$, $C_{23}H_{44}NO_3^-$) and *m/z* 328.291 (Acyl 20:0;O+O, $C_{20}H_{39}O_3^-$) were used for acyl chain characterizations. The product ions of *m/z* 316.285 (SPB 18:0;3O, $C_{18}H_{38}NO_3^+$), *m/z* 298.274 (SPB 18:0;3O-$H_2O$, $C_{18}H_{36}NO_2^+$), *m/z* 280.264 (SPB 18:0;3O-2$H_2O$, $C_{18}H_{34}NO^+$), and *m/z* 262.253 (SPB 18:0;3O-3$H_2O$, $C_{18}H_{32}N^+$) were the characteristic ions to determine acyl chains in positive ion mode.

The ceramide (Cer-ADS[24], Cer 34:0;3O, Cer 18:0;2O/16:0;(2OH), level 2) incorporating sphinganine (DS) and alpha-hydroxy fatty acid (A) moieties was characterized as $[M-H]^-$ and $[M+CH_3COO]^-$. For the acetate adduct form of Cer 18:0;2O/16:0;(2OH), the product ion of *m/z* 225.222 (Acyl 16:0;O-$CH_2O$) was used to characterize the hydroxy position and the product ions of *m/z* 282.28 (SPB 18:0;2O-$H_2O$, $C_{18}H_{36}NO^-$) and *m/z* 253.217 (Acyl 16:0;O-2H, $C_{16}H_{29}O_2^-$) were used to characterize the acyl chain moieties. Furthermore, the product ion of *m/z* 239.238 (SPB 18:0;2O-$C_2H_7NO$, $C_{16}H_{31}O^-$) was the unique ion characterizing the hydroxy position at γ-C in the sphinganine moiety. The ceramide (Cer-AS, Cer 34:1;3O, Cer 18:1;2O/16:0;(2OH), level 1) incorporating sphingosine (S) and alpha-hydroxy fatty acid (A) was characterized by the product ions derived from the same fragmentation scheme as those of Cer-ADS. The ceramide (Cer-BDS[24], Cer 34:0;3O, Cer 18:0;2O/16:0;(3OH), level 2) incorporating sphinganine (DS) and beta-hydroxy fatty acid (B) was characterized as $[M-H]^-$ and $[M+CH_3COO]^-$. For the acetate adduct form of Cer 18:0;2O/16:0;(3OH), the product ions of *m/z* 342.301 (SPB 18:0;2O+$C_2H_2O$, $C_{20}H_{40}NO_3^-$) and *m/z* 310.275 (*m/z* 342.301-$CH_4O$, $C_{19}H_{36}NO_2^-$) were used to characterize the hydroxy position and the sphingosine moiety. Furthermore, the product ion of *m/z* 239.238 (SPB 18:0;2O-$C_2H_7NO$, $C_{16}H_{31}O^-$) was the unique ion characterizing the γ-hydroxy position in the sphingosine moiety. The ceramide (Cer-BS[24], Cer 40:1;3O, Cer 16:1;2O/24:0;(3OH), level 2) incorporating sphingosine (S) and beta-hydroxy fatty acid (B) was characterized by the product ions derived from the same fragmentation scheme as those of Cer-BDS. In comparison, the hydroxy position of *N*-acyl chain could not be characterized in ESI(+)-MS/MS, and therefore the character of "H" instead of "A" or "B" was utilized to describe the existence of the hydroxy moiety in the *N*-acyl chain in the MS-DIAL ontology. For the $[M+H]^+$ form of Cer 20:1;2O/24:0;O incorporating SPB 20:1;2O (S) and hydroxy fatty acid 24:0;O (H), the product ions of *m/z* 310.311 (SPB 20:1;2O-$H_2O$, $C_{20}H_{40}NO^+$), *m/z* 292.3 (SPB 20:1;2O-2$H_2O$, $C_{20}H_{38}N^+$), and *m/z* 280.3 (SPB 20:1;2O-$CH_4O_2$, $C_{19}H_{38}N^+$) were used to characterize the acyl chain moieties. Moreover, the ceramide (Cer-HDS[24], Cer 38:0;3O, Cer 18:0;2O/20:0;O, level 2) incorporating sphinganine (DS) and hydroxy fatty acid (H) was characterized by the product ions derived from the same fragmentation scheme as those of Cer-HDS. In negative ion mode, the term of "A" and "B" was replaced by "H" when the product ion characterizing the hydroxy position was not observed in ESI(−)-MS/MS. According to our experience and a previous study[54], the case was considered as the ceramide incorporating omega-hydroxy fatty acid (O), which is known as the precursor of acyl ceramide at least in mammalian tissues.

The ceramide (Cer-EOS[24], Cer 70:3;4O, Cer 18:1;2O/32:0;O(FA 18:2), level 2) incorporating sphingosine (S), omega-hydroxy fatty acid (O), and esterified fatty acid (E) linked to the omega hydroxy moiety was characterized as $[M+H]^+$, $[M-H]^-$, and $[M+CH_3COO]^-$. For the acetate adduct form of Cer 18:1;2O/32:0;O(FA 18:2), the product ion of *m/z* 279.233 (FA 18:2) denoted the esterified fatty acid, and the neutral loss of Acyl 18:2 was also used to characterize the esterified acyl chain. The product ion of *m/z* 494.494 (*N*-acyl 32:0;O, $C_{32}H_{64}NO_2^-$) was used to characterize the *N*-acyl chain moiety. Because of the low sensitivity of the product ion involved with the *N*-acyl chain, the annotation was performed by the summed form as Cer d50:1(FA 18:2) if the ion was not detected in ESI(−)-MS/MS: the annotation term of "chain semi-resolved" was used in such case. Alternatively, the product ions of *m/z* 282.279 (SPB 18:1;2O-$H_2O$, $C_{18}H_{36}NO^+$), *m/z* 264.269 (SPB 18:1;2O-2$H_2O$, $C_{18}H_{34}N^+$), and *m/z* 252.269 (SPB 18:1;2O-$CH_4O_2$, $C_{17}H_{34}N^+$) were used to characterize the sphingosine moiety in ESI(+)-MS/MS whereas no fragment ion characterizing the other acyl chains was detected, at least in our experimental condition. Therefore, the annotation was described as the semi-resolved form as Cer 18:1;2O/52:3;2O: Note that the double bond for the ketone moiety was also counted, resulting in the term of 52:3;2O for summing 32:0;O(FA 18:2). The ceramide (Cer-EODS[24], Cer 49:1;4O, Cer 18:0;2O/16:0;O(FA 15:0), level 2) incorporating sphinganine (DS), omega-hydroxy fatty acid (O), and esterified fatty acid (E) was characterized by the same procedure as described in Cer-EOS. The ceramide (Cer-EBDS, Cer 50:1;4O, Cer 18:0;2O/17:0;(3OH)(FA 15:0), level 3) incorporating sphinganine (DS), beta-hydroxy fatty acid (B), and esterified fatty acid linked to the beta-hydroxy position was only characterized as $[M+CH_3COO]^-$ in feces samples. For the acetate adduct form of Cer 18:0;2O/17:0;(3OH)(FA 15:0), the product ion of *m/z* 241.217 (FA 15:0) and the neutral loss of Acyl 15:0;O were used to characterize the esterified fatty acid. Moreover, the product ion of *m/z* 342.3 (SPB 18:0;2O+$C_2H_2O$, $C_{20}H_{40}NO_3^-$), which is derived from the cleavage of the beta-hydroxy position of *N*-acyl chain as described in Cer-BDS, was used to characterize the sphingobase moiety and the existence of beta-hydroxy fatty acid.

The ceramide (Cer-NP, Cer 34:0;3O, Cer 18:0;3O/16:0, level 1) incorporating phytosphingosine (P) and normal fatty acid (N) was characterized as $[M-H]^-$ and $[M+CH_3COO]^-$. For the acetate adduct form of Cer 18:0;3O/16:0, the product ions of *m/z* 298.275 (SPB 18:0;3O-H2O, $C_{18}H_{36}NO_2^-$), *m/z* 267.232 (SPB 18:0;3O-$CH_7NO$, $C_{17}H_{31}O_2^-$), and *m/z* 255.232 (SPB 18:0;3O-$C_2H_7NO$, $C_{16}H_{31}O_2^-$) were used to characterize the acyl chain moieties. Furthermore, the product ion of *m/z* 310.274 (Acyl 16:0;O+$C_3H_5NO$, $C_{19}H_{36}NO_2^-$) was used as the unique ion characterizing the γ-hydroxy position in the sphingosine moiety.

The monohexosyl form[24] (HexCer) of Cer-NS, Cer-NDS, Cer-AP, Cer-HS, Cer-HDS, and Cer-EOS was characterized by the same product ions as described in the non-hexosyl forms with the confirmation of the hexosyl moiety loss (termed as “Hex” loss, $C_6H_{10}O_5$). Notably, the hydroxy position in the hydroxy fatty acid moiety was not determined because of the low sensitivity of product ions involved with the hydroxy position in the *N*-acyl chain. In ESI(+)-MS/MS, the product ions derived from the neutral losses of Hex-$H_2O$ and Hex-$2H_2O$ were also observed. The dihexosyl form (Hex2Cer, level 1) of Cer-NS was quantified as $[M+H]^+$ because the acyl chain moieties could only be characterized in positive ion mode. In addition to two sequential hexosyl losses, the product ions involved with SPB-$H_2O$, SPB-$2H_2O$, and SPB-$CH_4O_2$ were used to characterize the sphingobase moiety. The Hex2Cer lipid was also characterized as $[M+CH_3COO]^-$, where the sequential hexosyl losses and the product ion of *m/z* 179.056 (Hex ion, $C_6H_{12}O_6^-$) were used to characterize the lipid class: the Hex2Cer lipid in positive ion mode was always annotated as the summed form. For the trihexosyl form[49] (Hex3Cer, level 2), the annotation strategy was also the same as described for Hex2Cer.

The ceramide phosphoinositol[55] (PI-Cer, PI-Cer 34:0;3O, PI-Cer 18:0;2O/16:0;O, level 2) incorporating sphingosine and fatty acid moieties was quantified in negative ion mode whereas it could be characterized in both ion modes as $[M+H]^+$ and $[M-H]^-$. In this study, the PI-Cer lipid incorporating sphingosine and hydroxy fatty acid moiety was characterized. For PI-Cer 18:0;2O/16:0;O, the product ions of *m/z* 78.959 ($PO_3^-$) and *m/z* 241.012 ($C_6H_{10}O_8P^-$) and the neutral loss of the inositol moiety (162.053 Da, $C_6H_{10}O_5$) were used to characterize the lipid class, and the neutral loss of 254.226 Da (Acyl 16:0;O) was used to characterize the acyl chain moiety in ESI(−)-MS/MS. Alternatively, only the lipid class was characterized by the neutral loss of $C_6H_{13}O_9P$ (260.03 Da, phosphoinositol moiety) in ESI(+)-MS/MS. The ceramide phosphoethanolamine[56] (PE-Cer, PE-Cer 36:1;2O, PE-Cer 18:1;2O/18:0, level 2) incorporating sphingosine and fatty acid was characterized only in negative ion mode in this study. For PE-Cer 18:1;2O/18:0, the product ions of *m/z* 78.959 ($PO_3^-$) and *m/z* 140.012 ($C_2H_7NO_4P^-$) were used for the lipid class characterization, and the Acyl 18:0 loss (266.261 Da) was used for the acyl chain characterization. The ceramide phosphoethanolamine[56] (PE-Cer+O, PE-Cer 35:0;3O, PE-Cer 18:0;2O/17:0;O, level 2) incorporating sphingosine and hydroxy fatty acid was also characterized in negative ion mode by the same scheme as described in PE-Cer. The ceramide 1-phosphate (CerP, CerP 30:1;2O, CerP 18:1;2O/12:0, level 1) incorporating sphingosine and fatty acid moieties was quantified as $[M+H]^+$ and was characterized in both ion modes. For CerP 18:1;2O/12:0, the product ion of *m/z* 264.269 (SPB 18:1;2O-2$H_2O$) and the neutral loss of $H_3PO_4$ (97.977 Da) were used to characterize the acyl chain and lipid class characterizations, respectively. In negative ion mode, the product ions of *m/z* 78.959 ($PO_3^-$) and *m/z* 96.969 ($H_2PO_4^-$) were used to characterize the lipid class and the neutral loss of Acyl 12:0 (182.167 Da) was utilized for the acyl chain characterization.

For the sulfatide (SHexCer, SHexCer 36:1;2O, SHexCer 18:1;2O/18:0, level 1) incorporating sphingosine and fatty acid, the ion mode for quantifications depended on the injection volume and biological sample analyzed although the acyl chain compositions could only be determined in positive ion mode. Basically, the sensitivity of SHexCer was higher in negative ion mode and the quantification was performed by negative ion mode whereas the annotation was carried out by positive ion mode. The SHexCer lipids could be characterized as $[M+H]^+$ and $[M-H]^-$. For SHexCer 18:1;2O/18:0, the neutral loss of 260.02 Da ($H_2SO_4$+$C_6H_{10}O_5$) was used to characterize the lipid class and the product ions of *m/z* 282.279 (SPB 18:1;2O-$H_2O$, $C_{18}H_{36}NO^+$), *m/z* 264.269 (SPB 18:1;2O-2$H_2O$, $C_{18}H_{34}N^+$), and *m/z* 252.269 (SPB 18:1;2O-$CH_4O_2$, $C_{17}H_{34}N^+$) were used to characterize the acyl chain moiety in positive ion mode. Alternatively, the product ion of *m/z* 96.96 ($HSO_4^-$) was used for the lipid class characterization in positive ion mode. The sulfatide (SHexCer+O[49], SHexCer 36:1;3O, SHexCer 18:1;2O/18:0;O, level 2) incorporating sphingosine and hydroxy fatty acid was quantified and characterized using the same scheme as described for SHexCer. The monosialodihexosylganglioside (GM3, GM3 36:1;2O, GM3 18:1;2O/18:0, level 1) incorporating sphingosine and fatty acid was quantified as $[M-H]^-$ whereas the acyl chain composition could only be characterized by the $[M+NH_4]^+$ form. In positive ion mode, the product ion of *m/z* 292.103 ($C_{11}H_{18}NO_8^+$, *N*-acetyl-neuraminidate moiety) and the sequential neutral losses of 291.096

Da ($C_{11}H_{17}NO_8$, *N*-acetyl-neuraminidate moiety), 162.053 Da ($C_6H_{10}O_5$, galactosyl moiety), and 162.053 Da ($C_6H_{10}O_5$, glucosyl moiety) were used for the lipid class characterization. For GM3 18:1;2O/18:0, the product ions of *m/z* 282.279 (SPB 18:1;2O-$H_2O$, $C_{18}H_{36}NO^+$) and *m/z* 264.269 (SPB 18:1;2O-2$H_2O$, $C_{18}H_{34}N^+$) were used for characterizing the sphingobase moiety. In negative ion mode, the product ion of *m/z* 290.088 ($C_{11}H_{16}NO_8^-$, *N*-acetyl-neuraminidate moiety) and the neutral loss of 291.096 Da ($C_{11}H_{17}NO_8$, *N*-acetyl-neuraminidate moiety) were used to characterize the lipid class in negative ion mode. The sulfonolipid[57] (SL, SL 33:0;O, SL 18:0;O/15:0, level 2) incorporating deoxysphinganine/deoxysphingosine and fatty acid was quantified as $[M+H]^+$ and it could be characterized as $[M+H]^+$, $[M+NH_4]^+$, and $[M-H]^-$. For the ammonium adduct form of SL 18:0;O/15:0, the neutral losses of 241.241 Da (Acyl 15:0+$NH_3$), 259.251 Da (241.241 Da+$H_2O$), 255.256 Da (SPB 18:0;O+$NH_3$-$C_2H_8N$), and 273.267 Da (255.256 Da+$H_2O$) were used to characterize the acyl chain moieties whereas the product ion of *m/z* 124.006 ($C_2H_6NO_3S^+$) was used for the lipid class characterization. For the proton loss form of SL 18:0;O/16:0 in negative ion mode, the neutral loss of Acyl 16:0 (238.229 Da) and the product ion of 79.957 ($HSO_3^-$) were used to characterize the acyl chain moiety and the lipid class, respectively. The sulfonolipid[57] (SL+O, SL 33:1;2O, SL 17:0;O/16:1;O, level 2) incorporating deoxysphinganine/deoxysphingosine and hydroxy fatty acid was quantified as [M+H]+, and it could be characterized as $[M+H]^+$, $[M+NH_4]^+$, and $[M-H]^-$. For the proton adduct form of SL 17:0;O/16:1h in positive ion mode, the product ion of *m/z* 253.216 ($C_{16}H_{29}O_2^+$, Acyl 16:1;$O^+$) and the neutral loss of 270.22 Da (Acyl 16:1;O+$H_2O$) were used to characterize the acyl chains with the same product ion of *m/z* 124.006 being used for the lipid class annotation. The same diagnostic ions were used in negative ion mode as described for the normal SL lipid.

**Annotations in LipidMAPS Sterol Lipids [ST]: Supplementary Fig. 6.** The annotation of cholesterol (level 1) should be performed by checking the retention time of the internal standard, and it was quantified as $[M-H_2O+H]^+$ because of the highest sensitivity compared to other adduct forms whereas it could also be detected as $[M+H]^+$, $[M+NH_4]^+$, and $[M+Na]^+$. Although MS-DIAL 4 exports the annotation as "cholesterol" as default, the terminology should be changed to ST 27:1;O as recommended in LSI where ST and O denote the sterol backbone and oxygen-atom added to the normal ST moiety, respectively, if no standard is available. For other sterols, the structural backbone for brassicasterol, campesterol, sitosterol, and stigmasterol were described as ST 28:2;O, ST 28:1;O, ST 29:1;O, and ST 29:2;O, respectively. The cholesteryl ester (CE, level 1) was quantified as $[M+NH_4]^+$ and characterized by the product ion of *m/z* 369.349 (ST $27:1^+$, $C_{27}H_{45}^+$). The other sterol esters[58] including brassicasteryl ester (BRSE, level 2), campesteryl ester (CASE, level 2), sitosteryl ester (SISE, level 2), and stigmasteryl ester (STSE, level 2) were quantified as $[M+NH_4]^+$ and characterized by the product ions of *m/z* 381.341 (ST $28:2^+$, $C_{28}H_{45}^+$), *m/z* 383.376 (ST $28:1^+$, $C_{28}H_{47}^+$), *m/z* 397.386 (ST $29:1^+$, $C_{29}H_{49}^+$), and *m/z* 395.363 (ST $29:2^+$, $C_{29}H_{47}^+$), respectively, where the annotations were based on the species levels of SE 28:2/18:1, SE 28:1/18:1, SE 29:1/18:1, and SE 29:2/18:1, respectively, for oleate ester description.

The sulfate conjugate[59] to sterols (SSulfate, level 2) were characterized as $[M-H]^-$ by the product ion of *m/z* 96.96 ($HSO_4^-$) and annotated as ST 27:1;O;S for cholesterol sulfate. The hexosyl conjugate, known as sterolglycosides[58] (SHex, level 2), were quantified as $[M+NH_4]^+$ and could also be characterized as the $[M-H]^-$ form. For example, the stigmasterol hexoside was annotated as ST 29:2;O;Hex with the confirmation of the product ions of *m/z* 395.367 (ST $29:2^+$, $C_{29}H_{47}^+$) and *m/z* 179.0561 (hexose moiety, $C_6H_{11}O_6^-$) in positive and negative ion mode, respectively. The acylhexosyl sterols[60], also known as acylsterolglycosides, including acylhexosyl- cholesterol (AHexCS, level 2), brassicasterol (AHexBRS, level 2), campesterol (AHexCAS, level 2), sitosterol (AHexSIS, level 2), and stigmasterol (AHexSTS, level 2) were quantified as $[M+NH_4]^+$ and also characterized as $[M-H]^-$ and $[M+CH_3COO]^-$, where the annotations were performed according to the species levels of ST 27:1;O;Hex;FA 18:1, ST 28:2;O;Hex;FA 18:1, ST 28:1;O;Hex;FA 18:1, ST 29:1;O;Hex;FA 18:1, and ST 29:2;O;Hex;FA 18:1, respectively, for oleate ester description. The characterizations for AHexCS, AHexBRS, AHexCAS, AHexSIS, and AHexSTS were performed in positive ion mode by the product ions of *m/z* 369.349 (ST $27:1^+$, $C_{27}H_{45}^+$), *m/z* 381.341 (ST $28:2^+$, $C_{28}H_{45}^+$), *m/z* 383.376 (ST $28:1^+$, $C_{28}H_{47}^+$), *m/z* 397.386 (ST $29:1^+$, $C_{29}H_{49}^+$), and *m/z* 395.363 (ST $29:2^+$, $C_{29}H_{47}^+$), respectively, whereas the FA ions derived from the ester acyl chain moiety were used for the lipid annotations in negative ion mode.

The annotation of bile acids (level 1) was ideally performed by matching the retention time and MS/MS spectrum of authentic standards. Otherwise, the annotations for e.g. cholic acid, glycodeoxycholic acid, and taurodeoxycholic acid were described as ST 24:1;O5, ST 24:1;O4;G, and ST 24:1;O4;T, respectively. The quantification was performed by the proton loss form in negative ion mode. The sulfate conjugate[59] (BASulfate, level 2) was quantified and characterized as $[M-H]^-$ by the product ion of *m/z* 96.96 ($HSO_4^-$) and was annotated as ST 24:1;O5;S for cholic acid sulfate. Bile acid esters such as deoxycholic acid ester (DCAE, level 3) were quantified as $[M+NH_4]^+$ and also characterized by $[M-H]^-$, where the annotation was performed according to the species level as ST 24:1;O4/18:1 for oleate ester description. For DCAE lipid, the product ion of *m/z* 357.274 ($C_{24}H_{37}O_2^+$, ST 24:1;$O2^+$) and the FA ion derived from the ester chain moiety were used for the lipid characterizations in positive- and negative ion mode, respectively. Only one lipid, calcidiol, for Vitamin D (level 1) was characterized as $[M+H]^+$ in this study by the sequential $H_2O$ losses from the precursor *m/z* value.

**Annotations in LipidMAPS Prenol Lipids [PR]: Supplementary Fig. 7.** Three phenol lipids including coenzyme Q (CoQ, level 1), vitamin A fatty acid ester (VAE, level 1), and vitamin E (tocopherol, level 1) were incorporated in MS-DIAL 4. The CoQ lipid was annotated as $[M+H]^+$ by the product ion of *m/z* 197.081 ($C_{10}H_{13}O_4^+$): CoQ8, 9, and 10 were characterized in this study. The VAE lipid was characterized as $[M+H]^+$ and $[M+Na]^+$, and the sodium adduct form was utilized for quantification because of higher sensitivity than that of the proton adduct form in our experimental condition. The product ions of *m/z* 269.226 ($C_{20}H_{29}^+$) and *m/z* 119.086 ($C_9H_{11}^+$) were used for the diagnostic ions. The tocopherol lipid was characterized as $[M-H]^-$ and $[M+CH_3COO]^-$ and quantified by the proton loss form. The product ion of *m/z* 163.075 ($C_{10}H_{11}O_2^-$) was used for the characteristic ion.

### Reference (Supplementary Note 2)
